# Differential Impact of Multiple Sensory Deprivation on Spatially Modulated Cells in Medial Entorhinal Cortex

**DOI:** 10.1101/2024.09.09.611946

**Authors:** Jiayi Tian, Shidan Wen, Yuqian Zhou, Nan Hu, Shuyang Yao, Xuan Zhang, Yuanjing Liu, Yuheng He, Zhouyuan Wang, Chenglin Miao

**Author notes:** Correspondence: Chenglin Miao.

## Abstract

Spatial navigation depends on anchoring internal spatial maps to external environments, guided by sensory cues such as visual and tactile information. The Medial Entorhinal Cortex (MEC) is crucial for integrating these sensory inputs during the formation of spatial maps. While the responsiveness of many spatially modulated cells to visual stimuli is well-established, the role of tactile sensation in spatial representation is less understood. Rodents primarily gather tactile information through their whiskers, which provide essential spatial and textural details via whisking movements, potentially vital for constructing accurate spatial maps. In our study, we employed miniature two-photon microscopy to monitor neural activity in the MEC of freely moving male mice subjected to visual and tactile deprivation. We found that after spatial maps were established under visual guidance, head direction and border cells were disrupted by both light deprivation and whisker trimming, whereas grid and spatial cells were primarily dependent on visual input. In complete darkness with enriched tactile cues, however, all spatially modulated cell types could anchor their tuning to environmental tactile cues, and whisker trimming markedly impaired grid and spatial cell representations. A subset of neurons remained stable under combined deprivation, possibly relying on boundary-related tactile inputs from direct environmental contact. Additionally, we identified certain MEC neurons whose activity correlated with whisker movements, suggesting a potential role in integrating tactile information into spatial representations. These findings demonstrate that the MEC flexibly integrates multiple sensory inputs to sustain spatial coding and highlight a previously underappreciated contribution of tactile information to spatial navigation.

## Introduction

Spatial navigation depends on the reliable integration of environmental reference cues from various sensory inputs, including visual^1^, auditory^2,3^, olfactory^4^ and tactile stimuli^5^, into internal spatial maps. While path integration ^6–8^—computed from self-motion signals without the need for external references—forms a core component of spatial computation, external cues, particularly distal and proximal visual landmarks ^8,9^, are crucial for anchoring and calibrating internally generated estimates to improve navigational accuracy. The medial entorhinal cortex (MEC) plays a key role in this process by combining self-motion cues with external sensory information such as environmental boundaries and landmarks ^10–12^, thereby providing the hippocampus with a coherent framework for spatial cognition and memory.

The grid cell, initially characterized by Hafting et al. in 2005, constitutes the first identified spatially modulated cell type within the MEC^13^, which exhibit action potentials at locations forming a hexagonal grid pattern. Alongside grid cells, the MEC encompasses other functionally distinct cell types, including head-direction (HD) cells^14^, border cells^15,16^, non-grid spatial cells^17^ and speed cells^18,19^. It is hypothesized that grid cells play a dominant role in path integration, while HD cells and speed cells contribute to this process by likely providing self-motion-related information, as suggested by the continuous attractor model^12^. Additionally, border cells may help correct errors accumulated over time and distance in this process^11^. However, compared to grid cells, other spatially modulated cells in MEC sometimes exhibit different responses when environment changes^17,20,21^, which implies their respective sensitivity to external sensory input.

Although numerous studies have demonstrated how sensory inputs, particularly visual inputs, influence spatially modulated cells in MEC, such as HD cells^22,23^ and grid cells^21,24^, the specific role of tactile inputs in constructing a spatial cognitive map remains largely unexplored. Whiskers, also called facial vibrissae, are the primary tactile organs in rodents and their corresponding bilateral “whisking” behavior playing a crucial role in exploring environment^25^. Trimming animals’ whiskers results in impaired performance in spatial learning tasks^26^. And it has been discovered that the hippocampal CA1 region responds to whisker stimulation and is likely involved in the process of updating spatial cognitive maps^27^. Furthermore, the establishment of place fields in the hippocampus is modulated by tactile input, with different cells exhibiting varying response intensities^5^. These findings suggest that spatial map can be constructed using tactile information, encompassing behaviorally relevant spatial elements such as object positions^28^, specific boundaries^29,30^, and environmental surfaces^31,32^. A strong piece of evidence is that tactile information plays a crucial compensatory role in the absence of visual information when rats are required to complete a gap crossing task in darkness^30^. Beyond the vibrissae, cutaneous mechanoreceptors distributed across other body surfaces—such as the glabrous skin of the plantar paws and the rhinarium—are represented in the rodent somatosensory system and participate in active exploratory behaviors^33,34^. However, whether and how the spatially modulated cells in MEC respond to tactile inputs remain unclear.

In this study, we examined the alterations in spatial representation within the MEC following light deprivation and tactile deprivation using miniature two-photon microscopy. Our findings demonstrate that the MEC flexibly integrates available sensory inputs from the environment to construct spatial representations, with distinct functional cell types differentially modulated by visual and tactile cues. Border cells rely heavily on tactile perception and are consistently and strongly affected by whisker trimming regardless of visual condition; whereas HD cells, grid cells, and spatial cells are more profoundly disrupted by whisker trimming specifically when visual cues are absent and animals must rely predominantly on tactile information to maintain spatial representations. When both whisker trimming and removal of environmental walls are combined in darkness, spatial representations across all cell types are severely degraded—most strikingly, the fields of border cells are nearly completely abolished. Furthermore, we identified a significant correlation between calcium signals in a subset of MEC neurons and whisking movements, suggesting that these neurons may possess the capacity to process tactile information for the purposes of spatial navigation.

## Results

### Head-direction tuning in the MEC is disrupted by darkness and whisker trimming

To investigate the influence of visual and tactile inputs on spatially modulated cells in the MEC, recombinant adeno-associated virus (rAAV) was used to express the Ca2+ indicators GCaMP6 in the MEC of the animal (Figure S1). Using an autoregressive (AR) model-based algorithm to estimate the time constant, we found that the two calcium indicators, GCaMP6s and GCaMP6f, did not exhibit significant differences in calcium signal extraction performance in our data (Figures S2A–C). A prism was implanted between the cortex and the cerebellum, enabling the acquisition of calcium signals via a miniature two-photon microscope^35^ when mice are freely exploring the open-field (Figures 1A–D). Initially, the mice were allowed to freely explore an open field for approximately 30 minutes under illuminated conditions, establishing a stable baseline representation (Light, L). To assess the impact of visual deprivation, the lights were turned off and the mice were allowed to explore the same environment in complete darkness (Dark, D). Subsequently, the mice underwent whisker trimming and were reintroduced to the environment, now deprived of both visual and tactile inputs (Dark/Whisker-trimmed, DW) (Figure 1A). This design allowed us to evaluate the individual effects of light deprivation as well as the combined impact of darkness and whisker trimming, based on the premise that animals tend to rely on alternative sensory cues to support spatial cognition when a specific sensory modality is compromised^36,37^. A cohort of 13 male mice was used in this paradigm, and multiple types of spatially modulated neurons were identified within each field of view (FOV) (Figures S2D–E). All sessions were conducted in an open-field arena equipped with three distinct visual cue cards mounted on the surrounding walls (Environment 1).

**Figure 1.**
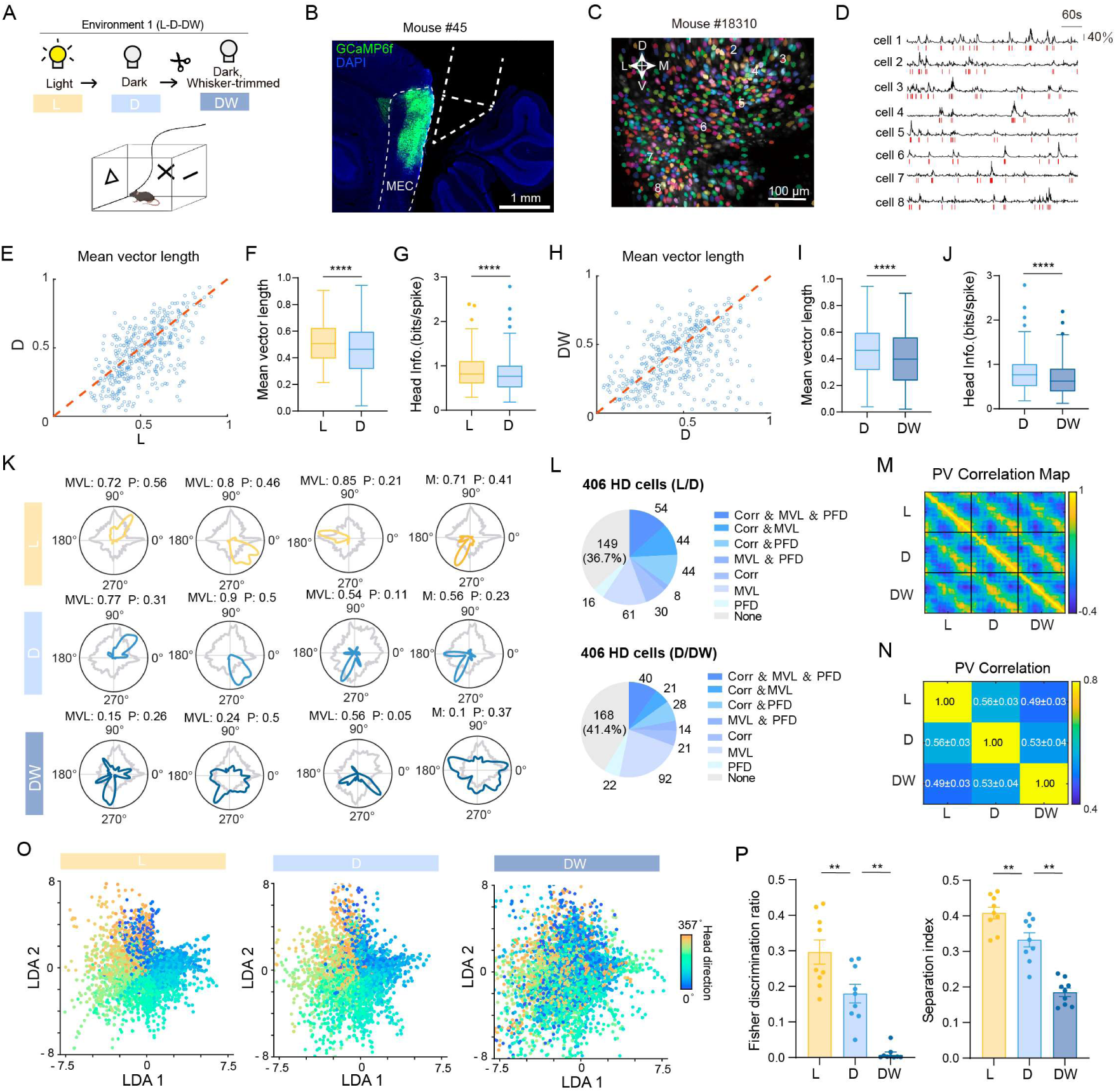
Two-photon imaging reveals degradation of head-direction tuning after light deprivation and whisker trimming. (A) Schematic of the experimental design. Mice explored an open-field arena in light (Environment 1, E1) with three visual cue cards on the arena walls, followed by recording in complete darkness (D) and then darkness with whisker trimming (DW) during miniaturized two-photon imaging of MEC activity. (B) Representative histological image showing GCaMP6f expression in the MEC with a prism inserted at the cerebellar interface. Scale bar, 1 mm. (C) Representative field of view (FOV). Left, maximum projection image showing the spatial distribution of extracted regions of interest (ROIs), with each ROI displayed in a distinct color. A directional compass indicates the dorsal (D)-ventral (V) and medial (M)-lateral (L) orientation within the imaging plane. Scale bar, 100 μm. (D) Deconvolved calcium activity traces from the eight example cells in Fig.C over a 60-second period. Detected calcium events are marked with red ticks. Vertical scale bar, 40% ΔF/F. (E) Scatter plot showing the distribution of MVL of all HD cells (n = 406) in Light (L) and Dark (D) conditions. (F) Box plot showing a significant reduction in MVL from L to D (Wilcoxon signed-rank test, n = 406, P = 2.5 × 10^-10^). (G) Box plot showing a significant decrease in head information (bits/spike) from L to D (Wilcoxon signed-rank test, n = 406, P = 1.1 × 10^-5^). (H–J) Similar analyses as in (E–G), showing changes in MVL and head information from Dark (D) to Dark, Whisker-trimmed (DW) (Wilcoxon signed-rank test, n = 406, MVL: P = 1.2 × 10^-6^, Head Info.: P = 2.9 × 10^-9^). (K) Representative HD tuning curves from example neurons across L, D, and DW sessions. MVL and peak firing rate (P) are indicated for each cell. (L) Pie charts summarizing the proportions of HD cells showing significant tuning changes relative to the corresponding baseline session (L for L/D; D for D/DW), quantified using three metrics: tuning-curve correlation (Corr), mean vector length (MVL), and preferred firing direction (PFD). Cells were classified as exhibiting a significant decrease in Corr, a significant decrease in MVL, and/or a significant shift in PFD, either individually or in combination, as indicated in the legend. (M) Population vector correlation of all HD cells across sessions, with 120 angular bins per session. Color represents correlation values, with yellow indicating the highest and blue the lowest. (N) Correlation between population vector matrices of all HD cells within individual mouse between each session pair, shown as mean ± SEM, n = 13 mice. (O) LDA projections of HD cell activity in each session along axes of maximal head direction separability, with colors representing different head direction angles. (P) Fisher discrimination ratio and Separation index across sessions for all mice, showing significant decreases after light deprivation (L/D) and whisker trimming (D/DW) (Wilcoxon signed-rank test, n = 9 mice with more than 20 HD cells, Fisher discrimination ratio: L/D: 0.0039, D/DW: P = 0.0039; Separation index: L/D: P = 0.0039, D/DW: P = 0.0039). ***P ≤ 0.001; **P ≤ 0.01; *P ≤ 0.05; ns, P > 0.05. Box plots are presented in Tukey style: the horizontal line within each box represents the median; the bottom and top edges indicate the 25th and 75th percentiles (Q1 and Q3), respectively. Whiskers extend to the most extreme data points within 1.5 times the interquartile range (IQR) from the quartiles. Data points beyond this range are shown individually as outliers. All statistical tests were two-tailed unless otherwise stated. Unless otherwise stated, error bars in all bar graphs represent the standard error of the mean (SEM).

First, we investigated the influence of visual and tactile inputs on head-direction tuning in the MEC. Calcium traces were extracted from 3,758 neurons, of which 406 (10.8%) were identified as HD cells from baseline session and used for further analysis. HD cells were identified by comparing each neuron’s real mean vector length (MVL) to a shuffle distribution generated by 1,000 circular time shifts of its calcium trace. A neuron was classified as an HD cell only if its real MVL exceeded the 99^th^ percentile of its shuffle distribution and its within-session tuning stability was greater than 0.5. In the baseline light session, the mean MVL of these HD cells was 0.52 (Figure S3A), with their preferred firing directions (PFD) and the mice’s head orientations being nearly uniformly distributed between 0 and 360° (Figures S3B–C). We subsequently calculated the tuning curves of selected HD cells for each experimental session and performed statistical analysis using Wilcoxon signed-rank test to compare the tuning abilities of the HD cell population. Our analysis revealed that both the MVL and head information (Head Info.) were significantly reduced in the dark environment compared to the baseline light condition (Figures 1E–G; L/D; Wilcoxon signed-rank test, n = 406, MVL: P = 2.5 × 10^-10^; Head Info.: P = 1.1 × 10^-5^). Moreover, the subsequent whisker trimming procedure further disrupted HD tuning in the MEC. (Figure 1H–J; D/DW; MVL: P = 1.2× 10^-6^; Head Info.: P = 2.9 × 10^-9^). Furthermore, we applied a more stringent criterion by using the 99^th^ percentile of the shuffled MVL distribution pooled across all mice (0.42; Figure S3A) and then re-evaluated the originally identified HD cells whose MVL exceeded this threshold. This analysis yielded identical statistical outcomes, further supporting the robustness of our conclusions (L/D: Wilcoxon signed-rank test, n = 279, MVL: P = 6.7 × 10^-9^; Head Info.: P = 1.1× 10^-6^; D/DW: MVL: P = 3.4 × 10^-5^; Head Info.: P = 6.1 × 10^-7^).

Individual HD cells exhibited heterogeneous responses to darkness or whisker trimming, with tuning strength and MVL showing varying degrees of change following sensory deprivation (Figure 1K). To systematically assess the impact of sensory deprivation on HD cell tuning properties, we developed a subsampling-based approach to quantify whether individual HD cells exhibited statistically significant changes across three metrics: MVL, PFD shift, and tuning curve correlation. For each pairwise session comparison, equal-duration data segments were randomly drawn from each session and this procedure was repeated 100 times; cells were classified as significantly affected if they showed a significant decline in MVL, a significant shift in PFD, or a significant reduction in cross-session tuning curve correlation relative to within-session stability (see Methods). Following the transition from light baseline to darkness, 167 of 406 (41.1%) HD cells exhibited a significant MVL decline, 122 of 406 (30.0%) cells showed a significant PFD shift, and 172 of 406 (42.4%) cells displayed a significant reduction in tuning curve correlation. When considering all three criteria jointly, 54 of 406 (13.3%) cells were significantly affected across all three metrics, while 149 of 406 (36.7%) remained unaffected by visual deprivation on any metric (Figure 1L, upper). Following subsequent whisker trimming in darkness, 167 of 406 (41.1%) HD cells showed a significant MVL decline, 104 of 406 (26.1%) exhibited a significant PFD shift, and 110 of 406 (27.1%) showed a significant reduction in tuning curve correlation; 168 of 406 (41.4%) cells remained unaffected across all metrics (Figure 1L, lower). Cells that remained unchanged across all three metrics were defined as resistant to a given form of sensory deprivation. The proportion of resistant cells was comparable between the darkness and whisker-trimming conditions (36.7% vs. 41.4%; χ² = 1.9, P = 0.17), indicating that the two manipulations produced similar fractions of unaffected HD cells.

Also, we computed the population vector (PV) correlation for all HD cells between each session pair. The results showed that both darkness and whisker-trimming led to destabilization of population-level HD coding compared with its previous state (Figures 1M–N; n = 13, mean± SEM: L/D: 0.56 ± 0.03; D/DW: 0.49 ± 0.03). To further examine changes in HD encoding ability, we applied Linear Discriminant Analysis (LDA) to reduce the dimensionality of neural activity recorded from the HD cell population. In the light condition, neural representations of different head direction angles were well-separated, demonstrating accurate HD tuning under normal sensory conditions (Figure 1O and Figure S3F). This representation remained stable across time within the baseline session, as evidenced by the consistent separation of head directions when projecting neural activity from the first and second halves of the session separately (Figure S3D). In the dark condition, the separation between head directions was notably reduced, suggesting a drift in population-level HD representation away from that established under illuminated conditions, likely due to the absence of visual cues. This drift was further amplified following whisker trimming, where the encoding of head direction was almost entirely abolished in some mice (Figure 1O and Figure S3F). Quantitative analysis further confirmed these effects, since both the Fisher’s discrimination ratio and the separation index exhibit a significant decline from the light condition to the dark and further to the dark with whisker trimming conditions (Figure 1P; Wilcoxon signed-rank test, n = 9, Fisher’s discrimination ratio: L/D:P = 0.0039, D/DW:P = 0.0039; separation index: L/D: P = 0.0039, D/DW: P = 0.0039). These metrics indicate a progressive drift in the original population-level HD representation, with the most pronounced impairment occurring under combined visual and tactile deprivation. Nevertheless, when we applied LDA to the DW session and used it as the reference for dimensionality reduction, we found that the HD neural population still retained some capacity to distinguish head direction angles (Figure S3E), although the discriminability was reduced compared the baseline light session. Thus, while sensory deprivation reduced the fidelity of HD tuning, the directional signal was not completely abolished.

Following the DW session, we reintroduced visual input by turning the lights back on and allowed the animals to freely re-explore the environment (Light/Whisker-trimmed, LW; Figure S3G). However, the population-level HD tuning strength did not recover (Figure S3H), indicating that the effects of whisker trimming persisted despite restoration of visual cues. At the single-cell level, HD responses exhibited substantial heterogeneity. In some cells, MVL increased and a clear PFD re-emerged (Figure S3I), whereas others failed to recover, potentially reflecting differential dependence on tactile inputs. When LDA was applied again to evaluate population-level encoding of head direction, considerable variability was observed across mice (Figure S3J). In some animals, the drifted HD representation did not recover during the subsequent LW session (Figures S3J), indicating that reintroduction of visual cues alone was insufficient to restore the original HD representation. However, when LDA was trained and projected using data from the LW session itself, the population activity appeared to encode head direction more effectively than when baseline light-session activity was used as the reference (Figures S3J). This finding suggests that, following reintroduction of visual input, the HD network may have stabilized to a new, internally consistent reference frame rather than returning to the original alignment.

In addition, the accuracy of DLC in tracking animals’ position and the quality of the imaging data were consistent across all sessions, thereby ruling out the possibility that the observed changes in cells’ firing patterns were due to these factors (Figures S4A–C). Behavioral performance also remained largely unaffected following whisker trimming (Figure S4D). Animals maintained extensive spatial coverage of the arena throughout all sessions (Figure S4E), and no significant differences were observed in running speed, time spent near the boundaries, or wall-directed exploratory behavior after whisker trimming (Figures S4F–H). Notably, however, whisker trimming resulted in a significant reduction in head angular velocity and angular acceleration (Figures S4I–J). Given that whiskers contribute to head orientation, their removal may influence head movement dynamics in addition to reducing tactile input. Therefore, we cannot exclude the possibility that changes in angular head-velocity-related signals contributed to the disruption of HD tuning, as vestibular/AHV inputs are known to influence HD circuits^38,39^.

Importantly, we subjected another three mice to the same experimental paradigm but without whisker trimming, to serve as control experiments (Figures S5A–B). The results showed that, the MVL and head information of HD cells decreased after light deprivation. However, no further changes were observed during a subsequent dark session. These findings suggest that the deterioration in HD tuning after whisker-trimming was not due to a progressive drift in cellular representations during darkness, but rather caused by the deprivation of tactile cues provided by the whiskers. Notably, similar patterns were observed in other types of spatially modulated cells (Figures S5).

### Border cells in MEC are modulated by both visual and tactile inputs

In addition to HD cells, MEC contains various types of spatially modulated cells, including border cells, grid cells and non-grid spatial cells. We next sought to determine whether border cells in MEC are influenced by visual inputs and whisker-mediated tactile inputs. A total of 223 neurons (5.9%) were identified as border cells (see Methods), with an average border score of 0.63 (Figure S6A).

Our analysis revealed that the maintenance of boundary fields is supported by both visual and tactile inputs. Specifically, both border score and spatial information significantly decreased in the dark condition compared to light condition (Figure 2A–C; L/D; Wilcoxon signed-rank test, n = 223, Border score: P = 5.2 × 10^⁻24^; Spatial Info.: P = 0.0085). Furthermore, whisker trimming in darkness led to an additional decrease in both characters (Figure 2D–F; D/DW; Wilcoxon signed-rank test, n = 223, Border score: P = 0.026; Spatial Info.: P = 0.00015). Following disruption of sensory input, border cells exhibited markedly increased firing in non-border regions, and a subset lost border-selective tuning entirely after whisker trimming (Figure 2G). To quantify the extent to which visual and tactile deprivation impaired border cels, we applied an approach analogous to that used for HD cells, assessing three complementary metrics: the border score, the spatial precision of border firing fields (measured as the distance from the peak firing location to the nearest wall), and the stability of the spatial firing pattern across sessions (see Methods). Under dark conditions, 131 of 223 border cells (58.7%) showed a significant reduction in border score relative to the baseline session; 56 of 223 cells (25.1%) exhibited a significant increase in peak firing distance to the nearest wall; and 85 of 223 cells (38.1%) showed a significant reduction in cross-session rate map correlation (L/D) relative to their within-session correlation of baseline session (L/D). When each border cell was classified into one of eight mutually exclusive categories—defined by whether none, one, two, or all three metrics were significantly altered—only 27.4% of border cells were entirely unaffected by darkness. Following additional whisker trimming under dark conditions, 100 of 223 border cells (44.8%) showed a significant reduction; 67 of 223 cells (30.0%) displayed a significant increase in peak-to-wall distance; and 83 of 223 cells (37.2%) showed a significant reduction in cross-session rate map correlation (D/DW) relative to within-session correlation of dark session (D/D). Only 34.5% of border cells remained completely unaffected by whisker trimming. No significant difference was found in the proportion of border cells unaffected by darkness versus whisker trimming (Chi-square test, χ² = 1.6, *P* = 0.10).

**Figure 2.**
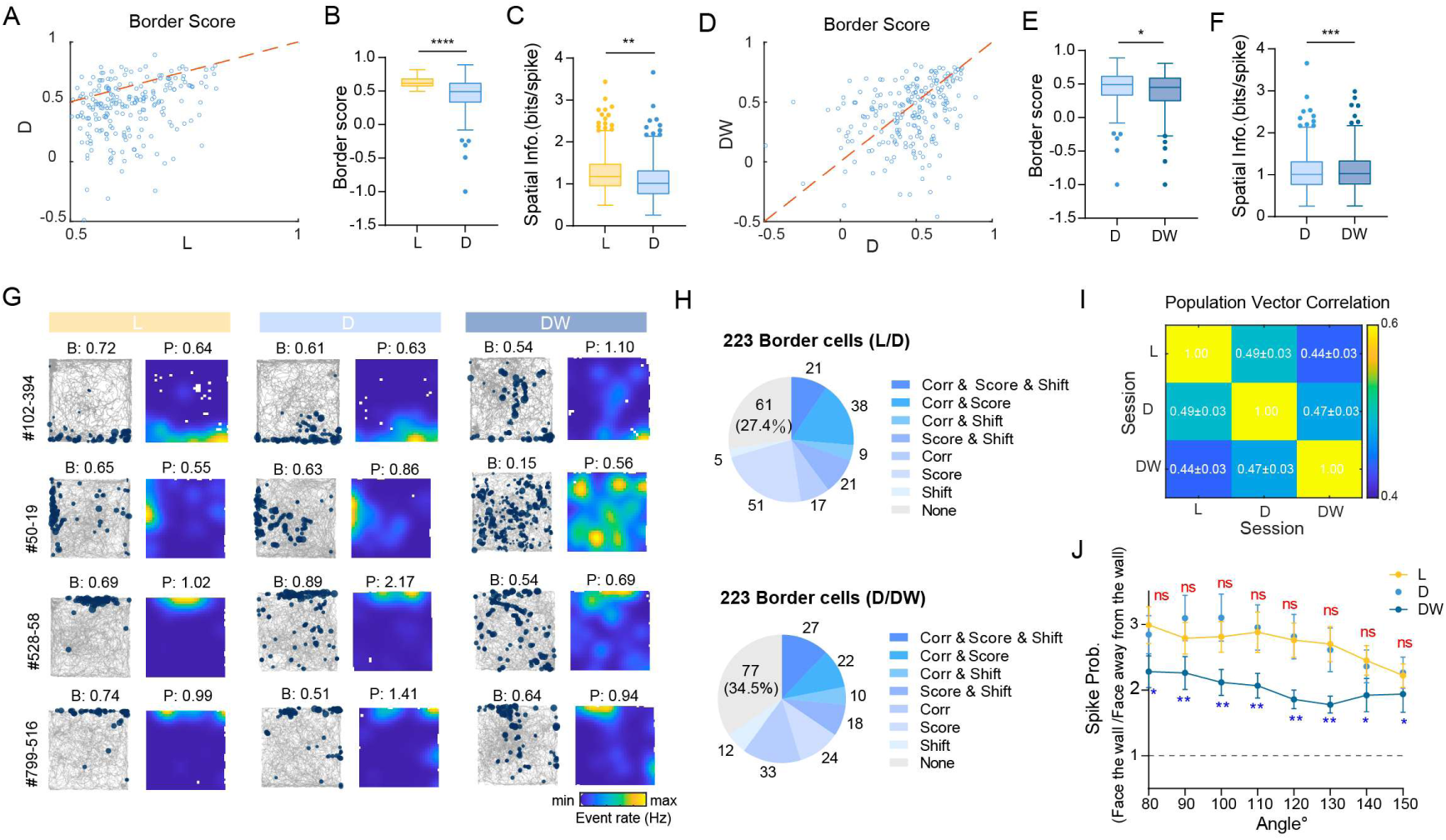
MEC border cells are jointly modulated by visual inputs and whisker-supported tactile inputs. (A) Scatter plot illustrating distribution of border score for all border cells (n = 223) in Light (L) and Dark (D) conditions. (B) Box plot showing a significant reduction in border score from L to D (Wilcoxon signed-rank test, n = 223, P = 5.2 × 10^-24^). (C) Box plot demonstrating a significant decrease in spatial information (bits/spike) from L to D (Wilcoxon signed-rank test, n = 223, P = 0.0085). (D–F) Similar analyses as shown in (A–C), depicting changes in border scores and spatial information after whisker trimming in darkness (D/DW) (Wilcoxon signed-rank test, n = 223, border score: P = 0.026; spatial information: P = 0.00015). (G) Representative rate maps of border cells across sessions (L, D, DW). For each cell, the animal’s trajectory (gray) and the corresponding spatial rate map (color-coded by event rate) are shown; B and P indicate border score and peak rate, respectively. (H) Pie chart summarizing the proportions of border cells exhibiting significant changes in multiple tuning metrics between L/D and D/DW: a significant decrease in rate map correlation (Corr), a significant decrease in border score (Score), and/or an increase in the distance of the peak firing field from the nearest boundary (Shift), either individually or in combination (legend). (I) Correlation of population vector matrices of border cell activity across sessions (L, D, DW), averaged across mice (mean ± SEM, n = 13 mice). (J) Wall-facing modulation of border-cell spiking quantified as the ratio of spike probability when facing the wall to that when facing away from the wall, computed within boundary regions and evaluated across angle limits (see Methods). For each session (L, D, DW), the ratio was tested against 1 using a one-sample Wilcoxon signed-rank test at each angle limit (ratio > 1 indicates stronger firing when facing the wall). Red “ns” denotes the result of paired between-session comparisons between L and D at each angle limit; blue asterisks denote comparisons between D and DW at each angle limit (Wilcoxon signed-rank tests) ****P ≤ 0.0001; ***P ≤ 0.001; **P ≤ 0.01; *P ≤ 0.05; ns, P > 0.05.

The disrupted firing pattern of the border cell population was not observed in the control experiments (Figures S5A and S5C, Wilcoxon signed-rank test, n = 67, P = 0.58). Moreover, this effect remained evident even when a more stringent selection criterion was applied, restricting the analysis to border cells with border scores exceeding the 95^th^ percentile of the population shuffle distribution (0.53)(L/D: Wilcoxon signed-rank test, n = 279, border score: P = 6.7 × 10^-23^; Spatial Info.: P = 0.0097; D/DW: border score: P = 0.026; Spatial Info.: P = 3.0 × 10^-5^). As some border cells (38 of 223, 17.0%) in our data also exhibited direction-tuning, we further investigate the performance of these pure border cells. We found that pure border cells likewise showed significant reductions in both border score and spatial information following light deprivation or whisker trimming (Figures S6B–C; Wilcoxon signed-rank test, n = 185, L/D: border score: P = 8.0 × 10^-21^, Spatial Info.: P = 0.003; D/DW: border score: P = 0.0046, Spatial Info.: P = 0.0059). The presence of border cells in our recordings was validated by the emergence of new firing fields in a subset of cells when a wall was inserted into the environment one day prior to the whisker-trimming experiments for some mice (Figure S6D). Cross-day comparisons further demonstrated that, among the border cells responsive to the newly inserted wall, some maintained stable firing properties across sensory conditions, while others were significantly affected by whisker trimming (Figure S6E). Border score and spatial information did not improve during the final LW session, indicating that the degraded spatial coding of border cells could not be restored after only visual inputs were reintroduced (Figure S6C; DW/LW; Wilcoxon signed-rank test, n = 223, Border score: P = 0.48; Spatial Info.: P = 0.98).

Unlike HD cells and grid cells, border cells have not been shown to possess an attractor network structure that could autonomously maintain stable representations. Consequently, border cells likely depend heavily on continuous sensory input to establish and sustain boundary representations—whether through direct visual observation of the enclosure walls or through physical contact mediated by the whiskers or snout. To directly examine the sensory basis of border cell activation, we analyzed firing patterns restricted to epochs when the animal occupied the border zone, computing for each cell the ratio of inward-facing to outward-facing event rate: the number of calcium events recorded during inward-facing epochs (headings directed toward the wall) divided by the corresponding dwell time, normalized by the rate during outward-facing epochs (Figure S6F). A ratio exceeding 1 indicates that the cell fires at a higher rate when the animal faces toward the wall than away from it. Border cells showed a significantly elevated inward-to-outward event rate ratio under baseline conditions (Figure S6G). In darkness, this ratio was not significantly altered; however, subsequent whisker trimming produced a significant reduction, implicating whisker-wall contact as a meaningful contributor to border cell activation (Figure S6G). Nevertheless, the directional firing preference was not entirely abolished even under combined sensory deprivation, suggesting that additional sensory inputs—such as direct physical contact of the snout or body with the wall—may also contribute to border cell activation at environmental boundaries.

### Visual inputs predominate over tactile inputs in maintaining grid cell spatial coding when tactile cues are sparse

Grid cells are the predominant type of spatially modulated cells in MEC and have long been regarded as implicating in path integration^12,40^. Historically, grid cells were thought to operate independently of external sensory inputs, generating their characteristic hexagonal firing patterns based solely on self-motion information. However, recent studies have challenged this notion, demonstrating that spatial-coding performance of grid cells is modulated by various sensory inputs, especially by visual cues^10,21^.

In our study, we identified 215 grid cells (5.7% of all recorded neurons), with an average grid score of 0.65 (Figure S7A). We observed that the overall grid score significantly decreased in darkness, consistent with previous findings^21^ (Figures 3A–B; L/D; Wilcoxon signed-rank test, n =215, P = 2.5 × 10^-27^) and the spatial information exhibited no significant change, with only slight decreases in mean and median values (Figure 3C; mean ± SEM, L: 1.038 ± 0.03; D: 1.035 ± 0.03; Wilcoxon signed-rank test, n = 215, P = 0.78). However, the performance of grid cells did not further deteriorate following whisker trimming and the population-level grid score even slightly recovered (Figures 3D–F; Wilcoxon signed-rank test, n = 215, Grid score: P = 0.044; Spatial Info.: P = 0.061), suggesting a small influence of interactive effects of visual and tactile deprivation on grid cell function. Many grid cells lost their characteristic hexagonal firing patterns in darkness but showed no further deterioration following whisker trimming (Figure 3G). To further validate these findings, we applied a more stringent criterion for grid cell classification, restricting analyses to cells whose baseline grid scores exceeded the 95^th^ percentile of a shuffle distribution (0.41) constructed from the recorded population. The statistical results remained consistent: grid scores declined significantly under darkness but showed a slight increase following subsequent whisker trimming (L/D: Wilcoxon signed-rank test, n = 184, grid score: P = 1.3 × 10^-23^; Spatial Info.: P = 0.12; D/DW: grid score: P = 0.028; Spatial Info.: P = 0.066). Furthermore, given recent evidence that grid cells are anatomically clustered^41^, we selected four mice with enriched grid cells in the imaging FOV (>20 grid cells per animal) and performed statistical analyses exclusively on grid cells from these animals to minimize the potential confound of false-positive grid cell inclusion. Consistent with the full dataset, grid cells from these four mice showed substantial degradation of spatial coding under visual deprivation, with no significant further decline following whisker trimming (Figure S7B). Grid cells with pure spatial tuning and those with directional tuning properties also exhibited similar patterns of sensory dependence (Figures S7D–E).

**Figure 3.**
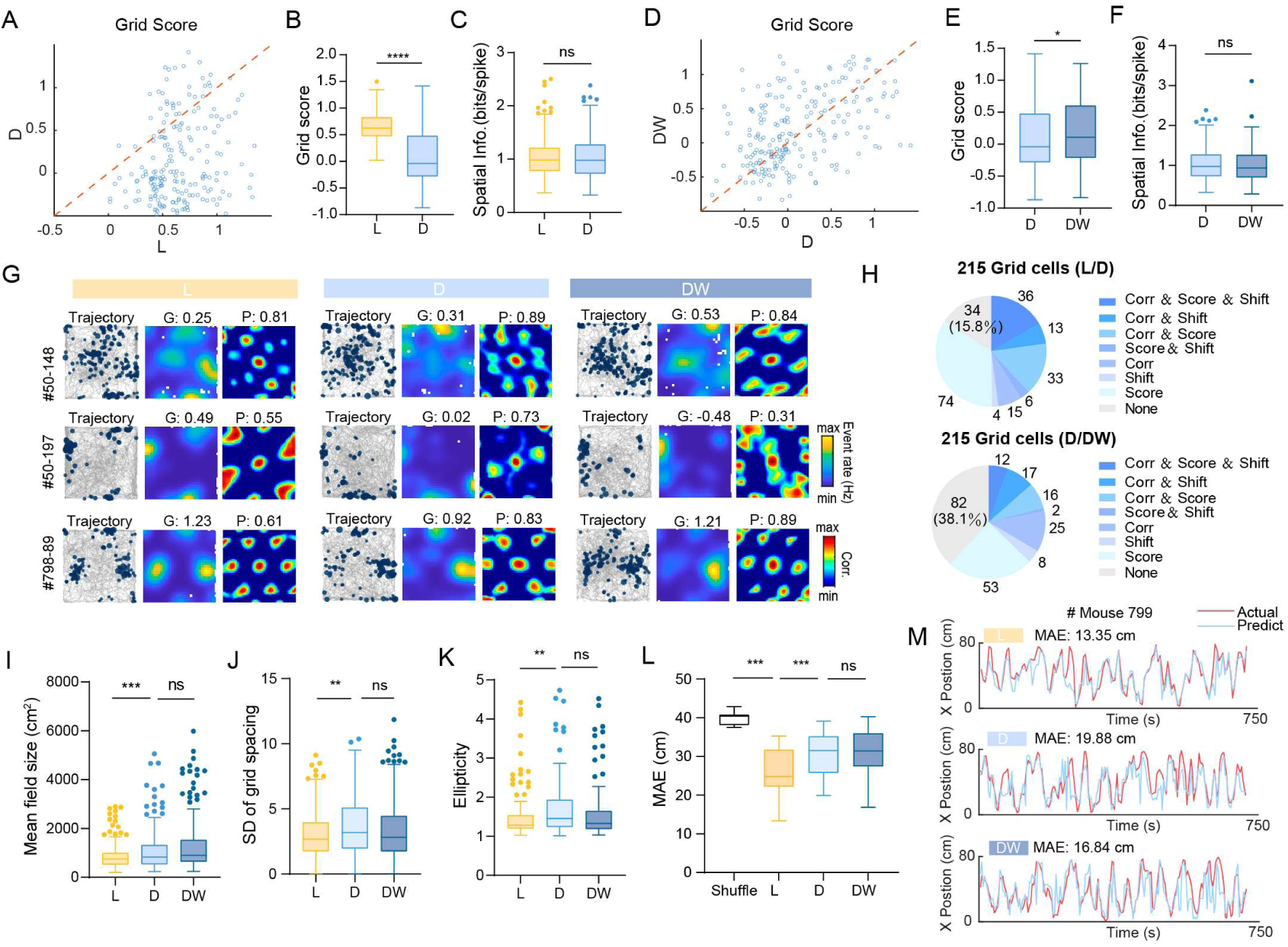
MEC border cells are modulated by both visual and whisker-mediated tactile inputs. (A) Scatter plot showing distribution of grid score for all grid cells with positive grid scores (n = 215) in Light (L) and Dark (D) conditions. (B) Box plot showing significant reduction in grid score after light-deprivation (L/D) (Wilcoxon signed-rank test, n = 215, P = 2.5 × 10^-27^). (C) No significant change in spatial information (bits/spike) was observed from L to D (Wilcoxon signed-rank test, n =215, P = 0.78). (D–F) Similar analyses as in (A–C), depicting changes in grid score and spatial information after whisker trimming (D/DW) (Wilcoxon signed-rank test, n = 215, grid score: P = 0.044; spatial information: P = 0.062). (G) Representative examples of color-coded rate maps and corresponding autocorrelograms for grid cells across different sessions. Grid score (G) and peak rate (P) are indicated. (H) Pie charts summarizing the proportions of grid cells exhibiting significant changes in rate map correlation, grid score, and/or peak firing location shift between session pairs. The upper pie chart shows changes from L to D, and the lower pie chart shows changes from D to DW. Cells were categorized based on whether they showed a significant decrease in rate-map correlation, a significant decrease in grid score, and/or a significant increase in peak firing location shift. (I) Box plots showing an increase in the average grid field size after light deprivation (L/D), with no further change after whisker trimming (D/DW) (Wilcoxon signed-rank test, L/D, n = 199 pairs, P = 0.00071; D/DW: P = 0.051). (J) Box plots showing a significant increase in the standard deviation (SD) of grid spacing after light deprivation, but not after whisker trimming (Wilcoxon signed-rank test, n = 179 pairs; L/D: P = 0.0038; D/DW: P = 0.20). (K) Box plots showing a significant increase in grid ellipticity after light deprivation, with no additional change after whisker trimming (Wilcoxon signed-rank test, n = 131 pairs; L/D: P = 0.0048 D/DW: P = 0.23). (L) Mean Absolute Error (MAE) in position prediction by all types of spatially-modulated cells across different conditions, showing that the spatial representation of the mice drifts in darkness (D) compared with its original state (L). In all experimental conditions, MAE is significantly lower than shuffle level (Wilcoxon signed-rank test, n = 13 mice, Shuffle/L: P = 0.00024; Shuffle/D: P = 0.00024; Shuffle/DW: P = 0.00049; L/D: P = 0.00024; D/DW: P = 0.59; L/DW: P = 0.00024). (M) Representative examples of actual (red) versus decoded (blue) positions across L, D, and DW sessions in one mouse, decoded from population activity of all spatially-modulated cells. MAE is indicated for each session. ****P ≤ 0.0001; ***P ≤ 0.001; **P ≤ 0.01; *P ≤ 0.05; ns, P > 0.05.

Following light deprivation, 143 of 215 grid cells (66.5%) exhibited a significant reduction in grid score relative to the baseline light session; 59 of 215 cells (27.4%) showed a significant shift in peak firing location; and 97 of 215 cells (45.1%) displayed a significant decrease in cross-session rate map correlation relative to within-session baseline. When all three metrics were considered jointly, only 34 of 215 cells (15.8%) showed no significant change in any measure (Figure 3H, upper). Following subsequent whisker trimming under dark conditions, 82 of 215 cells (38.1%) remained unaffected across all three metrics (Figure 3H, lower)—a proportion significantly greater than that observed under darkness alone (Chi-square test, χ² = 27.2, P = 1.8 × 10^-7^), further supporting the conclusion that visual deprivation exerts a greater disruptive influence on grid cell spatial coding than tactile deprivation.

To further validate these findings, we conducted a detailed examination of the firing maps of each grid cell (see Methods). The analysis revealed that, the mean field size of grid cells became larger in darkness (Figure 3I; L/D, Wilcoxon signed-rank test, n = 199, P = 0.00071), suggesting a potential expansion of the grid field. This change was accompanied by an increase in both the standard deviation of grid spacing (Figure 3J; Wilcoxon signed-rank test, n = 179, P = 0.0038) and ellipticity (Figure 3K; Wilcoxon signed-rank test, n = 131, P = 0.0048), indicating that grid pattern became less regular in darkness. These parameters remained stable following whisker trimming, suggesting that further deprivation of tactile inputs in animals already deprived of visual cues does not lead to a further deterioration of the regular firing patterns of individual grid cells (Figures 3I–K; mean field size: P = 0.051; SD of Grid spacing: P = 0.20; Grid ellipticity: P = 0.23;).

Additionally, we decoded the position based on the population activity of four mice that possessed more than 20 grid cells and compared their decoding performance to baseline conditions after sensory deprivation. Specifically, we split the baseline light session into two halves. The first half served as the training set, while the second half—along with the second halves of other sessions—was used to construct the test set for each experimental condition. The decoding performance improved as the number of participating grid cells increased (Figure S7F). Results showed that the mean absolute error (MAE) of the decoded position relative to the animals’ actual position slightly increased in the dark environment, although this difference did not reach statistical significance due to the limited sample size (Figure S7F; Wilcoxon signed-rank test, n = 4, Shuffle/L: P = 0.13; L/D: P = 0.13, D/DW: P = 0.63). Since grid cells are not the only contributors to spatial localization and the decoding results based solely on grid cells were not precise enough, we next included all functional cell types in the position decoder to assess the impact of darkness and whisker trimming on the animals’ spatial coding ability. MAE was significantly lower during the baseline light sessions compared to shuffled results (Figures 3L – M; see Methods). In darkness, the MAE increased compared to the light environment but remained unchanged after whisker trimming (Figure 3L; Wilcoxon signed-rank test, n = 13, Shuffle/L: P = 0.00024; L/D: P = 0.00024; D/DW: P = 0.59). This suggests that the original spatial-coding pattern of animals was influenced by the deprivation of visual inputs but remained unaffected by further deprivation of whisker-mediated tactile inputs. Notably, despite the sensory deprivation, the decoder across all sessions consistently produced a lower MAE than the shuffled results (Figure 3L), indicating that animals retain some ability to construct a cognitive map of the environment even in the absence of visual and tactile inputs, potentially through path integration. At the population level, grid scores and spatial information did not recover after re-exposure to the light condition (Figure S7G; DW/LW; Wilcoxon signed-rank test, n = 215; grid score: P = 0.53; spatial information: P = 0.30). However, a subset of grid cells exhibited more regular rate maps and increased grid scores, indicating pronounced heterogeneity within the population (Figure S7H).

Non-grid spatial cells exhibited response patterns comparable to those of grid cells. Among the 3,756 recorded cells, 501 were identified as non-grid spatial cells, with a mean spatial information of 1.26. Although their spatial coding was significantly disrupted in darkness, it was not further impaired by subsequent whisker trimming (Figure S8B; Wilcoxon signed-rank test, n = 501, L/D: P = 3.4 × 10^-23^ D/DW: P = 0.27). Notably, the mean field size of spatial cells increased significantly in darkness and was reduced again following whisker trimming (Figure S8C; Wilcoxon signed-rank test, n = 497, L/D: P = 9.9 × 10^-14^ D/DW: P = 0.026). Following lights-off, 109 of 501 spatial cells showed a concurrent significant decrease in spatial information (Score), significant alterations in their rate maps (Corr), and significant shifts in peak firing location (Shift). Only 96 cells (19.2%) remained completely unaffected across these three metrics. In contrast, after additional whisker trimming in darkness, 226 cells (45.1%) exhibited no significant changes in any of the three measures. Most spatial cells displayed increased out-of-field firing after lights-off, which did not worsen following whisker trimming (Figure S8D). In fact, a small subset of cells appeared to show slightly more precise spatial representations after trimming, possibly reflecting increased familiarity with the dark environment (Figure S8D).

### All spatially modulated cell types can anchor to tactile cues when tactile input dominates

Although grid scores of grid cells and spatial information of spatial cells did not significantly decline at the population level following whisker trimming (Figures 3E and S8A), and the accuracy of position decoding from neural activity was not further impaired in darkness (Figure 3L), a substantial subset of grid cells and spatial cells nonetheless exhibited significant changes in individual metrics after whisker deprivation (Figures 3H and S8D). Moreover, the reintroduction of light failed to restore population-level grid scores and spatial information (Figure S7G), suggesting that the removal of whisker-mediated tactile input did exert effects on some neurons. However, given that spatial representations in this experiment were established under lighted conditions with prominent visual cue cards (Environment 1, E1; Figure 1A), while the tactile cues present in the environment were comparatively subtle (e.g., minor textural variations and surface irregularities on the floor and walls of the paper-lined arena), the impact of whisker deprivation on grid and spatial cells was considerably more limited than that of light deprivation.

In naturalistic settings, rodents navigate environments of far greater complexity, characterized by objects of diverse materials, irregular terrain, and boundary structures with varied surface properties, often under conditions of dim ambient illumination. Under such circumstances, tactile input likely plays a role comparable to that of visual cues in supporting spatial orientation. To directly examine whether tactile cues can anchor spatial representations in the absence of visual information, we designed a new open-field arena rich in salient, polarizing tactile landmarks (Environment 2, E2). Two strips of sandpaper with distinct grit sizes were affixed to two walls of the arena, and an additional square piece of sandpaper was placed on the floor in one corner (Figure 4A, top). No visual cue cards were introduced, and the animals explored the arena in complete darkness throughout the training period. After approximately one week of habituation, we conducted a cue-rotation experiment to assess whether spatial representations rotated coherently with the tactile cues (Figure 4A, bottom).

**Figure 4.**
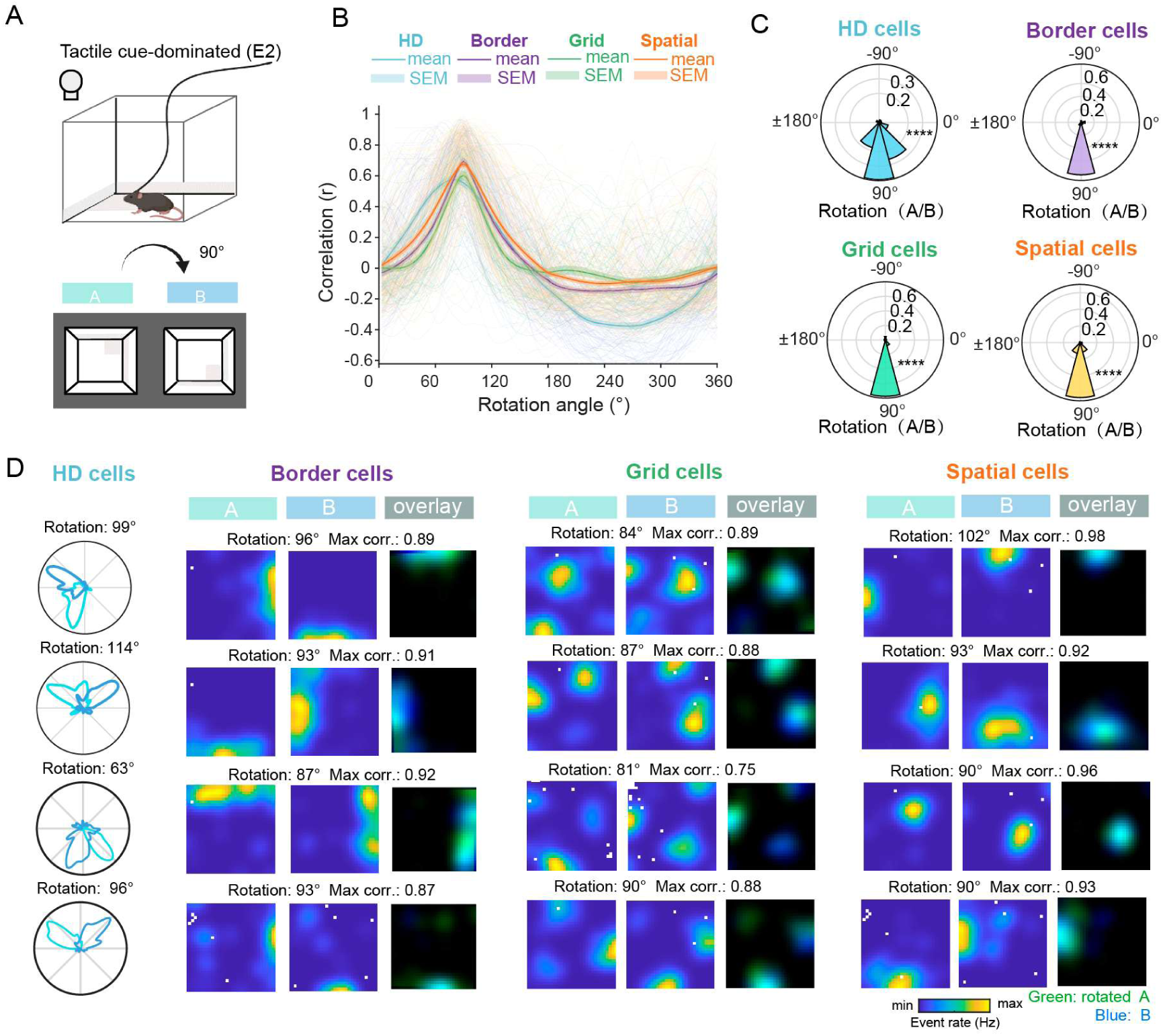
MEC spatial representations can anchor to tactile cues in darkness. (A) Top, schematic of the tactile cue-dominated environment (Environment 2, E2). Mice explored an open-field arena in complete darkness without any visual cue cards. Multiple tactile cues made of white sandpaper with different grit sizes were affixed to the arena floor and two walls. Bottom, After spatial representations were established (Session A), all tactile cues were rotated 90° clockwise, and the animals re-entered the environment for further exploration (Session B). (B) Rotation profiles of tuning curves for four classes of spatially modulated cells (HD, border, grid, and spatial cells) between Sessions A and B in the rotation experiments. The figure shows the rotation profiles from all cells included in the rotation experiments. For each cell, the tuning curve from Session A was circularly rotated in 3° increments, and the Pearson correlation between the rotated Session A tuning curve and the Session B tuning curve was computed at each rotation angle. Thin background traces represent individual cells, whereas thick traces denote the population mean for each cell type; shaded areas indicate SEM. Across cell types, the correlation profiles peaked near a 90° clockwise rotation, indicating that spatial representations were coherently rotated with the tactile cues. Clockwise rotation is defined as positive. (C) Rotation of tactile cues and corresponding rotation of neural representations. Top: Bottom: Polar plots showing the distribution of rotation angles for all cells within each of the four cell types. HD cells: n = 107, circular mean = 83.7°, Rayleigh test, Z = 65.9, P = 8.8 × 10^-28^; Border cells: n = 80, circular mean = 86.6°, Rayleigh test, Z = 60.57, P = 2.3 × 10^-25^; Grid cells: n = 62, circular mean = 90.3°, Rayleigh test, Z = 36.0, P = 1.6 × 10^-15^; Spatial cells: n = 180, circular mean = 88.9°, Rayleigh test, Z = 143.3, P = 2.1 × 10^-60^. (D) Representative tuning curves (rate maps) for each cell type. For HD cells, tuning curves from session A (light green) and session B (blue) are shown. For border, grid, and spatial cells, each row shows the rate map in session A, the rate map in session B, and their overlay. The overlay (dark green) illustrates the spatial correspondence between the session A rate map rotated clockwise by the optimal angle and the session B rate map. ****P ≤ 0.0001; ***P ≤ 0.001; **P ≤ 0.01; *P ≤ 0.05; ns, P > 0.05.

After mice established stable spatial representations in the initial configuration (Session A), the arena was carefully cleaned and the entire environment—including the sandpaper landmarks along with residual subtle tactile features on the walls and floor—was rotated 90° clockwise (Session B), and neural tuning curves were re-acquired. To quantify the alignment between sessions, each cell’s tuning curve from Session A was rotated in 3° increments and correlated with the corresponding Session B tuning curve across all rotation angles, generating a rotation-correlation profile (Figure 4B). The correlation peaked robustly at 90° clockwise rotation for nearly all recorded cells. Critically, spatial representations in all 6 of the 6 mice underwent coherent rotation (Figure S9). We defined the rotation angle as the angular offset at which the correlation between the rotated Session A and Session B tuning curves was maximized. The rotation angles of HD cells, border cells, grid cells, and spatial cells were all significantly clustered around 90° (Figure 4C; HD cells: n = 107, circular mean = 83.7°, Rayleigh test, Z = 65.9, P = 8.8 × 10^-28^; Border cells: n = 80, circular mean = 86.6°, Rayleigh test, Z = 60.57, P = 2.3 × 10^-25^; Grid cells: n = 62, circular mean = 90.3°, Rayleigh test, Z = 36.0, P = 1.6 × 10^-15^; Spatial cells: n = 180, circular mean = 88.9°, Rayleigh test, Z = 143.3, P = 2.1 × 10^-60^), consistent with the rotation of the tactile cues. Representative examples are illustrated in Figure 4D, demonstrating that individual neurons’ spatial representations from Session A, upon 90° rotation, closely matched those recorded in Session B. Collectively, these results demonstrate that in a visually deprived, completely dark environment, all classes of spatially modulated cells in the MEC can anchor to environmental tactile cues.

### Whisker trimming profoundly disrupts grid cells and spatial cells in a tactile cue-enriched environment

Given that various spatially modulated cells can anchor their firing to tactile cues within the open field in Environment 2 (Figure 4), whiskers—as the primary tactile sensory organ in rodents—likely play an indispensable role in this process. We therefore hypothesized that whisker trimming would exert a more pronounced impact on grid cells and other spatially modulated cells under these conditions compared to Environment 1 (Figures 1–3). To test this, we recorded MEC neuronal activity across 12 FOVs from 10 additional male mice and examined the effects of whisker trimming (Figure S10A). Whisker trimming did not produce any overt behavioral abnormalities (Figures S10B–E).

Consistent with observations in Experiment 1, tuning of HD cells and border cells was significantly disrupted following whisker deprivation (Figures S10F and S10I), as evidenced by marked reductions in tuning strength and firing specificity of individual cells (Figures S10G–H and J–K). In contrast to Environment 1, however, grid scores of grid cells also declined significantly after whisker trimming in Environment 2 (Figure 5A; Wilcoxon signed-rank test, n = 164, P = 1.5 × 10^-24^), though spatial information remained unchanged (Figure 5B; Wilcoxon signed-rank test, n = 164, P = 0.090). Moreover, grid cell properties that were unaffected by whisker trimming in Environment 1—including mean field size, standard deviation of grid spacing, and ellipticity— all showed significant changes in Environment 2 (Figures 5B–5E; mean field size: Wilcoxon signed-rank test, n = 154, P = 0.00047; SD of grid spacing: n = 154, P = 0.00034; Ellipticity: n = 130, P = 0.00084), indicating that the regularity of grid cell firing patterns was substantially degraded. Of the 164 grid cells recorded, only 24 (14.6%) exhibited no detectable change after whisker trimming across all metrics, the remaining cells showed varying degrees of grid score reduction, rate map reorganization, or peak firing location shifts (Figure 5F). Similarly, the spatial representations of spatial cells were severely disrupted by whisker deprivation (Figure 5G: Wilcoxon signed-rank test, n = 613, P = 4.1 × 10^-18^). Only 24.5% of spatial cells remained entirely unaffected in spatial information, rate map correlation, and peak firing location (Figure 5H). The position decoder trained on neuronal activity of all spatially modulated cells from the dark baseline session (D) showed a significant decrease in decoding accuracy following whisker trimming (DW), confirming that the loss of whisker-mediated tactile input under this conditions impaired the fidelity of animals’ spatial coding ability (Figures 5I–5J; Wilcoxon signed-rank test, n = 12, Shuffle/D: P = 0.00049, D/DW: P = 0.0024). At the single-cell level, the rate maps of the majority of grid cells and spatial cells underwent pronounced reorganization: some grid cells entirely lost their characteristic hexagonal firing patterns (Figure 5K), and some spatial cells showed complete abolition of spatial selectivity (Figure 5L).

**Figure 5.**
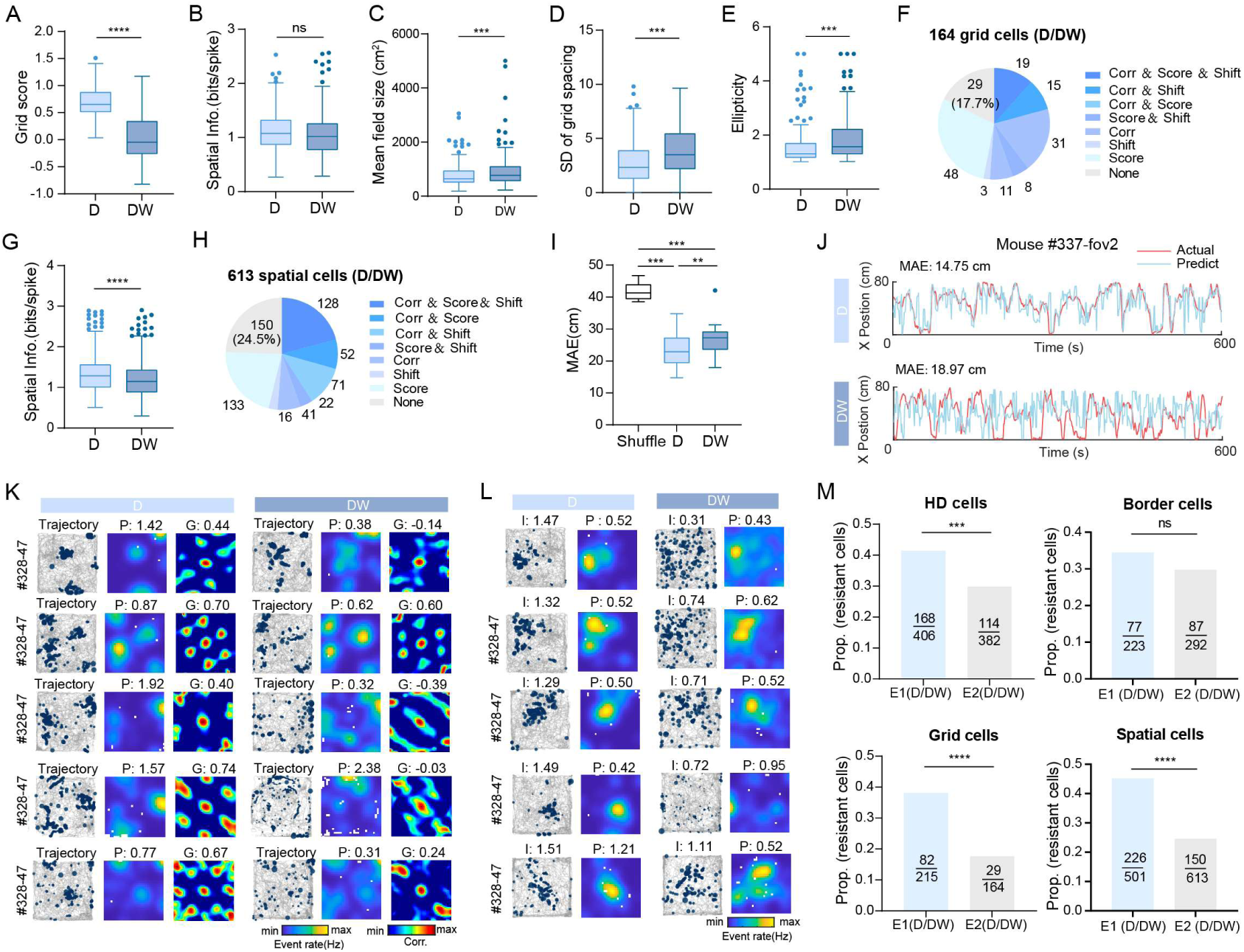
Whisker trimming strongly disrupts tuning of all types of spatially modulated cells in a tactile cue-enriched environment. (A) Box plot showing a significant decrease in grid score of grid cells after whisker trimming (D vs. DW) (Wilcoxon signed-rank test, n = 164, P = 1.5 × 10^-24^). (B) Box plot showing no significant decrease in spatial information (bits/spike) of grid cells after whisker trimming (Wilcoxon signed-rank test, n = 164, P = 0.090). (C) Box plot showing a significant increase in mean grid field size after whisker trimming (Wilcoxon signed-rank test, n = 154 pairs, P = 0.00047). (D) Box plot showing a significant increase in the standard deviation (SD) of grid spacing in DW compared with D (Wilcoxon signed-rank test, n = 154 pairs, P = 0.00034). (E) Box plot showing a significant increase in grid ellipticity after whisker trimming (D vs. DW) (Wilcoxon signed-rank test, n = 130 pair cells, P = 0.00084). (F) Pie chart summarizing the proportions of grid cells exhibiting significant changes in rate map correlation, grid score, and/or peak firing location shift between D and DW. Significance for each metric was determined using a shuffle-based baseline distribution. (G) Box plot showing a significant decrease in spatial information (bits/spike) of spatial cells after whisker trimming (Wilcoxon signed-rank test, n = 613, P = 4.1 × 10^-18^). (H) Pie chart summarizing the proportions of spatial cells exhibiting significant changes in rate map correlation, spatial information, and/or peak firing location shift between D and DW. (I) Mean absolute error (MAE) of position decoding using a decoder trained on neuronal activity from the D session. MAE was significantly lower than shuffle level in both sessions and significantly increased after whisker trimming (D vs. DW) (Wilcoxon signed-rank test, n = 12 FOVs from 10 mice; D vs. Shuffle: P = 0.00049; DW vs. Shuffle: P = 0.00049; D vs. DW: P = 0.0024). (J) Representative example of actual (red) versus decoded (blue) position in D and DW sessions from one mouse. (K) Representative examples of grid cells recorded in D and DW sessions, showing trajectories, spatial rate maps, and spatial autocorrelograms. Grid score (G) and peak rate are indicated. (L) Representative examples of spatial cells recorded in D and DW sessions, showing spike locations and spatial rate maps. Spatial information (I) and peak rate (P) are indicated. (M) Comparison of the proportions of whisker-deprivation-resistant cells across two environments (E1 vs. E2). Resistant cells were defined as those showing no significant change in any metric after whisker trimming (i.e., classified as “None”). The fractions of resistant cells differed significantly between environments for HD cells (chi-square test, P = 0.00026), grid cells (chi-square test, P = 1.5 × 10^-5^), and spatial cells (chi-square test, P = 6.0 × 10^-5^), but not for border cells (chi-square test, P = 0.38). ****P ≤ 0.0001; ***P ≤ 0.001; **P ≤ 0.01; *P ≤ 0.05; ns, P > 0.05.

In this experiment, two mice underwent a second round of whisker trimming after their whiskers had regrown to sufficient length following the initial deprivation. During the second manipulation, neural activity was recorded from newly imaged fields of view. Notably, repeated whisker trimming produced effects on all classes of spatially modulated cells that were comparable in magnitude to those observed after the first trimming (Figure S10L). The reproducibility of these effects argues against the possibility that the observed degradation in spatial representations was driven by transient behavioral abnormalities, stress responses, or incomplete task adaptation.

### Spatial representations anchored to tactile cues are preferentially disrupted by whisker trimming

As Environment 2 provides rich tactile but no visual cues, spatially modulated cells there are expected to rely predominantly on whisker-mediated input for spatial coding, predicting greater disruption from whisker trimming than in Environment 1. Consistent with this, the proportion of whisker deprivation-resistant cells (defined as those showing no significant change in any spatial metric following trimming) was significantly lower in Environment 2 than in Environment 1 for HD cells, grid cells, and spatial cells (Figure 5M). This finding further proves that the effects of whisker trimming cannot be solely attributed to trimming-induced alterations in locomotor behavior or arousal state. Border cells were the exception, showing no significant difference between environments (Figure 5M), likely because they mainly rely on whisker-mediated boundary sensing regardless of the broader tactile context.

Rodents rely on their whiskers to detect surface textures. In our arena, the sandpaper patch positioned in the upper-right corner provided the dominant tactile cue apart from the enclosing walls. This cue could therefore serve either as a local tactile landmark directly anchoring the firing fields of spatially modulated cells or as a stable reference to correct errors accumulated during path integration. We thus hypothesized that whisker trimming would preferentially disrupt firing patterns within the sandpaper region-of-interest (ROI) relative to the rest of the arena. To test this hypothesis, we first computed population-averaged rate maps for each of the four spatially modulated cell types by averaging event rates across all cells of a given type within each spatial bin (Figures S11A–D). HD cells and border cells showed no pronounced reduction in mean firing rate within the ROI following whisker trimming, consistent with the fact that HD cell activity is basically independent of spatial location and border cell firing is primarily constrained to environmental boundaries. In contrast, grid cells and spatial cells exhibited a markedly greater reduction in population mean rate within the ROI compared to other regions of the arena after whisker trimming (Figure S11C–D). We next examined whether whisker trimming preferentially affected cells whose spatial representations depended on tactile information from the ROI. Grid cells were classified as ROI cells if at least one grid field was located within the ROI, and as non-ROI cells otherwise (see Methods), yielding 44 ROI grid cells and 120 non-ROI grid cells (Figure S11E). Grid scores decreased significantly in both groups after whisker trimming and the magnitude of reduction did not differ between ROI and non-ROI grid cells (Figure S11F). However, field-level analysis revealed a selective effect: grid fields located within the ROI (ROI fields) were significantly more likely to disappear following whisker trimming than non-ROI fields (Figures S11G–H, see Methods). Field was considered lost if no corresponding field could be identified in the subsequent session, or if the spatial overlap between the new field and the original field was less than 20%. We performed a parallel analysis for spatial cells. Based on the location of their primary (largest) firing field, 52 cells were classified as ROI spatial cells and 561 as non-ROI spatial cells (Figure S11I, see Methods). ROI spatial cells indeed exhibited a significantly greater reduction in spatial information compared to non-ROI spatial cells following whisker trimming (Figure S11J). Moreover, spatial fields located within the ROI were more likely to disappear than those outside the ROI (Figures S11K–L). This finding hints that tactile floor cues contribute to stabilizing spatial representations, and that removal of whisker-mediated tactile input preferentially disrupts the coding of fields anchored to the salient landmarks.

### Cross-day stability modestly predicts vulnerability to sensory deprivation

To examine whether long-term representational stability predicts sensitivity to sensory deprivation, we performed five days of recordings in Environment 2 in three mice. The imaging fields of view remained stable and the spatial footprints looked very similar across all five sessions (Figure S12A), enabling reliable longitudinal tracking of the same neurons. In darkness, all four classes of spatially modulated cells could establish consistent spatial representations across sessions, indicating that tactile cues within the environment were sufficient to support stable coding (Figure S12B).

On the fifth day, whisker trimming was performed, allowing us to directly compare deprivation-induced changes in spatial tuning with each cell’s intrinsic cross-day stability. For HD cells, whisker-trimming-induced reductions in tuning strength (ΔMVL, D/DW) were significantly correlated with cross-day variability (SEM of MVL across five days), such that cells exhibiting greater baseline variability showed larger deprivation-induced declines (Figure S12C, left). Moreover, cells with higher cross-day stability (mean correlation across five days) exhibited better preservation of tuning after trimming (Figure S12C, right). In contrast, no significant relationships were observed for border cells (Figure S12D). For grid cells, deprivation-induced reductions in grid score were significantly associated with cross-day variability, although the cross-day spatial correlation did not predict the magnitude of trimming effects (Figure S12E). Finally, spatial cells showed no significant association between cross-day stability and deprivation-induced changes in spatial information or spatial correlation (Figure S12F). Together, these results indicate that long-term representational stability modestly predicts vulnerability to sensory deprivation in a subset of tuning metrics, particularly for HD and grid cells. Thus, while tactile cues can support stable spatial representations in darkness, cells with less stable baseline tuning are more susceptible to disruption following whisker-mediated sensory loss.

### Environmental boundaries provide dominant stabilizing input after multisensory deprivation

Although visual cues were absent in Environment 2, a subset of cells maintained stable firing patterns following whisker trimming, showing no detectable changes in tuning strength or tuning curves (Figures 5F, 5H, S10G, and S10J). Given the complexity of the rodent somatosensory system, tactile inputs beyond whiskers—such as those conveyed via the paws or nose—may contribute to sustaining residual spatial representations. We therefore examined whether plantar tactile input supports spatial coding under whisker deprivation. Because complete elimination of paw sensation is technically impractical and would likely disrupt locomotion, we instead removed the dominant floor tactile cue—the sandpaper patch — which constituted the most salient remaining plantar-accessible landmark after whisker trimming. Four mice re-explored the arena under this condition (DWR; Figure S13A). Removal of the sandpaper patch did not induce additional significant changes in tuning metrics across spatially modulated cell types (Figure S13), nor did it impair positional decoding accuracy (Figure S13; Wilcoxon signed-rank test, n = 4 mice, P > 0.99). Importantly, this manipulation reduced—but did not abolish—plantar tactile input.

Environmental boundaries are known to provide strong corrective signals for path integration. In the arena lacking other salient protrusions, the remaining tactile feedback may be predominantly derived from physical contact between the animal’s nose or body and the surrounding walls. To assess the contribution of boundary-derived input, we eliminated access to the walls while preserving the floor landmark. In six mice tested on an elevated platform, the arena walls were removed following the DW session (DWR′; Figure 6A), preventing wall contact without altering the sandpaper cue. Under these conditions, spatial tuning was profoundly disrupted across all classes of spatially modulated cells (Figures 6B–D and S14A–B; Wilcoxon signed-rank test; n = 276 HD cells, MVL: P = 0.00020, Head Info.: P = 2.6 × 10^-8^; n = 190 border cells, border score: P = 2.9 × 10^-9^, Spatial Info.: P = 3.7 × 10^-15^ ; n = 94 grid cells, grid score: P = 0.0083; n = 328 spatial cells, Spatial Info.: P = 3.6 × 10^-18^). The spatial information of grid cells, which had remained stable under darkness and whisker trimming, was significantly reduced following wall removal (Figure S14C; Wilcoxon signed-rank test, n = 94, P = 0.028). Positional decoding errors increased markedly relative to the DW session (Figures 6F and S14D; Wilcoxon signed-rank test, n = 6 mice, P = 0.031). Cells previously unaffected by whisker trimming (e.g., the first two HD cells in Figure 6G) now showed significant changes in MVL and PFD, suggesting that their residual stability in DW sessions was supported by boundary-derived sensory feedback (Figures 6G–J). Rather than remapping, most border cells, grid cells, and spatial cells failed to meet classification criteria following wall removal. Quantitatively, 50.7% (140/276) of baseline HD cells remained classifiable in session DWR′, whereas only 12.1% of border cells (23/190), 11.7% of grid cells (11/94), and 15.5% of spatial cells (51/328) met classification thresholds, highlighting the strong dependence of allocentric spatial coding on boundary information.

**Figure 6.**
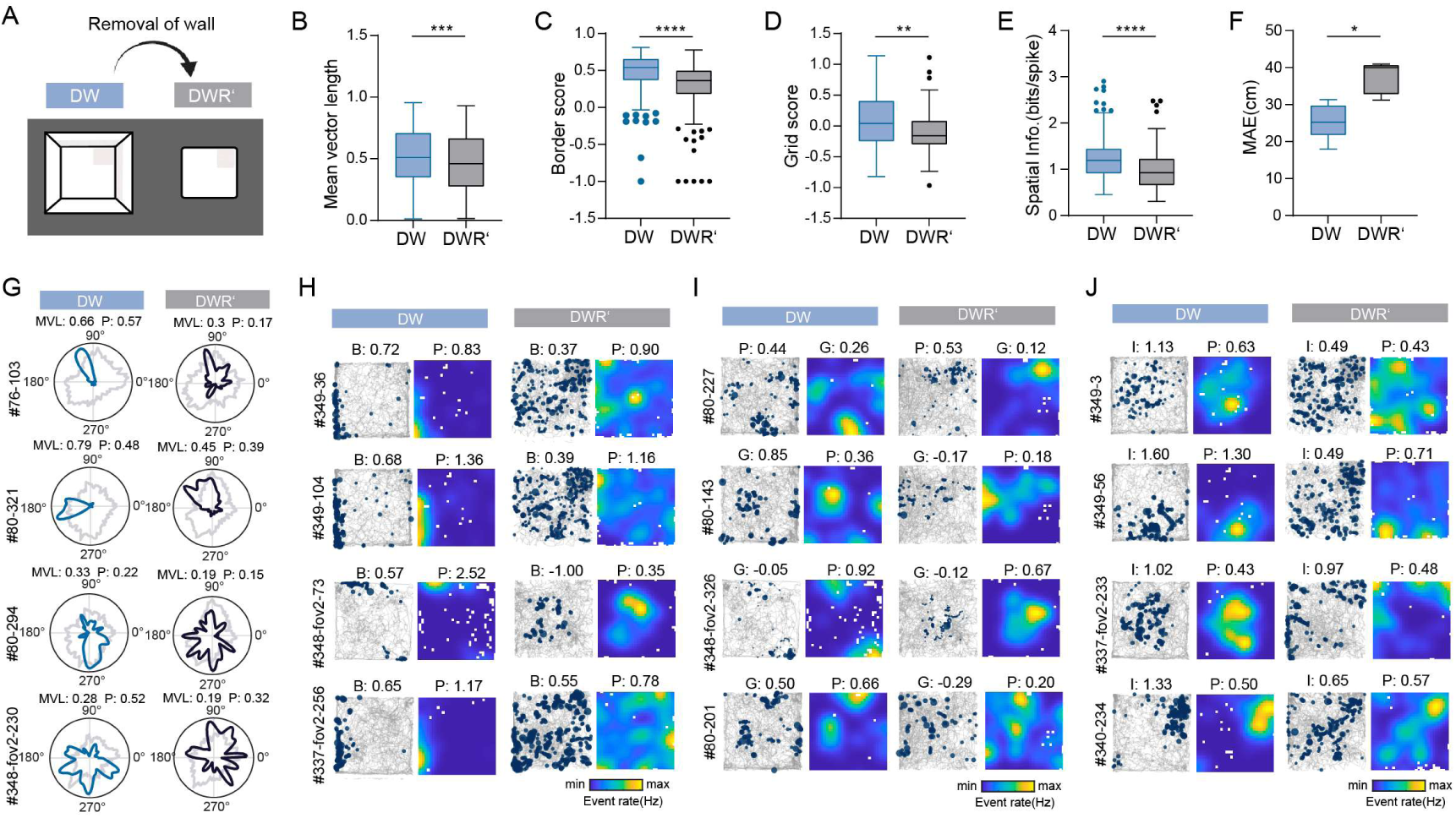
Removal of the arena wall significantly disrupts spatial representations following combined sensory deprivation. (A) Schematic illustration of the experimental manipulation, showing the transition from DW (Dark, Whisker-trimmed) to DWR′ by removing the arena wall. (B) Box plot showing a significant decrease in MVL of HD cells in DWR′ compared with DW (Wilcoxon signed-rank test, n = 276, P = 0.00020). (C) Box plot showing a significant decrease in border score of border cells after wall removal (Wilcoxon signed-rank test, n = 190, P = 2.9 × 10^-9^). (D) Box plot showing a significant decrease in grid score of grid cells after wall removal (Wilcoxon signed-rank test, n = 93, P = 0.0083). (E) Box plot showing a significant decrease in spatial information (bits/spike) of spatial cells after wall removal (Wilcoxon signed-rank test, n = 328, P = 3.6 × 10^-18^). (F) Mean absolute error (MAE) of position decoding using a decoder trained on neuronal activity from the baseline session (D) was significantly increased in DWR′ compared with DW, indicating reduced decoding accuracy after wall removal (Wilcoxon signed-rank test, n = 6 mice, P = 0.031). (G–J) Representative examples of HD cells (G), border cells (H), grid cells (I), and spatial cells (J) recorded in DW and DWR′ sessions. Corresponding cell-type metrics (MVL, border score (B), grid score (G), spatial information (I) and peak rate (P) are indicated. ****P ≤ 0.0001; ***P ≤ 0.001; **P ≤ 0.01; *P ≤ 0.05; ns, P > 0.05.

In two of six mice, exploration trajectories became biased toward the sandpaper region after wall removal, which closely matching the size and shape of the tactile patch (Figure S14F). Although observed in a subset of animals, this pattern suggests that, in the absence of boundary cues, the floor landmark may serve as a residual spatial reference detectable via plantar sensation. Population-averaged border-cell maps showed reduced firing along former arena boundaries but increased activity along the edges of the sandpaper patch in DWR′ (Figure S14G, top). After normalization, a new firing band emerged along the vertical edge of the tactile cue (Figure S14G, bottom), indicating a highly complex and adaptable spatial representation system in the MEC. In this context, animals may rely on plantar tactile input to detect the remaining sandpaper landmark on the floor and strengthen the encoding of that boundary.

### Subgroup of cells in MEC are modulated by whisker movement

During free exploration, rodents use their whiskers bilaterally to interact with objects, providing essential tactile inputs. Previous research has shown that the CA1 region of the hippocampus responds to both mechanical and passive whisker stimulation, potentially contributing to the updating of the spatial cognitive map^27^. Furthermore, it has been established that place fields in the hippocampus are modulated by tactile inputs from whiskers, although there is variability among individual cells^5^. Our earlier results indicated that spatially modulated cells in MEC can anchor to the environmental tactile cues in darkness and alter their coding in response to the tactile deprivation. We therefore asked whether the MEC contains neurons that participate in the processing of tactile information.

We conducted whisker-related experiments in ten male mice, three of which had also undergone whisker trimming in Environment 2. Each animal participated in three experimental sessions (Figure 6A). In the “spontaneous-whisking” session (Spont.), mice were head-fixed, and their spontaneous whisker movements along with corresponding neuronal activity were recorded simultaneously. Due to minimal whisker movement under head-fixed conditions, we performed an additional “stimulated-whisking” session (Stim.), during which an automated telescopic rod with a sandpaper sheet at its end was used to enhance whisker activity at regular intervals. In the final “free exploration” session, mice were allowed to explore an open field, enabling us to assess the spatial-coding properties of the neurons recorded in the previous sessions. Given that whiskers on the same side exhibit highly similar movement^42^ (Figure S15A), we selected the most distinguishable whisker for each animal for further analysis. To minimize false positives resulting from the low whisker movement observed during the “spontaneous-whisking” session, we primarily used data from the “stimulated-whisking” session to identify whisker-responsive cells, with the “spontaneous-whisking” session serving to validate our findings. The position of the selected whisker was extracted from each frame, and the corresponding angular position (*θ*) was calculated. Angular velocity (*θ′*) was derived by calculating the change in angular position between two adjacent frames. We referred smoothed angular velocity (*θ′*) as whisker movement and further calculated its Pearson correlation coefficient with each neuron’s denoised ΔF/F trace (see Methods). We used this correlation-based approach rather than comparing neuronal activity between stimulated and non-stimulated intervals due to the low temporal resolution of two-photon imaging and the potential for time-delayed responses, which could result in a mismatch between the timing of stimulation and neuronal activity. To identify whisker-responsive cells, both correlation strength and within-session stability criteria were applied (see Methods). Specifically, the correlation between *θ′*and neural activity was calculated separately for the first and second halves of the “stimulated-whisking” session. Only neurons whose correlation coefficients exceeded the 95^th^ percentile of the shuffled distribution in both halves were classified as whisker-responsive cells (Resp.); all others were categorized as non-responsive to whisker movement (Non-Resp.).

Of the 2,187 recorded neurons, 191 (8.7%) were identified as “whisker-responsive” cells, characterized by an increased ΔF/F in response to increased whisker movement (Figure 6B). In an independent “spontaneous-whisking” session, these whisker-responsive cells also exhibited significantly higher correlations with whisker movement compared to whisker-non-responsive cells (Figures 7C and S15E; Welch’s *t*-test, n = 1,996 and n = 191, P = 9.5 × 10^-46^), although correlation values were reduced relative to the “stimulated-whisking” session, likely due to weaker whisker activity in the absence of stimulation (Figure 7D; Wilcoxon signed-rank test, n = 191, P = 0.0021). Importantly, when correlations were recalculated using deconvolved calcium signals instead of denoised ΔF/F traces, whisker-responsive cells still showed significantly higher correlations than non-responsive neurons (Figure S15B; Welch’s *t*-test, n = 1,987 and n = 191, P = 7.8 × 10^-13^), and this effect was consistently observed across both independent sessions (Figure S15C; Welch’s *t*-test, P = 5.5 × 10^-6^) and demonstrated stable performance in the “spontaneous-whisking” session (Figure S15D; Wilcoxon signed-rank test, n = 191, P = 0.41).

**Figure 7.**
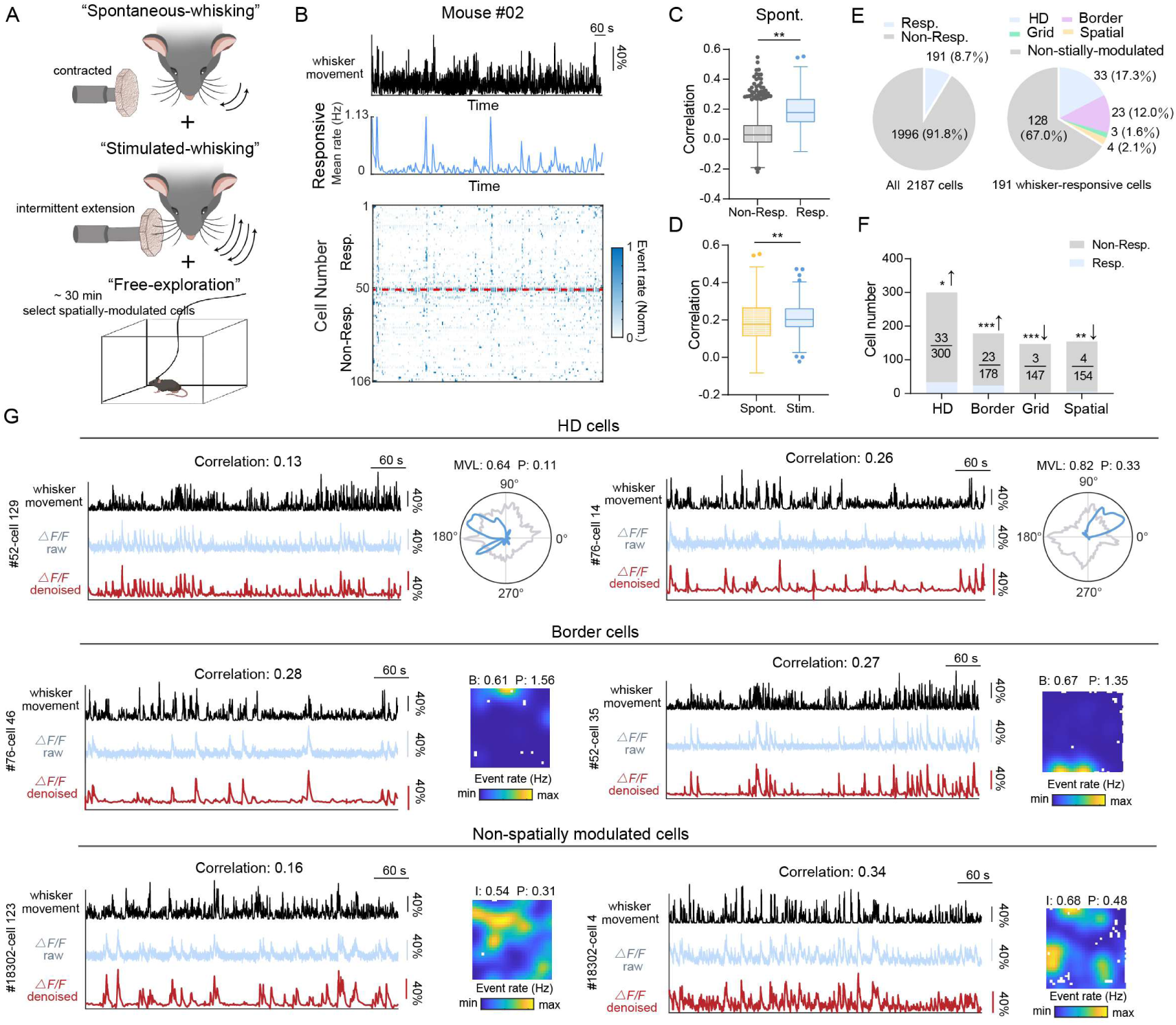
Identification of whisker-responsive cells in the MEC. (A) Experimental design and setup illustrating three whisker-related sessions: "spontaneous-whisking (Spont.)", "stimulated-whisking (Stim.)" and "free exploration". During the "stimulated-whisking" session, an automatic retractable rod with sandpaper pieces at the end was used to enhance whisker activity. (B) Top: Whisker movements during the "stimulated-whisking" session for a representative mouse. Middle: The mean rate curve of all “whisker-responsive” cells identified in this mouse demonstrates a high degree of consistency with whisker movement. Bottom: Heatmap showing the standardized event rate of all identified “whisker-responsive”(Resp.) and “whisker-non-responsive”(Non-Resp.) neurons in the same mouse, with white color indicating the lowest event rate and dark blue indicating the highest. (C) During the "spontaneous-whisking" session, neuron activity of "whisker-responsive” cells identified in the "stimulated-whisking" session also exhibits significantly higher correlations with whisker movement compared to "whisker-non-responsive” cells (Welch’ s *t*-test, n = 1,996 (Non-Resp.) and n = 191 (Resp.), P = 9.5 × 10^-42^). (D) Whisker-responsive cells defined during the whisker-stimulating session continue to show robust correlations between neuronal activity and whisker movement during the spontaneous-whisking session, although the correlation strength is reduced compared with the stimulated-whisking session (Wilcoxon signed-rank test, n = 191, P = 0.0021). (E) Left: Proportion of “whisker-responsive” and “whisker-non-responsive” cells among all recorded cells (total n = 2,187 cells). Right: Distribution of spatially-modulated cell types among whisker-responsive cells (total n = 191 whisker-responsive cells). (F) Proportion of whisker-responsive cells across different spatially modulated cell types. Whisker-responsive cells are significantly enriched among HD and border cells relative to chance levels predicted by a random sampling model, whereas grid cells and spatially selective cells show significant depletion. Expected counts were computed based on the relative abundance of each cell type across all recorded neurons (hypergeometric test; HD: expected = 24, observed = 33, P(enrich) = 0.036; Border: expected = 15, observed = 28, P(enrich) = 0.00036; Grid: expected = 12, observed = 3, P(deplete) = 0.00084; Spatial: expected = 13, observed = 4, P(deplete) = 0.0015). (G) Representative examples of whisker-responsive cells with distinct spatial tuning properties. From top to bottom: HD cells, border cells and non-spatially modulated cells. Black traces represent whisker movement, blue traces show raw ΔF/F calcium signals, and red traces indicate denoised ΔF/F. Right panels show the corresponding head-direction tuning curves or spatial firing rate maps recorded during the “free-exploration” session. ****P ≤ 0.0001; ***P ≤ 0.001; **P ≤ 0.01; *P ≤ 0.05; ns, P > 0.05.

Among the identified whisker-responsive cells, the majority (67.0%) did not exhibit spatial tuning during free exploration. The remaining 63 of 191 cells (33.0%) displayed spatially modulated properties, with HD cells comprising the largest fraction (17.3%), followed by border cells (12.0%), spatial cells (2.1%), and grid cells (1.6%) (Figure 6E). These neurons showed strong correlations with whisker movement during the “stimulated-whisking” session while also exhibiting head-direction or spatial tuning during free exploration (Figures 7G and S15F). Given that the simultaneous encoding of whisker movement and spatial information during natural exploration may impose computational constraints, it is possible that these dual-tuned neurons receive input from non-spatially modulated whisker-responsive cells. Consistent with this possibility, whisker-responsive cells exhibited a modest tendency toward anatomical clustering (Figures S15G – H; see Methods), which may reflect potential local connectivity. Because the total number of each spatially modulated cell type differed, we assessed whether the observed overlap between whisker-responsive cells and spatially tuned populations deviated from chance expectation. Using the overall proportion of whisker-responsive cells and the total counts of each cell type, we calculated the expected overlap under a hypergeometric distribution and compared it with the observed values. This analysis revealed significant enrichment of HD cells (expected = 24, observed = 33, P = 0.036) and border cells (expected = 15, observed = 28, P = 3.6 × 10⁻⁴), along with significant depletion of grid cells (expected = 12, observed = 3, P = 8.4 × 10⁻⁴) and spatial cells (expected = 13, observed = 4, P = 0.0015).

Additionally, we replicated the “stimulated-whisking” session in four independent mice under head-fixed conditions using a high-speed infrared camera (120 Hz) to more precisely capture whisker movement. A population of neurons exhibiting strong correlations between neuronal activity and whisker angular velocity was again identified, further validating our previous findings (Figure S16A). The correlation strength of whisker-responsive cells did not differ significantly between the two experimental setups (Figure S16B; Welch’s *t*-test, P = 0.38). In this experiment, we also employed a secondary infrared camera (50 Hz) positioned laterally to record paw movements and compute paw velocity, which served as an indicator of the animal’s body locomotion. Although some whisker-responsive cells displayed calcium activity patterns closely resembling whisker angular velocity, their correlations with paw movement were minimal or absent (Figures S16C–D). Because whisker and paw movement intensities can at times be tightly coupled and difficult to dissociate, these results indicate that at least a subset of whisker-responsive cells specifically encodes whisker angular velocity rather than merely reflecting global behavioral state changes.

## Discussion

Effective spatial navigation is fundamentally dependent on reliable references, which are derived from both internal and external sensory information^43^. When spatial representation was primarily established by visual cues, whisker trimming affected HD and border cells but had minimal impact on grid cells and non-grid spatial cells. In contrast, when the environment was enriched with tactile cues but lack of other sensory inputs, all cell types anchored their firing to tactile landmarks and formed stable representations; under these conditions, whisker trimming profoundly disrupted spatial tuning across cell classes. In addition to whisker-mediated input, tactile signals conveyed through the nose or plantar surface may also contribute to maintaining spatial representations. Together, these findings underscore the complexity of sensory integration in spatial navigation and highlight flexible strategies employed by the MEC to process external sensory information under varying environmental contexts. Moreover, a subset of MEC neurons displayed stable tuning to whisker movement, suggesting that they may participate in conveying tactile-related signals to the spatial navigation network. This observation further raises the possibility of functional interactions between the MEC and tactile-processing brain regions.

All known types of spatially modulated cells are modulated to varying degrees by sensory information, with HD cells being particularly sensitive. For instance, HD cells in the subiculum respond to environmental rotation^44^. In blind mice, HD cells in the anterior dorsal nucleus (ADN) use olfactory cues to finely encode head direction^36^. Moreover, disruption of vestibular input impaired the head direction tuning in the anterior thalamus ^45^, leading to subsequent effects on the place cells in the hippocampus^46^. Our results showed that head direction tuning in the MEC was impaired following the deprivation of visual and whisker-mediated tactile inputs (Figure 1 and Figure S3). The disruption observed under light deprivation may be explained by both direct and indirect anatomical connectivity between the MEC and visual cortical areas^47,48^, which provide a pathway through which visual information can influence spatial representations. In addition, it is well established that visual landmarks exert strong control over head direction tuning^49,50^. Numerous studies have demonstrated that removal of visual cues degrades HD stability^22^, whereas reintroduction of visual landmarks restores directional tuning^23,51^. However, we did not observe a robust recovery of HD tuning following light reinstatement after whisker trimming. This persistent impairment likely reflects the sustained consequences of whisker deprivation on the HD network, such that only a subset of cells whose directional tuning was primarily governed by visual input—and thus minimally affected by whisker trimming—were able to recover their tuning strength following light reinstatement (Figure S3I). Moreover, in some animals, LDA revealed that, although the HD population activity did not return to its initial state following light reinstatement, it nevertheless retained the capacity to encode head direction within LW session (Figure S3J). This may reflect the tendency of HD population activity, following an extended period of altered experience between the two light sessions, to stabilize around a shifted attractor state upon reintroduction of visual input^49,52^. Whiskers play a critical role in rodents’ environmental exploration and object recognition. They allow animals to perceive surface textures, detect contours and obstacles and localize nearby boundaries. These tactile cues likely contribute to the construction of spatial representations and may support the directional tuning of HD cells within a cognitive map. In addition, whisker activity is closely linked to head movements in rodents, with the right-left asymmetry of whisker positioning preceding head movement^53^, enabling obstacle detection and direction determination. Head turns and eye movements are flexibly coordinated, allowing animals to effectively integrate visual, vestibular, and whiskers play a critical role somatosensory information^53–55^. In our study, whisker trimming resulted in a measurable reduction in head angular velocity and angular acceleration (Figures S4I-J), which may reflect compromised balance or altered vestibular input during head turns. This reduction may affect the activity of angular velocity cells^56,57^—potentially involved in vestibular ocular reflex and in maintaining or updating head direction tuning^58^. Therefore, the impact of whisker deprivation on HD tuning may partially arise from changes in vestibular processing, especially in Environment 1. Thus, HD cell likely integrate multimodal sensory inputs together with self-generated signals to maintain accurate directional coding.

Border cells, another crucial type of spatially modulated cell, are also significantly impacted by light deprivation and whisker trimming (Figure 2 and Figure S10). Animals may rely on visual cues to estimate their distance from environmental boundaries, and prior studies have shown that boundary-related firing fields can persist even when boundary information is provided predominantly through visual input^59^. However, direct contact with walls via whiskers may allow border cells to refine their firing fields with greater spatial precision. Because environmental boundaries provide corrective signals for path integration, and border cells are thought to contribute to this process by modulating grid-cell firing fields^11^, our findings suggest that boundary-dependent spatial coding is sensitive to the quality of available sensory input. In darkness, the precision of distance estimation based on vision is reduced, and whisker trimming may further degrade boundary-related feedback. Consistent with this interpretation, we observed that border cells preferentially fired when the animal was oriented toward the wall (Figures 6F–G). Notably, this directional bias was completely abolished when the surrounding walls were removed (Figure S14E), and most border cells lost spatially specific firing rather than merely exhibiting remapping (Figure 6H). These findings indicate that the maintenance of boundary fields depends strongly on external sensory inputs. This result differs from that reported by Solstad et al.,^16^ likely because, in our wall-removal sessions, animals lacked both visual cues and the ability to detect environmental boundaries via whisker contact. Under these conditions, boundary-related sensory information was almost completely eliminated.

In the illuminated Environment 1 containing salient visual cue cards, spatial representations were likely established primarily on the basis of visual information. Under these conditions, removal of visual cues profoundly disrupted firing patterns in grid cells and spatial cells, whereas whisker trimming produced only modest effects (Figure 3 and Figure S8). In contrast, Environment 2 lacked visual input but contained a salient polarized tactile landmark. In this context, tactile cues likely contributed both to anchoring firing fields and to correcting accumulated path integration errors in the absence of vision. Accordingly, whisker trimming in Environment 2 led to pronounced reductions in spatial specificity in both grid and spatial cells, accompanied by alterations in field properties and decreased positional decoding accuracy (Figures 5A–L). Moreover, when multiple tuning metrics were considered jointly, the proportion of HD, grid, and spatial cells that maintained stable tuning in Environment 2 was significantly lower than the proportion of whisker-trimming-resistant cells observed in Environment 1 (Figure 5M). These findings suggest that the MEC flexibly constructs spatial maps depending on the dominant sensory modality available in the environment, spatial representations primarily anchored to tactile information appear more vulnerable to whisker deprivation than those supported by vision.

Unlike vision, which originates from retinal photoreceptors, tactile inputs arise from mechanoreceptors distributed across the body rather than being confined to whisker follicles^60^. Consequently, whisker trimming alone cannot abolish all somatosensory input, consistent with the persistence of spatial selectivity in a subset of cells under combined darkness and whisker deprivation. Additional removal of surrounding walls further destabilized spatial coding (Figure 6 and Figure S14), suggesting that somatosensory signals from the nose also contribute to navigation. Whereas boundary manipulations have been shown to induce grid cell remapping^16^, our results reveal a more severe outcome: when both visual input, whisker mediated tactile input and boundary-related physical feedback were eliminated in the DWR′ condition, most grid cells lost their hexagonal firing patterns and most spatial cells lost spatial specificity. Nevertheless, a minority of spatially modulated cells remained classifiable, indicating that self-motion-related signals—such as vestibular and speed inputs—can partially sustain spatial representations when external sensory cues are drastically reduced. Notably, the disruption caused by single sensory deprivation was milder than that caused by medial septum inactivation or passive transport, both of which abolish grid-like firing entirely^61,62^. Thus, selective deprivation of individual modalities produces partial degradation of grid firing, whereas collective reduction of sensory inputs — particularly elimination of boundary perception — profoundly disrupts spatial coding. Importantly, our manipulations likely did not eliminate all somatosensory input. Tactile signals from the paws were attenuated by removing the primary sandpaper landmark, and inputs from the nose were reduced by wall removal; however, residual somatosensory information may have persisted. Future studies employing more direct circuit-level interventions, such as transient inactivation of somatosensory pathways, will be required to more definitively establish the causal contribution of tactile signals to spatial representations.

Facial whiskers are vital sensory organs for nocturnal rodents like mice and rats, providing essential tactile inputs. The active movement of whiskers, known as whisking^63^, facilitates environmental exploration and triggers sparse neuronal activity patterns in the primary somatosensory cortex (S1)^64^. Previous research has shown that rodents can use their whiskers to assess aperture widths with an accuracy finer than 5 mm^65^. Additionally, blind rats rely heavily on whisker input for gap-crossing and it is also true for sighted rats in darkness^30^. Our findings show that the activity of certain MEC neurons correlates with whisker movement (Figure 7 and Figures S15–16). Combined with the observed impact of whisker trimming on spatially modulated cells, this implies that S1 may play a role in spatial navigation. This interpretation is consistent with prior evidence that whisker-mediated tactile inputs influence hippocampal function and spatial navigation. Whisker deflection-evoked responses observed in hippocampal CA1 imply that tactile inputs are transmitted to this area via the somatosensory cortex, contributing to experience-dependent learning^66^. Permanent whisker removal reduces hippocampal neurogenesis and impairs spatial memory^67^. The hippocampal CA1 region responds to whisker stimulation, likely aiding the update of spatial cognitive map^27^. Additionally, place fields are modulated by tactile inputs and can be disrupted by whisker pad inactivation^5^. Given the strong hippocampus-MEC connectivity, these findings suggest a potential neural circuit linking S1 and MEC. Although direct monosynaptic projections from S1 to MEC have not yet been confirmed anatomically, several multisynaptic pathways may mediate this communication. For instance, the perirhinal and postrhinal cortices receive extensive somatosensory input and project heavily to MEC^68–70^. Additionally, thalamocortical circuits involving the posterior thalamic nucleus—known to process vibrissal information—may modulate MEC activity indirectly via these cortical areas^71^. Furthermore, the claustrum (Cla) provides the MEC with multisensory information, including information from the primary motor cortex (M1), primary visual cortex (V1), and S1, thereby potentially influencing its spatial-coding process^72^. This network of connections suggests that whisker-related information may reach MEC through multiple parallel and converging pathways. Future investigations should employ techniques with higher temporal resolution, and incorporate more complex behavioral paradigms to more precisely characterize MEC neuronal responses to tactile stimuli and address the limitations of the current study.

## Methods

### Animal subjects

Our study used adult male C57BL/6 wild-type mice. A total of 13 mice were used for the L-D-DW-LW experiment in the visual cue-enriched (Environment 1), and 10 mice were assigned to the tactile cue-enriched environment (Environment 2). In Environment 2, three mice underwent continuous calcium signal recordings for five consecutive days and two mice were subjected to a second round of whisker trimming after their whiskers had regrown to their original length following the initial trimming. Among the 23 mice that underwent the open-field experiments, 8 were also used in the whisker-related experiments. The Peking University Animal Use and Care Committee approved all procedures, which adhered to the guidelines set by the Association for Assessment and Accreditation of Laboratory Animal Care. The mice were individually housed under a 12-hour light/dark cycle at a temperature of 22°C, with unrestricted access to food and water. A running wheel was placed in the cage to enhance the mice’s activity. Behavioral experiments were all conducted during the dark phase. All mice were 9 to 11 weeks old at the time of surgery and were 12 to 24 weeks old when the formal experiment was conducted.

### Surgery

The surgical protocol was refined based on previous research (Low et al., 2014)^73^. The mouse was placed in a glass chamber with isoflurane to be temporarily anesthetized and then fixed at a stereotactic frame and was continuously anesthetized by inhaling isoflurane through a mask with 1.2% isoflurane flowing at 0.2 L/min. During the experiment, mouse was treated with an eye cream of erythromycin to prevent vision damage. An appropriate portion of scalp was resected to expose the skull’s dorsal surface and the tissue on the surface was cleaned up. A rectangle craniotomy measuring 2.0 mm in length and 1.4 mm in width was created over the right hemisphere, whose proximal edge centered 2.7 mm lateral to the midline and the distal edge was 4.0 mm away. The muscle near the craniotomy was carefully detached from the skull by surgery scissors. 200 nL of rAAV-hSyn-GCaMP6f-WPRE-hGh polyA/ AAV-hSyn-GCaMP6s-WPRE-hGh polyA was injected at a rate of 100 nL/min using a glass capillary (NANOLITER2020 Injector, World Precision Instruments, FL, USA), 3.1 mm lateral to the midline, 0.3 mm anterior to the superior margin of the transverse sinus and at a depth of 1.7 mm below the brain surface. The capillary was left in place for 10 minutes to let virus spread. Similarly, another 200 nL virus was injected 3.7 mm lateral to the midline, 0.3 mm anterior to the transverse sinus, and at a depth of 1.5 mm below the brain surface.

The interface of the brain and the cerebellum was found and a proper “pocket” between the brain and the cerebellum was created by a surgical forceps while the cortical dura remained intact. A prism with its custom-made holder was connected to a stereotactic micromanipulator (1760, Kopf, CA, USA) and tilted 10 degrees laterally to ensure the image surface of the prism was parallel to the brain’s dorsal surface. The prism assembly, consisting of a customized 1 mm diameter GRIN (gradient-index) lens attached to a 1 mm square prism (Grintech, Germany), was slowly inserted to the previously made “pocket” (subdural space within the transverse fissure). During implantation, the prism assembly was slowly lowered and moved in the anterior direction until the imaging plane of the prism was flush against the brain. Kwik-cast (World Precision Instruments, FL, USA) was used to seal the junction of the prism and exposed brain tissue. The scalp and the skull were adhered together through tissue adhesive (3M, MN, USA). Then, dental cement was used to cover the full skull and a head bar was firmly sticked.

## Experiment

### Training process

After recovering from surgery, the mice were kept in food-restriction regime, maintained at 80∼90% of their normal body weight and given adequate water. For Environment 1, t arena was an 80 × 80 cm^2^ white floor box, with three visual cue cards pasted on the three walls of the box. A camera was placed above the open-field to track animal’s movement and two LEDs were used to emit either a white light or infrared light. Before the formal experiment, mice were trained to get familiar with the experimental environment for about one week. During the first adaptation stage, the mice freely ran in the open-field with no burden on the head, the experimenter stood outside and randomly sprinkled cookie crumbs into the arena to attract mice to forage for food. After mice were accustomed to persistently running in the box and had a good trajectory coverage, the training moved to the next stage. During this stage, the mice ran in the box with the optical fiber connected to its baseplate on the head and a helium balloon was attached to the fiber in an appropriate place to balance the weight. After they got used to run with the fiber and gained good performance, the formal experiment began.

For mice in Environment 2, all training and calcium imaging recordings were performed under near-dark conditions to minimize visual cues. The white open field arena was similar in shape and size to that used in Environment 1 but contained no visual cue cards. To provide tactile stimuli, two vertical strips of white sandpaper with different grit sizes (approximately 15 cm in height) were attached to two walls of the arena, and a square piece of white sandpaper (∼25 × 25 cm²) was placed in one corner. The arena was positioned inside a behavioral chamber and enclosed with black blackout curtains.

### Open-field experiments

Calcium imaging was performed with the miniature two-photon microscope^35^ (LFMTPM-V2.0.1, Resolution: ∼1 mm; working distance: ∼1 mm, NA: 0.45 mm). Before imaging, the cover of the GRIN lens on the prism were removed, the gel was added to cover the surface of lens and then microscope was placed into the baseplate and locked with screws. GCaMP6f/GCaMP6s was excited at 920 nm, with the laser intensity ranging from 50 to 80 mW at the sample. Image acquisition was managed using GINKGO-MTPM (Transcend Vivoscope Biotech Co., Ltd, China). Images of 600 pixel × 512 pixel were captured at a resolution of ∼0.89 μm/pixel and a frequency of 9–9.5 Hz with a femtosecond fiber laser (TVS-FL-01, Transcend Vivoscope Biotech Co., Ltd, China). Running trajectory was recorded through the camera on the ceiling at 25 Hz, behavioral videos and calcium imaging videos were simultaneously acquired by software GINKGO-MTPM (Transcend Vivoscope Biotech Co., Ltd, China).

### Experiments in environment 1 (E1)

The formal Light-Dark-Dark, Whisker-trimmed-Light, Whisker-trimmed (L-D-DW-LW) was conducted continuously within one day. In the first session (L), neuron activity was recorded as the mouse freely explored the open field with the warm-light LED on, to establish a baseline performance. In the second session (D), the light was turned off to create a dark environment, testing the influence of visual input. After completing dark session, the mouse was head-fixed, and the experimenter quickly removed all its whiskers using scissors. The whisker-trimmed mouse was then placed back into its home cage to acclimate for a while before the subsequent experiments began. Once the mouse resumed normal activities (e.g., running on the wheel), it was then reintroduced into the open field for the third session (DW) exploration, where the light remained off, and tactile input mediated by whiskers was deprived. In the fourth session (LW), the light was turned on again to test whether the regained visual input could restore spatial cognition while tactile input was deprived. Ambient light intensity in the experimental environment was measured using a multi-source illuminance meter (DL333205; Deli Instruments, Ningbo, China; measurement range: 0 – 200,000 lux). During light conditions, illumination was maintained at 4.5 – 4.6 lux, whereas under dark conditions, light intensity was measured as 0.0 lux. The lengths of the longest macro vibrissae in the experimental mice ranged from approximately 2.6 to 3.1 cm prior to trimming. Each session lasted 25 to 40 minutes to ensure that the trajectory evenly covered the whole open field. Prior to formal recording, mice were placed in the open field for habituation and then removed. The walls and floor of the arena were thoroughly cleaned before the start of the experiment. For the three animals in which the probe was detached between sessions D and DW, gel was applied to the lens surface before reinsertion.

### Experiments in environment 2 (E2)

For the rotation experiment, six male mice were first allowed to establish a stable representation of the tactile cue-enriched open field under dark conditions. The mice were then returned to their home cages. The arena was thoroughly cleaned by removing all feces and food debris with a vacuum cleaner, followed by careful wiping of the sandpaper, the floor and walls with alcohol. In addition, alcohol was sprayed into the air to minimize potential residual olfactory cues. Subsequently, the entire arena—including the primary tactile cues (sandpaper) as well as any subtle visual features on the floor and walls—was rotated 90° clockwise. The mice were then reintroduced into the rotated environment for a second recording session.

To directly assess the impact of whisker trimming in the tactile cue-enriched environment, we conducted an additional Dark-Dark, Whisker-trimmed task using 10 mice. In this task, animals were first allowed to establish stable spatial maps during a baseline dark session (D), followed direct whisker trimming (DW). Two of the mice underwent a second whisker-trimming experiment after their whiskers had regrown to sufficient length, with imaging performed in a newly selected field of view. In addition, four mice in this cohort were tested in a DWR session following the DW session, in which the sandpaper attached to the floor was removed. The remaining six mice underwent a DWR’ session after the DW session, in which the arena walls (together with the attached sandpaper) were removed while the sandpaper on the floor was retained. All sessions for these six mice were conducted on a platform elevated approximately 20 cm above the ground.

### Whisker-related experiment

For “stimulated-whisking“ sessions, mice were fixed in a platform, the movement of whiskers were recorded through a camera directly above the mice’ head at about 45 Hz and neuron activity were simultaneously recorded by two-photon calcium imaging. A self-made automatic stimulator was placed on the side of the mouse’s cheek to enhance their whisker movement, which contained an automatic telescopic rod with a piece of sandpaper at the end. A “stimulated-whisking” session contained 20 continuous trials, each trial lasted for 1 minute, consisting of 5 seconds of stimulation and 55 seconds of non-stimulation. During stimulation period, the rod was extended so that the sandpaper sheet at the end could be detected by most of the mice’ whiskers and the mice showed a higher frequency of whisker movement (see Video S2). During the remaining non-stimulation period, the rod was in its retracted state and the whiskers had no contact with the sandpaper (see Video S1).

For “spontaneous-whisking” sessions, everything was the same as the “stimulated-whisking” session, except that no stimulus was presented to the mice, that is, the rod was always in its retracted state and no piece of sandpaper sheet could be accessed by whiskers. Mice showed less whisker movement in this session.

### Histology

After completing all the experiments, the animals were anesthetized with isoflurane and then perfused with saline followed by 4% paraformaldehyde (PFA). Skin and muscles were removed from the skull cap with surgical scissors. The skull cap was fixed in PFA for at least 24 hours, to allow for fixation of the brain and cerebellum with the implant site of prism undeformed. The brain was cryoprotected in a 30% sucrose solution for 24 hours the day before slicing and the remaining skull cap and the prism were carefully removed. Sagittal sections of 30 μm thickness were obtained using a cryostat (Leica).

The brain slices were stained with DAPI, and for some mice, anti-GFP staining was also performed. For GFP staining, brain slices in PBS were first washed with a solution containing 10% goat serum and 0.3% TX100 in PBS for 1 h. They were then incubated overnight with the primary antibody (anti-GFP chicken, Thermo Fisher Scientific, catalog # A-10262, RRID AB_2534023, diluted 1:1000 in the goat serum/TX100/PBS solution) at 4℃. After this, the slices were washed again with the PBS for three 5-minute intervals before being incubated with the secondary antibody (goat anti-chicken IgY (H+L) Alexa Fluor 488, Thermo Fisher Scientific, catalog # A-11039, RRID AB_2534096, diluted 1:1000 in the goat serum/TX100/PBS solution) at room temperature for two hours. Finally, the slices were washed twice for 5 minutes each in PBS at room temperature and then mounted with DAPI (Sigma Aldrich, catalog # D9542). The stained sections and prism implantation position were visualized using an Olympus VS120 virtual slide microscope or Zeiss Axio Scan Z1. Images were further processed by ZEN microscope software (Carl Zeiss, Jena, Germany) and ImageJ (National Institutes of Health, Bethesda, MD, USA).

## Data analysis

### Position tracking

DeepLabCut (version 2.2)^74,75^ was utilized for tracking animals’ position in the open-field. Six key points were labeled for each mouse: the nose tip, head center, left ear, right ear, mid-body, and tail base. The head direction was calculated based on the coordinates of the nose tip and head center. For each mouse, approximately 0.5% of behavioral videos from all recorded sessions were extracted to create the training dataset, resulting in around 600–1200 frames per model. A ResNet-50-based neural network ^76^ was employed with default parameters for 650,000 to 1,030,000 training iterations. All models achieved a test error of less than 4 pixels and a training error of less than 3 pixels (image resolution: 1008 × 964 pixels). Subsequently, a p-cutoff value of 0.6 was applied to condition the X and Y coordinates for further analysis. This trained network was then used to analyze videos from all the sessions in the open field experiments. After obtaining the analysis results from DeepLabCut, the predicted pose and skeletal structure were overlaid onto the corresponding frame, generating a new video file for researchers to visually inspect the model’s performance. Additionally, in the subsequent analysis, frames in which the mouse’s position was outside the open field arena or where the distance between the nose tip and head center exceeded 4 cm were excluded as outliers.

### Calculation of behavior variables

Head angular velocity was calculated as the change in head direction angle between consecutive frames, divided by the elapsed time between frames (rad/s). Head angular acceleration was subsequently derived as the change in angular velocity between consecutive frames, similarly divided by the time difference (rad/s²).

For all boundary-related analyses, the boundary region was specifically defined as the area extending 10 cm inwards from the walls of the square arena. This width was chosen as it is widely adopted in existing literature for investigating boundary-related neuronal activity^77–79^.

To explore the relationship between border cell firing and animal’s perception for environmental boundary, we analyzed the firing patterns of border cells specifically during epochs when the animal was located within the border zone, and examined how the discharge probability related to the animal’s head direction. Firing events were categorized as either *Event*_facing_ (calcium events occurring when the animal’s heading fell within a defined angular range directed toward the wall) or *Event*_away_ (events occurring when the animal faced away from the wall) (Supplementary Figure S6F). To account for potential behavioral biases in the animal’s head direction distribution within the border zone, event counts were normalized by the corresponding dwell time in each heading category: *Heading* _facing_ (total time spent facing toward the wall) and *Heading*_away_ (total time spent facing away from the wall). The inward and outward event rates were thus defined as:

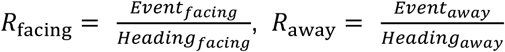

A firing preference index *C* was then computed as the ratio of inward-to-outward event rate:

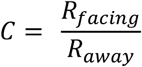

A value of *C* > 1 indicates that the cell fires at a higher rate when the animal faces toward the wall than away from it. Only cells for which both *Heading* _facing_ > 0 and *Heading*_away_ > 0 were included in subsequent analyses.

“Wall approaching time” was calculated to quantify the proportion of total recording frames during which the animal engaged in direct wall contact behavior. This was specifically defined as instances where the animal’s position was located with boundary area and its head was simultaneously oriented directly towards the wall (a head direction angle of ±45° relative to the wall normal).

### Calcium Signal Processing

Image stacks from all sessions recorded within a single day were processed using Suite2P^80^, which performed motion correction, region of interest (ROI) detection, and calcium signal extraction. The raw fluorescence signal (*F*_cell_) and the neuropil fluorescence signal (*F*_np_) were directly obtained from the Suite2P output file (Fall.mat) and further analyzed using a custom MATLAB pipeline. *F*_cell_ and *F*_np_ represent the weighted averages of fluorescence within and surrounding an ROI, excluding pixels overlapping with other ROIs. The fluorescence time course of each cell was calculated by averaging pixel intensities within the ROI and correcting for neuropil contamination, with the corrected signal defined as^81,82^:

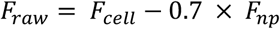

Baseline fluorescence (*F*_o_) was estimated using a moving-window method that smoothed the local 8^th^ percentile fluorescence and applied a correction term for stability. This method is consistent with the approach described by Zong et al. (2022)^82^. The normalized fluorescence change (*ΔF/F*) was then calculated as:

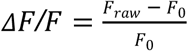

Artifacts were minimized by detecting and removing outliers based on z-scores. Wavelet-based denoising methods were applied to *ΔF /F* to further improve signal quality and obtain denoised *ΔF /F*. Significant transients were identified as periods during which denoised *ΔF/F* exceeded twice the local standard deviation for at least 0.75 seconds, following the criteria established by Zong et al. Denoised *ΔF / F* signals were then deconvolved using the Online Active Set method to Infer Spikes (OASIS) algorithm^83,84^, where the rise and decay time constants of calcium indicators were estimated automatically based on the fluorescence traces^84^. Frames containing significant transients were used to filter the raw deconvolved calcium activity (*E*_raw_), yielding the filtered signal ( *E*_filtered_). The filtered calcium activity was normalized, and non-zero events were classified as “calcium events”. The event rate was calculated as the number of calcium events per second for each cell. All subsequent spatial tuning analyses were performed using these deconvolved calcium events.

The signal-to-noise ratio (SNR) was determined as the ratio of the median amplitude of the 99^th^ percentile of *ΔF/F* during significant transients to the standard deviation of *ΔF/F* during resting periods.

### Calculation of tuning maps

For open-field experiments, the 80 × 80 cm^2^ square was divided into 2.5 × 2.5 cm² bins to create spatial tuning maps, which represent neuron’s activity normalized by occupancy. Each bin’s value in the tuning map was calculated as the ratio of the total calcium events to the duration spent in that bin. A Gaussian filter was applied to smooth position data. Arena coverage was defined as the proportion of visited bins to the total number of bins, with a bin considered visited if the animal spent at least 0.1 seconds there. To eliminate the effects of behavioral state changes (e.g., foraging versus resting), data points with running speeds below 2.5 cm/s or above 60 cm/s were excluded from all analyses of both observed and shuffled data.

Head-direction tuning maps and speed tuning curves were developed using similar methods (Range: 0–360° for head direction tuning curve and 2.5 cm/s – 18.5 cm/s for speed tuning function; bin size: 3° for head direction tuning curve, and 2 cm/s for speed tuning function). These values were then smoothed using a Gaussian filter (standard deviation: 2.5 cm for spatial heatmaps, 2° for head direction tuning curve, and 2 cm/s for speed tuning function).

Spatial information content, measured in bits per event, was determined using the formula^85^:

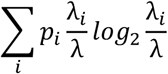

where λ_i_ is the mean event rate in the *i*-th location bin, λ is the overall mean rate, and *p*_i_ is the probability of the animal being in the *i*-th bin. Head information is calculated using the same method, except that *p*_i_ in this case is the probability of the animal’s head direction being in the *i*-th angle bin.

### Grid cells

All grid cells were selected from the baseline light session and assigned a grid score based on their autocorrelogram, as described by Sargolini et al.^14^ Briefly, each cell’s autocorrelogram was rotated at multiple angles, and Pearson correlations were calculated for rotations at 60° and 120° (Group 1) and at 30°, 90°, and 150° (Group-2). The minimum difference between any element in Group 1 and any element in Group 2 was determined. The grid score for each cell was defined as the highest minimum difference between Group 1 and Group 2 rotations across all successive circular samples. The central peak was not included in the analysis.

A cell was classified as a grid cell if its grid score exceeded a chance level determined by shuffling the experimental data 1,000 times. For each shuffle, the calcium event sequence of the cell was circularly time-shifted along the animal’s path by a random interval, following methods from Zong et al.^82^ Additionally, for each cell, we recalculated the spatial tuning map for the first and second halves of the baseline session and computed the correlation between these two maps to assess intra-session stability. A cell was defined as a grid cell if its recorded grid score exceeded the 95^th^ percentile of grid scores from the shuffled data and its intra-session stability was greater than 0.3 in baseline session. Besides, cells with a negative grid score were excluded from our data.

For detailed analysis of grid cells’ geometric features, we examined grid spacing and calculated ellipticity for each grid cell under each experimental conditional. Grid spacing was determined by measuring the distance from the center of each autocorrelogram to its six surrounding autocorrelation nodes, then averaging these distances. The standard deviation of these grid spacing values was calculated to reflect the regularity of grid pattern. To calculate ellipticity, an ellipse was fitted to the identified maxima using a least-squares method^86^. For cells where the fitting was successful, ellipticity (ε) was defined as the ratio of the semi-major (α) to semi-minor (β) axes of the fitted ellipse, yielding a value between 1 (a perfect circle) and infinity (∞). Only grid cells with an ellipticity value less than 5 in the baseline light session were included in the ellipticity-related analysis.

### Border cells

The border score was calculated to assess the spatial firing patterns of cells in relation to the environment’s boundaries, based on previously described methods^16^, with adjustments for calcium imaging data. Fields of a cell were identified using a threshold set at 30% of the maximum event rate within the map. The border score was then determined by calculating the difference between the highest coverage of any single field along any of the four walls and the average distance to the nearest wall for that field, weighted by event rate and normalized by half the shortest side of the environment. Border scores ranged from - 1 to +1, with a score of +1 indicating cells with infinitely narrow fields perfectly aligned along the entire wall.

Border cells were identified based on their border scores significantly surpassing chance levels determined by a shuffling procedure, similar to that used for grid cells. A cell was classified as a border cell if its true border score exceeded the 95^th^ percentile of scores from 1000 shuffled permutations of the same cell and the spatial correlation between the first and second halves of the baseline session was above 0.3.

### Spatial cells

Spatial cells were characterized by a spatial correlation greater than 0.5 between the two halves of the baseline light session and spatial information content exceeding the 99^th^ percentile of the shuffle distribution, excluding grid cells and border cells.

### Head direction cells

All HD cells were selected from the baseline light session. We calculated the mean resultant vector (mean vector length, MVL) from the head direction tuning curve following the method described by Sargolini et al^14^. A cell was considered to be modulated by head direction if its MVL exceeded a chance level established by repeatedly shuffling the experimental data for 1,000 times. For each time, the deconvolved calcium activity sequence was circularly time-shifted relative to the sequence of head-direction of the mice. We also assessed the intra-session tuning stability of HD cell candidates by calculating the Pearson correlation between the head direction tuning curves from the first and second halves of the session. A cell was identified as a HD cell if its MVL was larger than the 99th percentile of MVLs from its shuffled data and had an intra-session stability larger than 0.5. Head information was calculated as described above to measure the information about animals’ head direction conveyed by a HD cell. If a cell met the criteria for both HD cells and grid cells, it was classified as a Grid × HD cell; otherwise, it was categorized as a pure HD cell.

### Field extraction and ROI classification

Firing fields were identified from two-dimensional rate maps. A firing field was defined as a contiguous region comprising at least 30 spatial bins (2.5 × 2.5 cm² per bin) in which firing rates exceeded 20% of the cell’s peak rate. Candidate regions were retained only if their peak rate was ≥ 0.1 Hz.

The tactile cue region (region of interest, ROI) was defined a priori as a 25 × 25 cm^2^ area located in the upper-right quadrant of the arena. Cells were classified according to the centroid location of their baseline firing fields. At the cell level, a grid cell was designated as an ROI cell if at least one field centroid fell within the ROI, whereas spatial cells were classified based on the centroid of the largest-area field. At the field level, each baseline field was independently categorized as ROI or non-ROI according to its centroid position. For field in baseline dark session (D), the closest field in session DW within the same cell was identified, and spatial overlap was quantified as the proportion of baseline field pixels overlapping with the matched field. A field was considered disappeared if the overlap ratio was < 0.2. In three animals, the tactile floor cue in the baseline session was rotated 90° clockwise relative to the other subjects. To standardize the tactile reference frame across animals, behavioral trajectories, spike coordinates and spatial rate maps from these animals were rotated 90°counterclockwise prior to analysis.

### Position Decoding

Animal’s position was decoded through a Naïve Bayes decoder^87^ based on neural activity in a certain session. We modeled events from different cells as independent Poisson and adopted a uniform prior for all spatial bins^88^. Calcium event data was reassigned into intervals of 1 s and smoothed using a 5 s time window. The average position of the animal across all frames within each time window was used to represent the mouse’s position in that specific time bin. For each time bin, the probability of the animal being in a specific position bin was calculated using the formula:

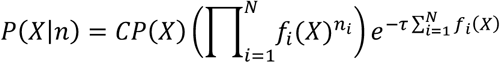

where *X* is the spatial bin, the element *n*_i_ of vector **n** = (*n*_1,_*n*_2_, …, *n*_N_) is the number of calcium events for neuron **i** within each time bin, *C* is a normalization factor, *P*(*X*) indicates the overall probability of the animal occupying spatial bin *X*, *f*_i_(*X*) is the mean event rate of cell **i** in spatial bin *X*, and *τ* is the length of the time window. While the vector *n* referred to testing data, other variables were derived from training data. The position with the maximum likelihood was identified as the animal’s decoded position for each time bin. The decoding error for test data was calculated as the mean absolute error (MAE) between the actual position and the decoded position.

For each mouse with *n* grid cells, we randomly selected *k* cells from the total grid cells for *k* ranging from 1 to *n* to establish decoder for baseline session. This process was repeated 100 times to obtain the MAE of the decoder with *k* cells. We observed that the MAE gradually decreased as *k* increased, indicating an improvement in the decoder’s performance (Figure S6D). Ultimately, we selected four mice with more than 20 grid cells each for subsequent statistical analysis related to decoding.

The first half of the baseline light session served as the training set, while the second half was used as the test set to establish a basic spatial-coding pattern when no sensory inputs were deprived. Subsequently, we decoded the animal’s position using the second half of the following sessions (D, DW, LW) as the test set, based on the initial training set in baseline session, to investigate whether the spatial-coding pattern of grid cells changed after sensory deprivation. A shuffled data set was created by randomly time-shifting the animal’s position in the baseline light session for 1,000 times.

For the decoding analysis based on all functional cell types, the neuron activity of HD cells, border cells, grid cells, and spatial cells was used, with the decoder construction method being the same as described above. The same procedure was used to construct the position decoder in Environment 2. The only difference was that the first half of the Dark baseline session was used for training, whereas the second half of the D, DW, DWR, and DWR’ sessions was used for testing.

### Population vector (PV) correlation

To assess the change of population coding of HD cells after sensory deprivation, we computed the population vector correlation between session pairs. The head direction was first divided into 120 angle bins (3° each). For each bin, we calculated the corresponding population vector (PV) by concatenating the mean event rates of all HD cells associated with that bin. Subsequently, the Pearson correlation between PV for every possible pair of angular bins was calculated for each mouse, resulting in a correlation matrix for each session. Each value within this matrix represents the correlation between the PV of two specific angular bins. We then computed the mean correlation across all mice for each bin pair, yielding an averaged correlation matrix for each session pair. These averaged matrices were then combined into a single comprehensive matrix, which provides a visual representation of the consistency in head direction tuning across different sessions and angular bins (Figure 1M). Additionally, we compiled the tuning curves of all HD cells into a 2-dimensional matrix for each mouse, referred to as the population vector matrix. The correlation between these PV matrices was computed across session pairs. The mean and SEM of the PV correlations for all mice (n = 13) are presented in Figure 1N, with color representing the correlation values.

For border cells. We rearranged the 2D rate map of each cell into a 1D row vector and combined the event rates from all cells. This resulted in a 2D population vector matrix for each mouse in each session. We subsequently computed the PV matrix correlation between session pairs for each mouse. By averaging these data across all mice, we assessed the changes in the population coding of border cell after sensory deprivation (Figure 2I).

### Quantifying session-to-session changes at the single-cell level

To assess session-to-session changes in HD tuning at the single-cell level (baseline session **i** vs. condition session *j*, e.g., L to D), we used a resampling-based paired framework applied independently to each recorded cell. For each cell, we defined the analysis window length as half of the total duration of session **i**, and included cells only when session *j* lasted at least as long as this window duration. We then performed 100 iterations in which we randomly sampled a contiguous time window of this fixed length from session **i** and an independent time window of the same duration from session *j*, extracting spikes and position samples within each window. To ensure data quality, sampled windows were accepted only if they contained at least 50 events; otherwise, we resampled up to 50 attempts per iteration, and iterations failing these criteria or yielding empty/invalid tuning estimates were treated as missing. For each accepted window, HD turning curves were computed and tuning statistics were extracted to quantify multiple properties of HD coding. Specifically, changes in tuning strength were computed as changes in MVL, directional drift across sessions was quantified as the absolute circular difference in PFD. In addition, tuning-curve similarity was assessed by comparing stability within session **i** (correlation between tuning curves computed from a randomly sampled window and its complementary time segment) against stability across sessions (correlation between the session **i** window and the matched-length window from session *j*). For all three metrics, statistical inference was performed independently within each cell using a paired one-tailed sign test across valid resampling iterations. For each cell, paired differences were computed between cross-session and within-session values, and the null hypothesis assumed that positive and negative differences were equally likely (Binomial distribution, p=0.5). The direction of the one-tailed test was defined according to the specific hypothesis for each metric. Specifically, for tuning strength (mean vector length, MVL) and tuning-curve similarity (correlation), we tested whether session *j* exhibited a significant reduction relative to the within-session baseline of session **i** (i.e., cross-session values < within-session values). In contrast, for PFD drift, we tested whether cross-session directional deviation was significantly greater than the within-session deviation observed in session **i**. Resulting p-values were corrected across cells using the Benjamini-Hochberg false discovery rate (FDR) procedure. Cells were classified as showing a significant change at q<0.01, additionally requiring that the median metric across iterations reflected the hypothesized direction of effect (i.e., median MVL or correlation decreased in session j, or median PFD deviation increased relative to within-session variability).

To quantify how many HD cells exhibited alterations in one or more aspects of tuning, each metric was binarized per cell (significant vs. not significant after FDR correction). Cells were then categorized according to multi-metric change patterns, and we reported the counts and proportions of cells showing a significant change in exactly one metric, in any two metrics, in all three metrics, or in none (Figures 1L and S10G).

A similar resampling-based framework and statistical logic were applied to grid cells, border cells, and spatial cells (Figures 2H,3H,5F,5H and S10J). In these analyses, spatial rate maps were computed for each resampled window, and corresponding spatial metrics were extracted, including grid score, border score, and spatial information, depending on cell type. For grid cells, we tested whether the grid score in session *j* was significantly reduced relative to session **i**, whether the cross-session spatial rate map correlation was significantly lower than the within-session correlation of session **i**, and whether the absolute displacement of the peak firing location between sessions exceeded the displacement observed between two within-session resampled segments. For spatial cells, the same procedures were applied, with spatial information replacing grid score as the tuning-strength metric. For border cells, we examined whether the border score was significantly reduced, whether spatial rate map correlation decreased, and whether the distance between the peak firing location and the nearest environmental boundary increased in session *j* relative to session **i**.

### LDA-based dimensional reduction

To investigate the structure of head-direction representation and its alteration under sensory deprivation, we applied linear discriminant analysis (LDA)^89^, a supervised dimensionality reduction technique. LDA seeks an optimal linear transformation that maximizes between-class variance while minimizing within-class variance. In our study, we used LDA-based method to project the high-dimensional neuronal activity on a 2-dimensional plane with best separation of head direction tuning.

First, we computed neural trajectories in an N-dimensional space using a binary matrix *A* ∈ ℝ^N×T^ to represent the calcium events, where *N* is the number of identified head-direction cells in the baseline session, and *T* is the number of time frames. Each matrix *A*_i,j_, indicate the present (1) or absence (0) of a calcium event for a neuron **i** at time *j*. We then divided the time frames into overlapping time bins, smoothing the data to reduce variance. Event rates of each neuron were calculated for each bin, and head-direction angles were assigned based on the direction nearest to the bin center, resulting in an N-dimensional neural trajectory matrix *B* ∈ ℝ^N×Tr^and a corresponding head-direction array *H* ∈ ℝ^Tr^. These were then subjected to LDA for dimensionality reduction.

LDA calculates two scatter matrices, *S*_W_(within-class) and *S*_B_(between-class), and finds the optimal linear transformation Θ^∗^ that maximizes the Fisher’s linear discriminant ratio

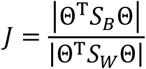

We used data from the baseline light session to derive Θ^∗^, which was then applied to project neuron activity from subsequent sensory deprivation sessions.

Intuitively, the Fisher’s linear discriminant ratio is an appropriate metric to assess the separability of data points in terms of their head directions. Additionally, we adopted the separation index

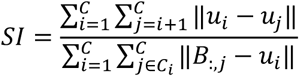

to cross-validate our findings on the deterioration of the separability of head directions caused by sensory deprivation.

When assess the intra-session stability of the representation of baseline light session (Figure S6B), Θ^∗^ derived from the baseline was used to separately project neural activity from the first and second halves of the session.

## Whisker-related analysis

### Identification of “whisker-responsive” cells

Videos recording whisker movement were analyzed by Facemap^90^ or DeepLabCut^74^, which yielded the position of the start and middle points of a selected whisker in each frame, also named as root and tail points. The root and tail positions were determined by comparing the standard deviations of the initial position data, with the position set having a higher standard deviation designated as the tail, and the other as the root. The mean positions for both the root and tail points were calculated to establish the initial vector (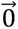) between these mean positions. For each frame, the vector (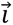) between the root and tail positions was computed, and the angle between 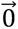 and 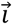 was calculated as angular position (*θ*). Following the angle calculation, the angles were smoothed using a Gaussian filter with a window size of 2s to obtain *θ*_smoo_. Angular velocity (*θ′*) was then calculated as the difference between consecutive smoothed angles. For each time in the neuronal activity data, the corresponding time in the behavioral data was identified by finding the closest matching time point. The mean absolute values of *θ′*over each interval between consecutive timestamps of calcium trace were then smoothed using a Gaussian filter with a window size of 1s to obtain the final value of whisker movement. The whisker movement sequences of three distinct whiskers from the same animal were highly similar, demonstrating the reliability of our method and suggesting that this whisker movement sequence can be effectively used in whisker-related neuronal activity studies.

A cell was classified as “whisker-responsive” based on the Pearson correlation between its denoised Δ F/F trace and the whisker angular velocity sequence. To assess significance, a shuffle procedure was performed 1,000 times. For each shuffle, the deconvolved calcium activity trace was randomly circularly shifted relative to the whisker movement, and the Pearson correlation was recalculated to generate a null distribution. To increase robustness, the “stimulated-whisking” session was divided into two equal halves, and the shuffle procedure was applied independently to each half. A cell was identified as “whisker-responsive” only if its true correlation exceeded the 95^th^ percentile of the shuffled correlation distribution in both halves of the session.

Calcium imaging data collected from the “free-exploration” session, “spontaneous-whisking” session, and “stimulated-whisking” session were analyzed together using Suite2P to identify the same cells across these sessions and assign spatial-coding identities based on their performance in the “free-exploration” session. The standards used for cell detection were consistent with those used in open-field experiments.

### Anatomical analysis

To determine whether whisker-responsive cells exhibited anatomical clustering, we applied a nearest-neighbor (NN) distance-based permutation analysis, similar to approaches previously used to assess spatial organization of functionally defined cell populations^41^. Briefly, the anatomical location of each neuron was defined by the centroid of its spatial footprint extracted from Suite2p segmentation.

We restricted the analysis to animals with at least 10 responsive cells per imaging plane to ensure reliable estimation of spatial statistics. For a given FOV, we computed the Euclidean distances between all pairs of responsive cells and quantified clustering using the mean distance to the k nearest neighbors (k = 1–5). For each neuron, the average distance to its k closest neighbors within the responsive population was calculated, and the global clustering statistic was defined as the mean of these per-cell nearest-neighbor distances. To evaluate whether the observed spatial organization deviated from chance, we performed a label-permutation test in which the anatomical positions of all recorded cells were held fixed while randomly assigning the “whisker-responsive” label to an equal number of cells drawn from the full population. For each value of k, 1,000 permutations were generated to construct a null distribution of the mean k-NN distance expected under spatial randomness with preserved cell density. The empirical p-value was calculated as the proportion of shuffled datasets in which the mean k-NN distance was less than or equal to the observed value (left-tailed test). Z-scores were additionally computed relative to the shuffle distribution.

Significant clustering was defined as a mean NN distance significantly smaller than that expected under label permutation, indicating that whisker-responsive cells were spatially closer to one another than predicted by chance given the overall cellular layout. Results were evaluated across multiple k values (1 – 5 NN) to ensure robustness to scale (Figure S15G). When pooling data across mice, we normalized the observed mean nearest-neighbor distance for each NN group size within each mouse to the mean of its corresponding shuffled-label distribution. Thus, values significantly below one indicates that whisker-responsive cells are spatially closer to one another than expected by chance (Fig. S15H).

## Data availability

Example data supporting this work are available in https://zenodo.org/records/15656068. The full dataset is available from the corresponding author upon reasonable request. Source data are provided with this paper.

## Code availability

Scripts supporting this work are available in https://github.com/Bluemooncell/Analysis-on-MEC-neural-Activity.

## Acknowledgments

This research was funded by the Qidong-SLS Innovation Fund (2023002029 and 2020001540; C.M.), the Space Brain Project of Lingang Laboratory (Grant No. LG-TKN-202204-01; C.M.), the Science Fund for Distinguished Young Scholars in Beijing (JQ23023; C.M.), the Chinese Government Foreign Expert Project for Emilio Kropff (G2022101003L; C.M.), and the PICT-2019-2596 grant from the Argentine Ministry of Science (E.K.). We are grateful to the National Center for Protein Sciences at Peking University in Beijing, China, for their support with imaging and histology. Thank Emilio Kropff, Xiang Zhang and Meng Zhong for their invaluable guidance on experiment techniques and data processing during the early stages of this project.

## Author contributions

C.M, J.T designed research, J.T, N.H and Y.H collected the data, J.T, W.S, Y.Z, X.Z, Y.L and Z.W performed the surgery, J.T, Y.Z, S.Y analyzed the data. C.M supervised the project, J.T and C.M. wrote the paper with input from all authors.

## Declaration of interests

The authors declare no competing interests.

## Supplementary information

Figures S1–S16. Videos S1–3.

---

**Supplementary Figure 1.**
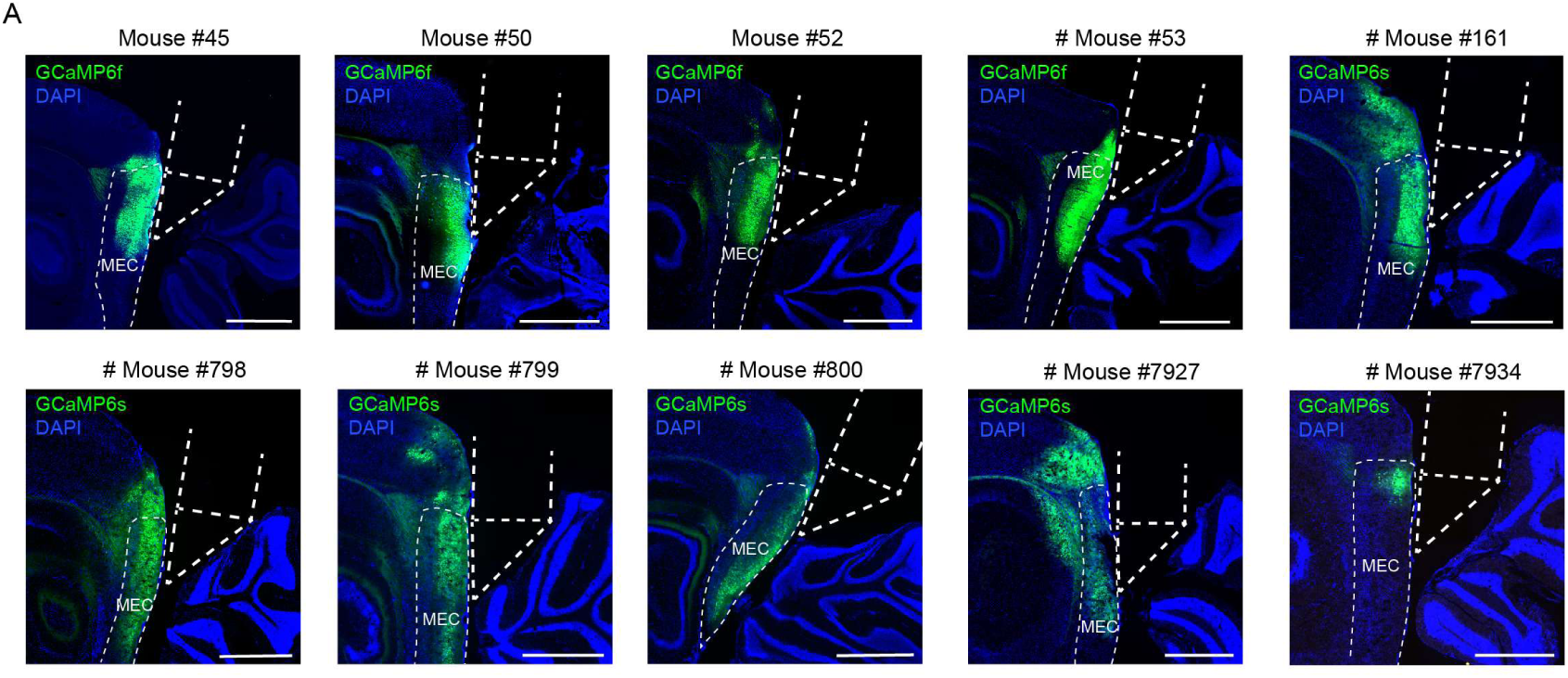
Histological reconstructions of MEC-targeted imaging. (A) Representative histological sections showing GCaMP6f or GCaMP6s expression (green) in the MEC across ten example mice. DAPI staining (blue) highlights cellular nuclei. A GRIN lens coupled to a microprism was implanted at the interface between the cortex and the cerebellum to facilitate optical access for imaging. The inferred position of the prism and the outline of MEC is indicated by white dashed lines. Scale bars represent 1 mm.

**Supplementary Figure 2.**
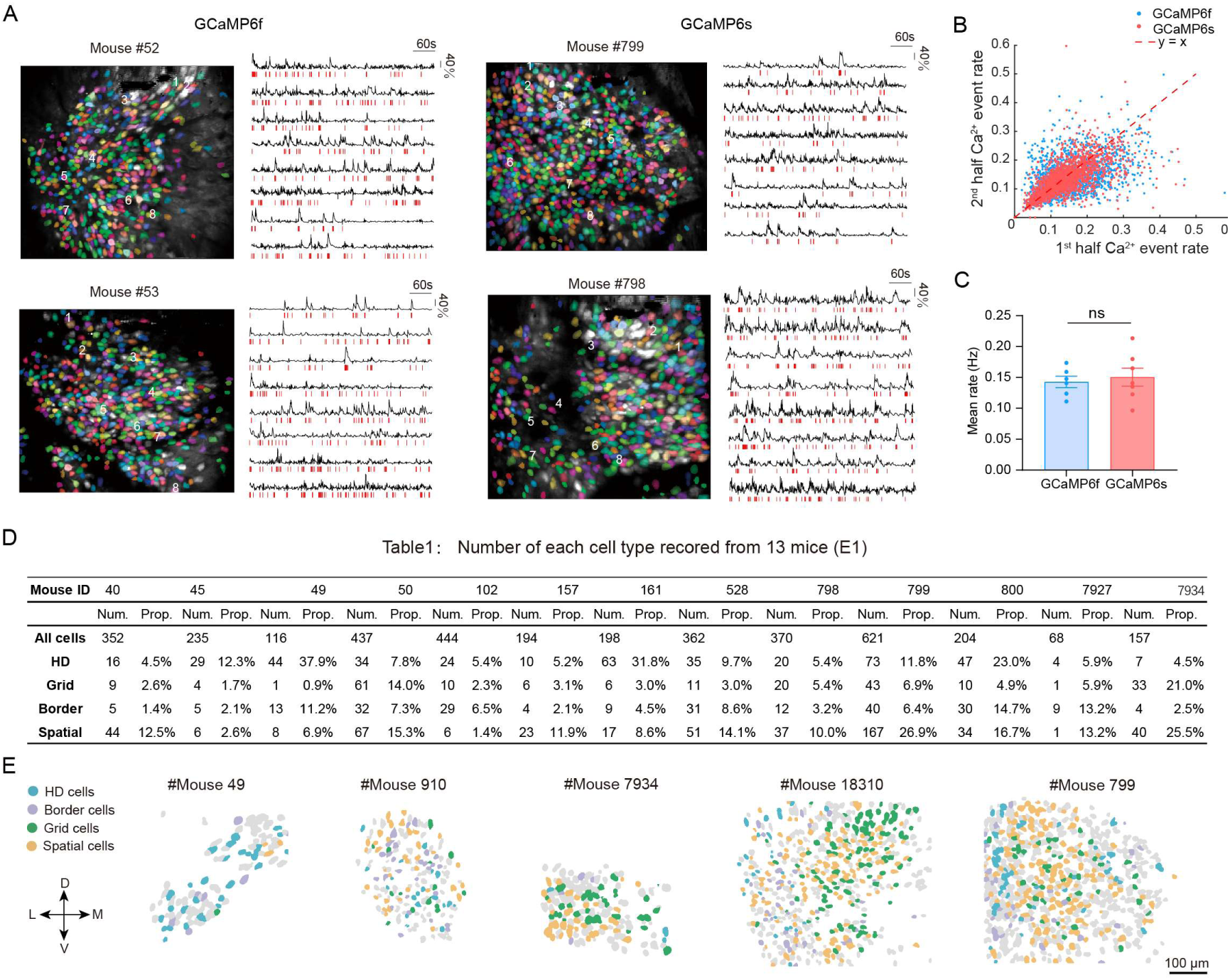
Distribution of spatially modulated cells across individual mice. (A) Example fields of view (FOVs) from four mice and the calcium signal traces of eight individual cells. The colored shapes represent the regions of interest (ROI) extracted and identified as cells using Suite2P. The black line represents the denoised ΔF/F trace, and the red vertical lines indicate the detected calcium events. The left column shows data from two mice injected with the GCaMP6f indicator, while the right column shows data from two mice injected with the GCaMP6s indicator. (B) Scatter plot depicting the distribution of event rates for cells expressing either the GCaMP6s or GCaMP6f indicator in the first and second halves of the experiments. (C) Bar chart showing no significant difference in the mean event rate across all sessions between two cell groups expressing the GCaMP6s and GCaMP6f indicators (Welch’s t-test, n1 =6 mice, n2 = 7 mice, p = 0.66). (D) Summary of the number and proportion of different functional cell types (HD, Grid, Border and Spatial) recorded from 13 mice in E1 (visual cue-dominated environment). The table shows the number of cells recorded (Num.) and the proportion of each cell type (Prop.). (E) Representative spatial footprints from several mice illustrating the anatomical distribution of different spatially modulated cells. Priority: Grid > Border > Spatial cells, HD cells are not excluded from these three spatial-coding cell types. ****P ≤ 0.0001; ***P ≤ 0.001; **P ≤ 0.01; *P ≤ 0.05; ns, P > 0.05.

**Supplementary Figure 3.**
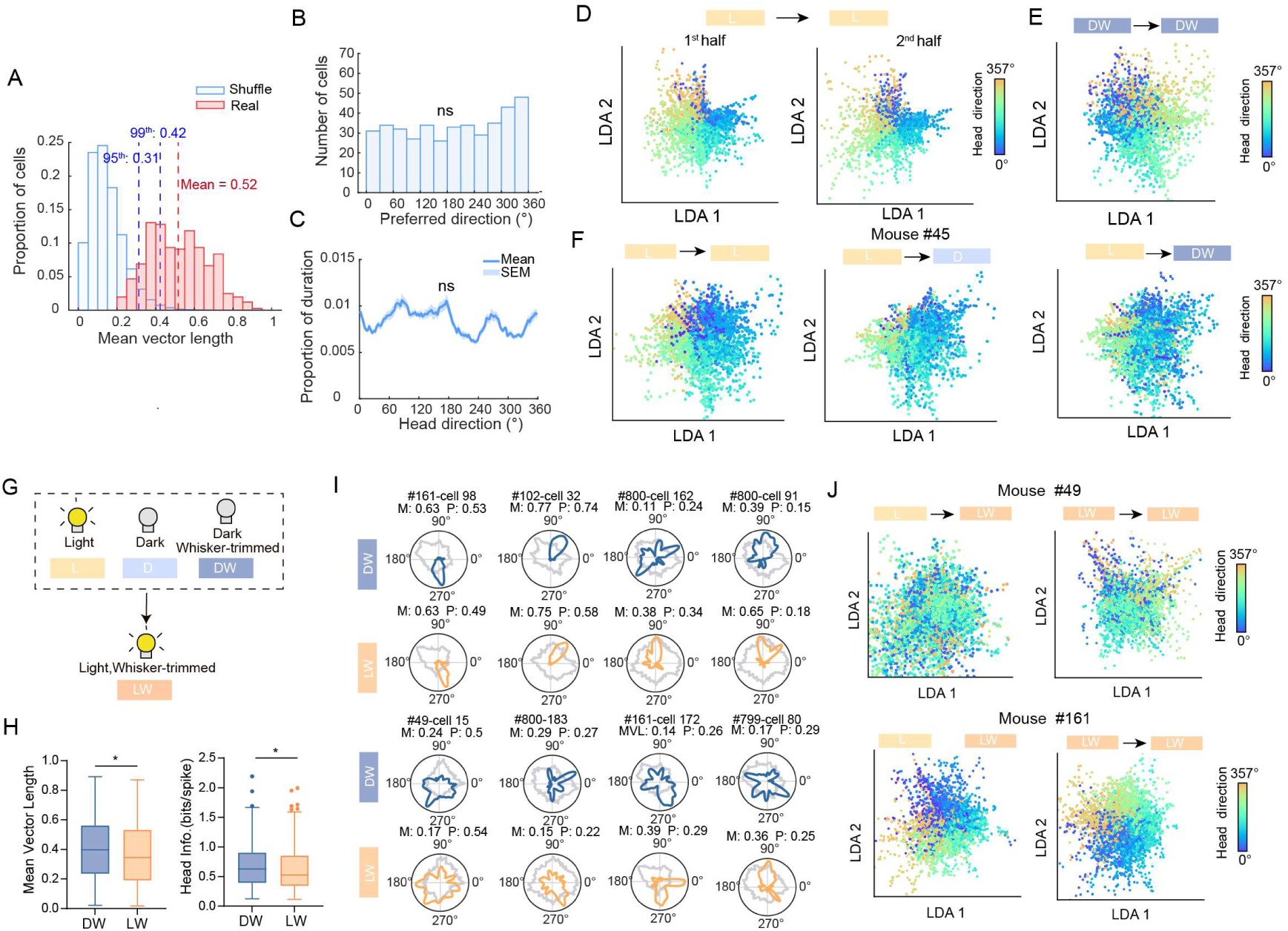
Tuning properties of HD cells were modulated by sensory inputs. (A) Distribution of mean vector length (MVL) for all HD cells during the light baseline session (L). The true distribution (red) is compared with a shuffle distribution (blue) generated by circularly shifting spike times relative to head direction. Dashed blue lines indicate the 95th and 99th percentiles of the shuffle distribution. (B) Histogram of preferred firing directions (PFDs) for all HD cells during the light baseline session, showing near-uniform coverage across the full 0-360° range (Rayleigh test: mean resultant length R = 0.078, Z = 2.47, P = 0.084). (C) Angular distribution of head direction sampled by animals during the light baseline session across all mice (mean ± SEM; bin size = 3°). No significant directional bias in behavior was observed (Rayleigh test: R = 0.066, Z = 0.53, P = 0.59). (D) Linear discriminant analysis (LDA) projections of HD population activity within the light baseline session (L). Left, activity from the first half of the session projected onto LDA axes trained on the whole session (L → L, first half). Right, activity from the second half projected onto LDA axes trained on the whole session (L → L, second half). (E) LDA projection of HD population activity in darkness with whisker-trimmed (DW), using LDA axes trained on DW itself. (F) Example LDA projections from Mouse #45 across experimental stages. Left: L → L, LDA trained and projected within the light session. Middle: L → D, neuronal activity in session D projected onto L-trained axes. Right: L → DW, neuronal activity in DW projected onto L-trained axes, revealing progressive drift of HD population representation across sensory deprivation. (G) Visual cues were reintroduced by turning on the light after the animals’ whiskers had been trimmed (Light/Whisker-trimmed, LW). (H) Left, box plot showing a decrease in MVL of HD cells after reintroduction of visual cues (Wilcoxon signed-rank test, n = 406, P = 0.032). Right, box plot showing a decrease in head information (bits/spike) of HD cells (Wilcoxon signed-rank test, n=406, P=0.044). (I) Representative tuning curves of individual HD cells in DW and LW sessions. Some cells exhibit partial or near-complete recovery of tuning, whereas others show no recovery. For each cell, MVL (M) and peak rate (P) are indicated. (J) LDA projections of HD population activity from two additional mice (#49 and #161). For each mouse, LW-session activity is projected onto LDA spaces trained on baseline L session (L → LW) or LW itself (LW → LW). ****P < 0.0001; ***P < 0.001; **P < 0.01; *P < 0.01; ns, P > 0.05.

**Supplementary Figure 4.**
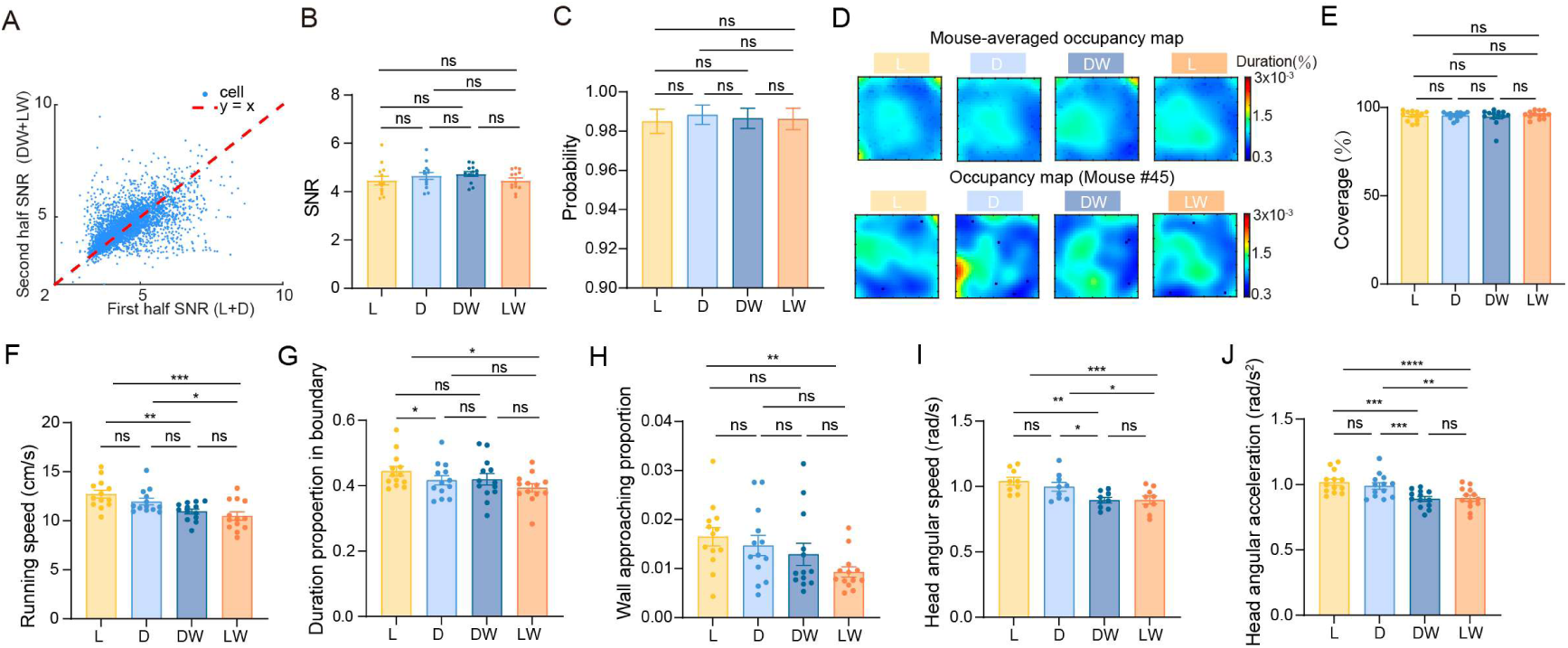
Cross-session imaging quality and animal behavior were not affected by sensory deprivation. (A) Scatter plot of signal-to-noise ratio (SNR) for all cells in the first half (L + D) versus the second half (DW + LW) of the sessions, with data points distributed around the line y = x (n = 3,758). (B) SNR across different sessions remained stable, indicating consistent imaging quality (RM-one-way ANOVA with Geisser-Greenhouse correction, n = 13, p = 0.22; post-hoc Tukey’s multiple comparisons test). Unless otherwise stated, all repeated-measures one-way ANOVA analyses in this study were performed with Geisser-Greenhouse correction and Tukey’s multiple comparisons test. (C) Bar chart shows no significant differences in the accuracy of DLC prediction results across sessions (RM one-way ANOVA, n = 13, P = 0.21). (D) Animal movement trajectory occupancy maps, showing the distribution of time spent by all mice (top) and a specific mouse (bottom). The color intensity represents the duration of time spent in each spatial bin, with red indicating high duration and blue indicating low duration. (E) Bar chart shows a consistency of the trajectory coverage in the open-field environment across sessions (RM one-way ANOVA, n = 13, P = 0.47). (F) Bar charts show no significant differences between adjacent sessions, but reveal a gradual decrease in average running speed across sessions, possibly reflecting progressive fatigue. (RM one-way ANOVA, n = 13, L/D: P = 0.065; D/DW: P = 0.11; DW/LW: P = 0.61; L/DW: P = 0.0059; D/LW: P = 0.023; L/LW: P = 0.00014). (G) The proportion of time spent within the boundary area (boundary width = 10 cm) showed a significant reduction only after the transition to darkness. (RM one-way ANOVA, n = 13, L/D: P = 0.010; D/DW: P = 0.97; DW/LW: P = 0.45; L/DW: P = 0.31; D/LW: P = 0.49; L1LW: P = 0.015). (H) Wall-approaching behavior declined in several sessions following whisker trimming. The wall-approaching proportion was defined as the fraction of total session time during which animals explored within the boundary area (Boundary width = 10 cm) while facing the wall. (RM one-way ANOVA, n = 13, L/D: P = 0.80; D/DW: P = 0.62; DW/LW: P = 0.28; L/DW: P = 0.0030; D/LW: P = 0.013; L/LW: P = 0.00018). (I) Head angular speed significantly decreases after whisker-trimming when animals explore the dark condition (D/DW) (RM one-way ANOVA, n = 13, L/D: P = 0.24; D/DW: P = 0.022; DW/LW: P > 0.99; L/DW: P = 0.0030; D/LW: P = 0.013; L/LW: P = 0.00013). (J) Head angular acceleration significantly decreases after whisker-trimming (RM one-way ANOVA, n = 13, L/D: P = 0.30; D/DW: P = 0.00098; DW/LW: P = 0.99; L/DW: P = 0.00025; D/LW: P = 0.0012; L/LW: P = 2.2 × 10^-5^). ****P < 0.0001; ***P < 0.001; **P < 0.01; *P < 0.01; ns, P > 0.05.

**Supplementary Figure 5.**
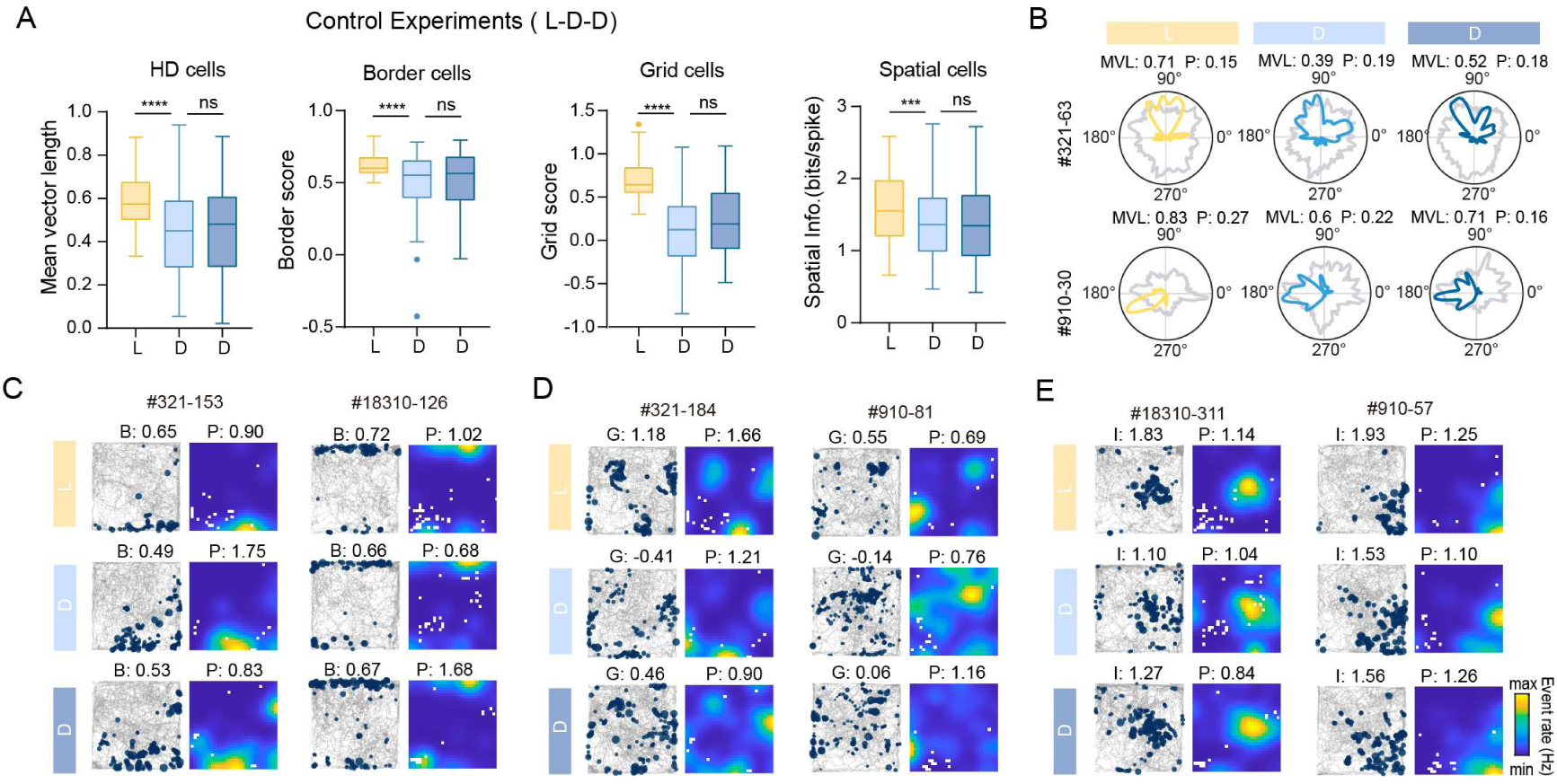
Tuning properties do not further deteriorate during prolonged darkness in the absence of whisker trimming. (A) In control experiments, mice underwent a second dark session without whisker trimming, No significant decline in the tuning of any spatially modulated cell types was observed. (Wilcoxon signed-rank test, MVL : n = 72, L/D:P = 2.0 × 10^-9^,D/D: P = 0.61; head information: n = 72, L/D: P = 2.8 × 10^-7^, D/D: P = 0.61; border score: n = 67, L/D: P = 3.8 × 10^-5^, D/D: P = 0.58; grid score: n = 64, L/D: P = 5.7 × 10^-10^, D/D: P = 0.25; spatial information of spatial cells, n = 182, L/D: P = 3.3 × 10^-7^, D/D: P = 0.42). (B) Representative tuning curves from two example HD cells across L, D, and the second dark session (D). MVL and peak rate (P) are indicated for each session. (C) Representative examples of border cells, showing spike locations (left) and firing rate maps (right). Border score (B) and peak rate (P) are indicated. (D) Representative examples of grid cells. Grid score (G) and peak rate (P) are indicated. (E) Representative examples of spatial cells. Spatial information (I) and peak rate (P) are indicated. ****P < 0.0001; ***P < 0.001; **P < 0.01; *P < 0.01; ns, P > 0.05.

**Supplementary Figure 6.**
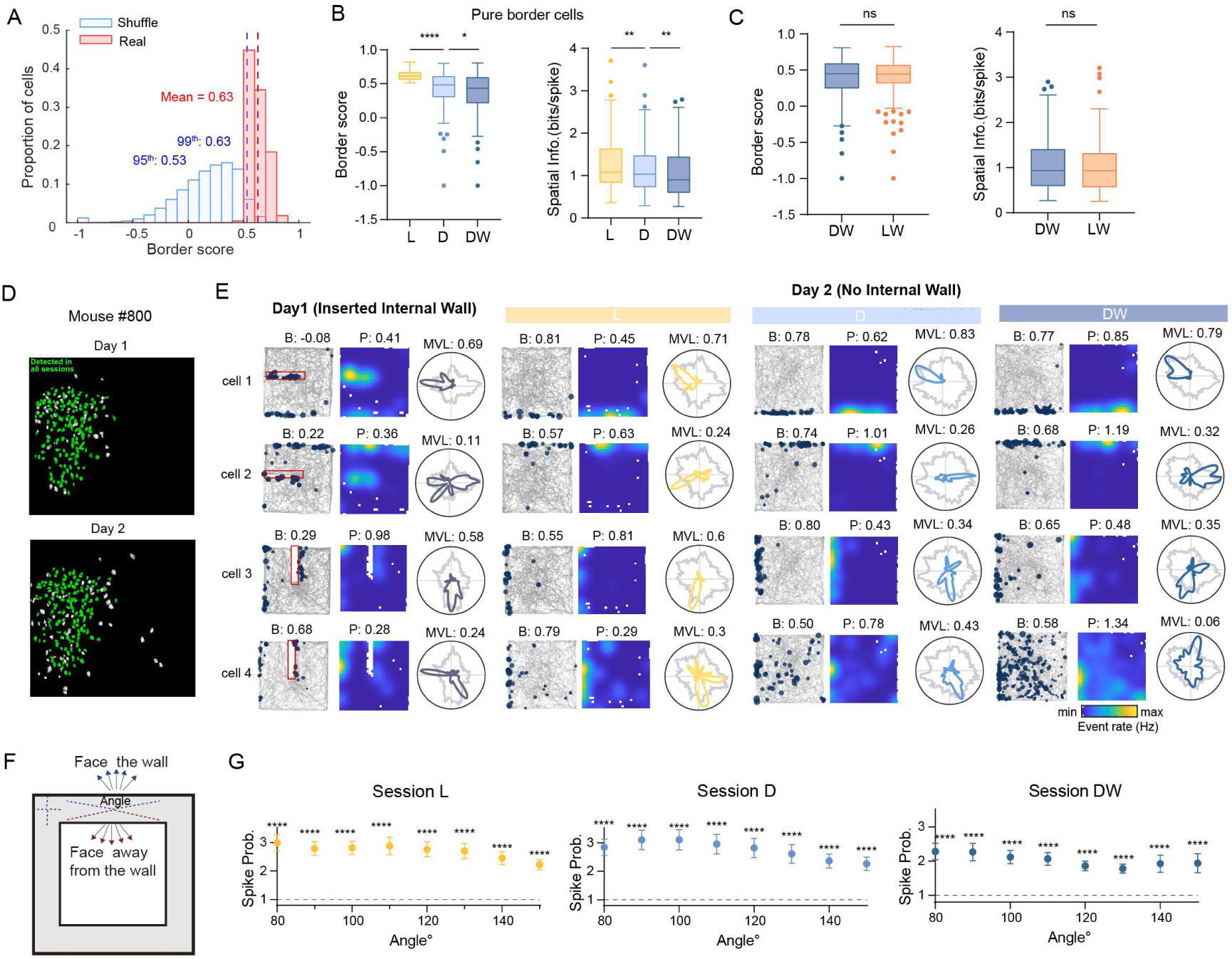
Border cell representations are shaped by both visual cues and tactile inputs. (A) Distribution of border scores for identified border cells. The blue histograms represent the distribution of shuffled data across all cells from all mice, while the red histograms show the distribution of true scores for the identified Border cells. The blue dashed lines indicate the 95th and 99th percentiles of the shuffled data, and the red dashed line indicates the mean score of the identified cell groups. (B) Border cells without head-direction tuning (Pure border cells) also show degraded performance following both light deprivation and whisker trimming (Wilcoxon signed-rank test, n = 185, L/D: border score: P = 8.0 × 10^-21^,spatial Information: P = 0.003; D/DW: border score: P = 0.0046, spatial information: P = 0.0059). (C) Population-level border score and spatial information do not significantly recover upon re-exposure to light (Wilcoxon signed-rank test, n = 223, border score: P = 0.48; spatial information: P = 0.98). (D) Example spatial footprints of MEC neurons imaged in the same field of view on Day 1 (with an inserted internal wall) and Day 2 (performing whisker-trimming manipulations), showing stable cell registration across days. (E) Representative rate maps and head-direction tuning curves of border cells that developed new firing fields adjacent to the inserted internal wall on Day 1. For each cell, the spatial firing rate map, head-direction tuning curve, border score (B), peak firing rate (P), and MVL are shown. (F) Schematic illustration of the analysis for wall-related head-direction modulation in border cells. Spikes and positional samples within a 10-cm-wide boundary region were extracted. For a given angular window, events and head-direction angles were separated into periods when the animal faced toward the wall or faced away from the wall, enabling computation of the spike probability ratio (face wall / face away). (G) Population-averaged spike probability ratio (face wall / face away) of border cells as a function of head-wall angle across Light (L), Dark (D), and Dark with whisker trimming (DW) sessions. In all sessions and across all tested angles, the spike probability ratio was significantly greater than 1, indicating a robust preference for firing when the animal faced the wall (one-sample Wilcoxon signed-rank test vs. 1, P < 0.0001 for all angles and sessions). **** P ≤ 0.0001; *** P ≤ 0.001; ** P ≤ 0.01; * P ≤ 0.05; ns, P > 0.05.

**Supplementary Figure 7.**
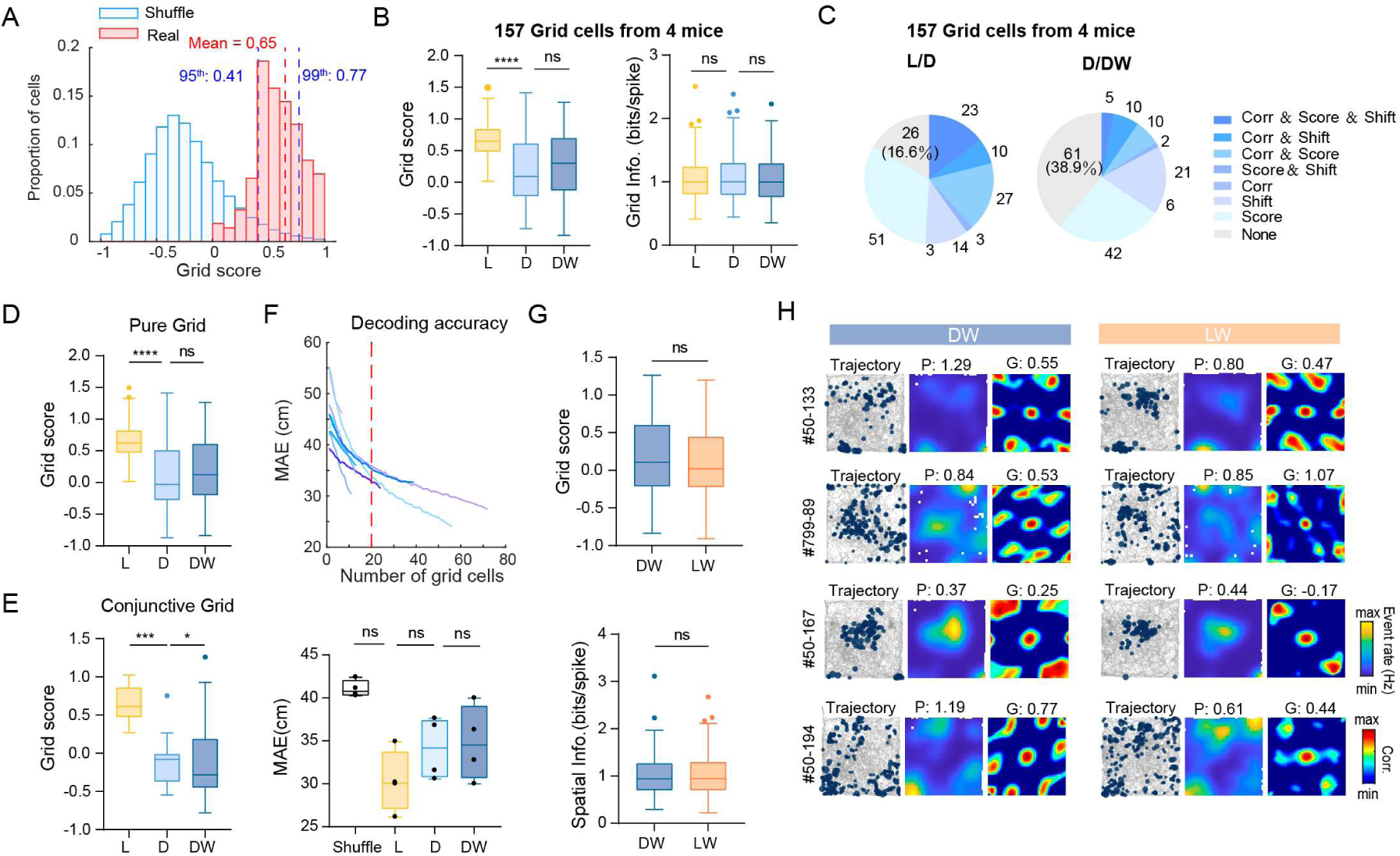
Spatial representations of grid cells are more influenced by light deprivation. (A) Distribution of grid score for all identified grid cells compared with shuffle level. The vertical dashed lines indicate the 95th and 99th percentiles of the shuffle distribution, and the red histogram shows the real data. (B) Box plots showing population grid score (left) and spatial information (right) of grid cells across Light (L), Dark (D), and Dark with whisker trimming (DW) sessions, computed from four mice in which grid cells were sufficiently abundant (>20 grid cells per field of view). Grid score showed a pronounced reduction from L to D, whereas spatial information was relatively preserved across conditions (Wilcoxon signed-rank test; n = 157, grid score: L/D, P = 9.0 × 10^-17^; D/DW, P = 0.060; spatial information: L/D, P = 0.88; D/DW: P = 0.27). (C) Pie charts summarizing the proportions of grid cells exhibiting significant changes in rate map correlation, grid score, and/or peak firing location shift between session pairs similar in Fig.3 (H). This analysis was restricted to the same four mice with grid cell-enriched fields of view (>20 grid cells). (D) Box plots confirming a decrease in grid score following light deprivation (L/D) and no additional reduction after whisker trimming (D/DW) in pure grid cells (Wilcoxon signed-rank test, n = 199, L/D: P = 1.2 × 10^-24^; D/DW: P = 0.040). (E) Box plot showing a strong decrease in grid score after light deprivation and a modest increase after whisker trimming in conjunctive grid cells (Wilcoxon signed-rank test, n = 16, grid score: L/D, P = 0.00053; D/DW, P = 0.92). (F) Top, decoding accuracy improves with an increasing number of grid cells. For decoding analyses based on grid cells, only four mice with more than 20 grid cells were used. Bottom, a linear decoder was trained using the neuronal activity of grid cells in the first half of the light baseline session, and tested on the second halves of the L, D, and DW sessions, respectively (Wilcoxon signed-rank test, n = 4 mice, Shuffle/L: P = 0.13; L/D: P = 0.13; D/DW: P = 0.63). (G) Box plots showing no significant recovery in grid scores and spatial information upon re-exposure to light after whisker trimming (DW/LW) (Wilcoxon signed-rank test, n = 215, grid score: P = 0.53; spatial information: P = 0.30). (H) Representative examples of grid cells recorded in DW and LW sessions. Some grid cells exhibited partial recovery of regular firing pattern upon reintroduction of visual cues, whereas others showed little or no recovery, highlighting pronounced heterogeneity in grid cell responses. For each cell, the trajectory, firing rate map, spatial autocorrelogram, peak rate (P), and grid score (G) are shown. **** P ≤ 0.0001; *** P ≤ 0.001; ** P ≤ 0.01; * P ≤ 0.05; ns, P > 0.05.

**Supplementary Figure 8.**
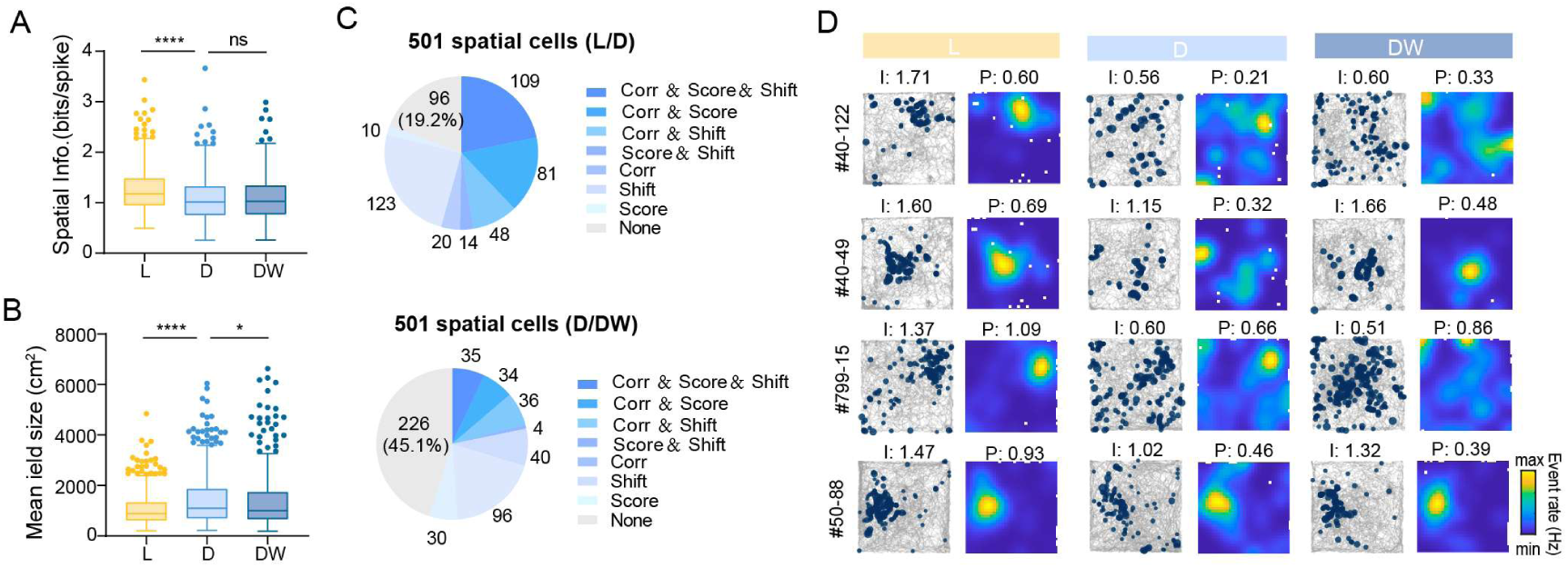
Spatial representations of MEC spatial cells are more influenced by light deprivation. (A) Box plots showing spatial information (bits/spike) of spatial cells across Light (L), Dark (D), and Dark with whisker trimming (DW) sessions. Spatial information is significantly reduced from L to D, but shows no further decrease from D to DW (Wilcoxon signed-rank test, n = 501, L/D: P = 3.4 × 10^-23^; D/DW: P = 0.27). (B) Box plots showing the spatial field size of spatial cells across L, D, and DW sessions. Field size significantly increases after light deprivation and decreases following whisker trimming (Wilcoxon signed-rank test, L/D: n = 497 pairs, P = 9.9 × 10^-14^; D/DW: n = 497 pairs, P = 0.026). (C) Pie charts summarizing the proportions of spatial cells exhibiting significant changes in multiple tuning metrics from L to D (top) and D to DW (bottom). Metrics include rate map correlation (Corr), change of spatial information (Score) and shift of peak rate position (Shift). Cells were classified as showing no significant change, single-metric changes, combinations of two metrics, or concurrent changes in all three metrics. (D) Representative examples of spatial cells across L, D, and DW sessions. For each cell, the animal’s trajectory (gray) and spatial rate map are shown. Spatial information (I) and peak rate (P) are indicated above each rate map. ****P ≤ 0.0001; ***P ≤ 0.001; **P ≤ 0.01; *P ≤ 0.05; ns, P > 0.05.

**Supplementary Figure 9.**
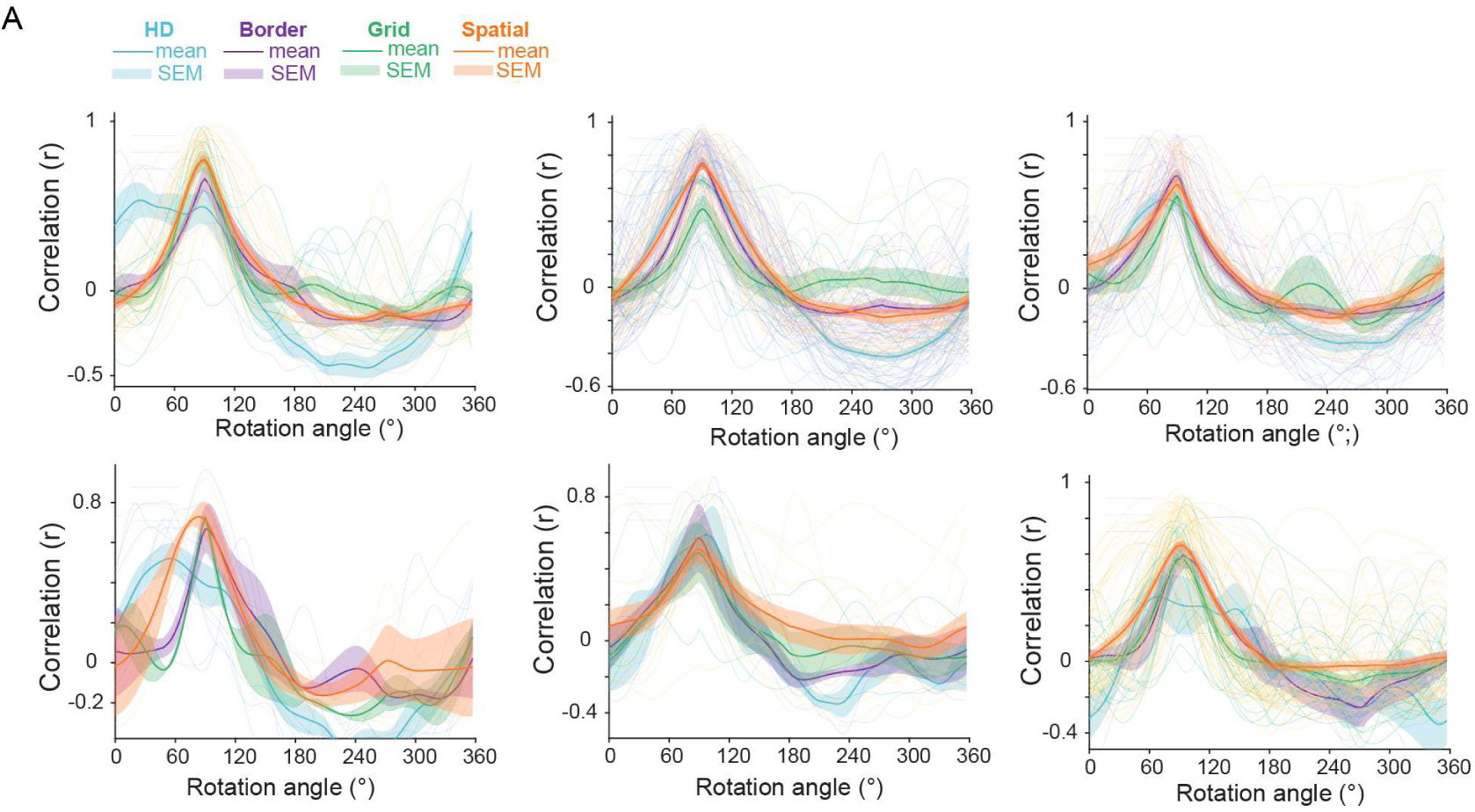
Individual rotation profiles of spatial tuning curves for six mice. (A) Rotation profiles of tuning curves for four classes of spatially modulated cells (HD, border, grid, and spatial cells) between Sessions A and B in the rotation experiments. The figure shows the rotation profiles for each of the six mice included in the rotation experiments. For each cell, the tuning curve from Session A was circularly rotated in 3° increments, and the Pearson correlation between the rotated Session A tuning curve and the Session B tuning curve was computed at each rotation angle. Thin background traces represent individual cells, whereas thick traces denote the population mean for each cell type; shaded areas indicate SEM.

**Supplementary Figure 11.**
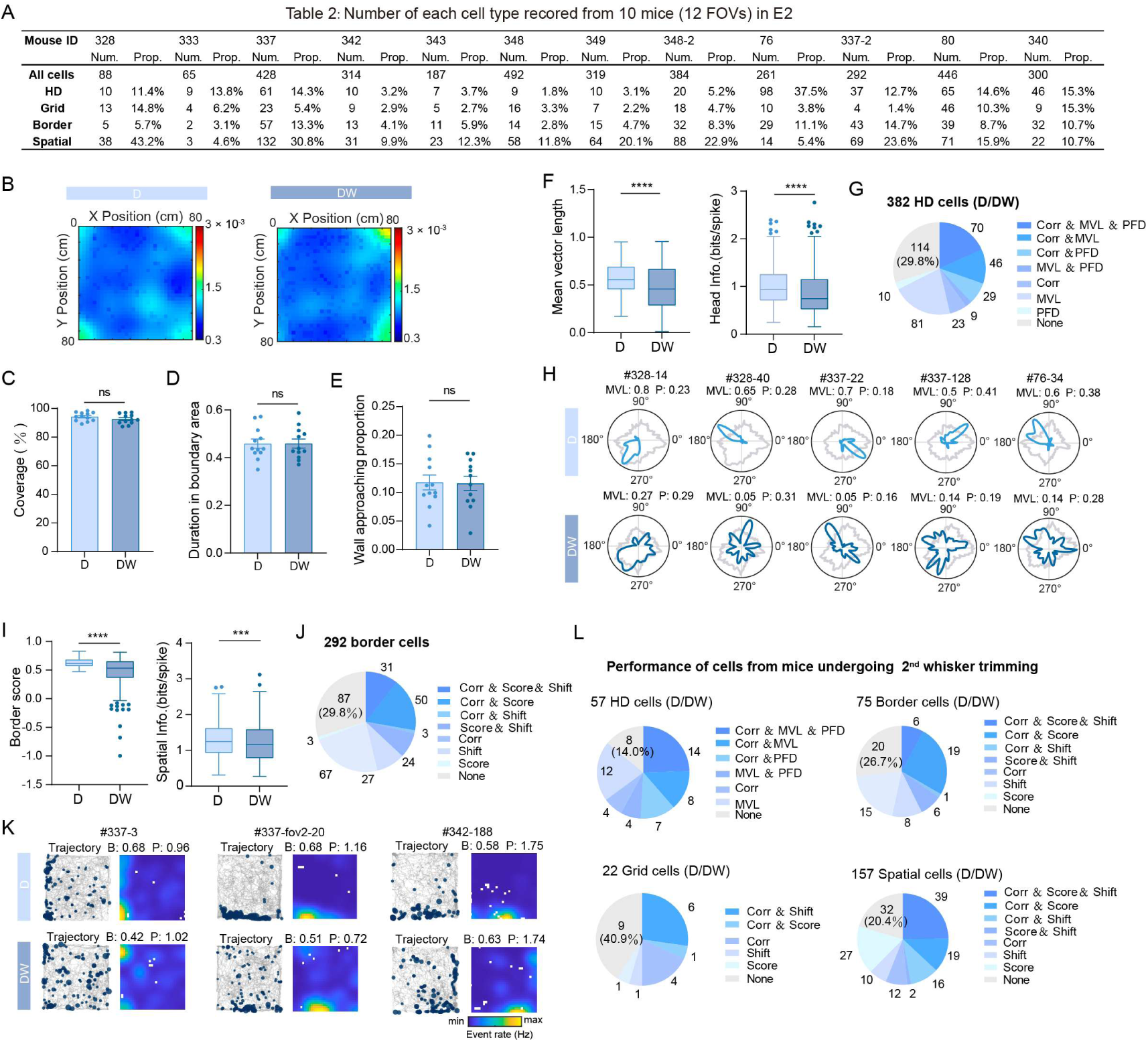
Responses of grid cells under different conditions suggest a predominant influence of visual input. (A) Summary table showing the number and proportion of each spatially modulated cell type recorded from 10 mice (12 FOVs) in the tactile cue-enriched environment (E2). (B) Averaged occupancy maps of animal trajectories across mice during session D (Dark) and DW (Dark, whisker-trimmed). (C) Bar plot showing no significant difference of arena coverage between D and DW (Wilcoxon signed-rank test, n = 12 mice, P = 0.06). (D) Bar plot showing no significant difference in the fraction of time spent in the boundary area (border width = 10 cm) between D and DW (Wilcoxon signed-rank test, n = 12 mice, P = 0.97). (E) Bar plot showing no significant difference of the proportion of wall-approaching behavior from D to DW (Wilcoxon signed-rank test, n = 12 mice, P = 0.62). (F) Box plots showing a significant decrease in tuning strength of HD cells after whisker trimming, quantified by MVL (Wilcoxon signed-rank test, n = 382, P = 2.3 × 10^-24^) and head information (bits/spike) (Wilcoxon signed-rank test, n = 382, P = 9.8 × 10^-15^). (G) Pie chart summarizing the proportions of HD cells exhibiting significant changes across sessions, including reduced tuning curve correlation (Corr), decreased MVL and/or drifted PFD (Shift). (H) Representative examples of HD tuning curves in D and DW. MVL and peak rate (P) are indicated. (I) Box plots showing a significant decrease in border score of border cells after whisker trimming (Wilcoxon test, n = 292, P = 2.63 × 10^-23^) and a significant reduction in spatial information (bits/spike) (Wilcoxon signed-rank test, n = 292, P = 0.0060). (J) Pie chart summarizing the proportions of border cells exhibiting significant changes across sessions, including decreases in rate map correlation (Corr), border score, and/or increases in the location shift of peak firing (Shift). (K) Representative examples of firing maps of border cels in D and DW. Border score (B) and peak rate (P) are indicated. (L) Pie charts summarizing the proportions of cells showing significant changes in multiple tuning metrics during a second whisker-trimming exposure, recorded from two mice after whisker regrowth in a newly imaged field of view (the second FOV). Pie charts are shown for four spatially modulated cell types (HD, border, grid, and spatial cells; cell numbers indicated above each chart). **** P ≤ 0.0001; *** P ≤ 0.001; ** P ≤ 0.01; * P ≤ 0.05; ns, P > 0.05.

**Supplementary Figure 11.**
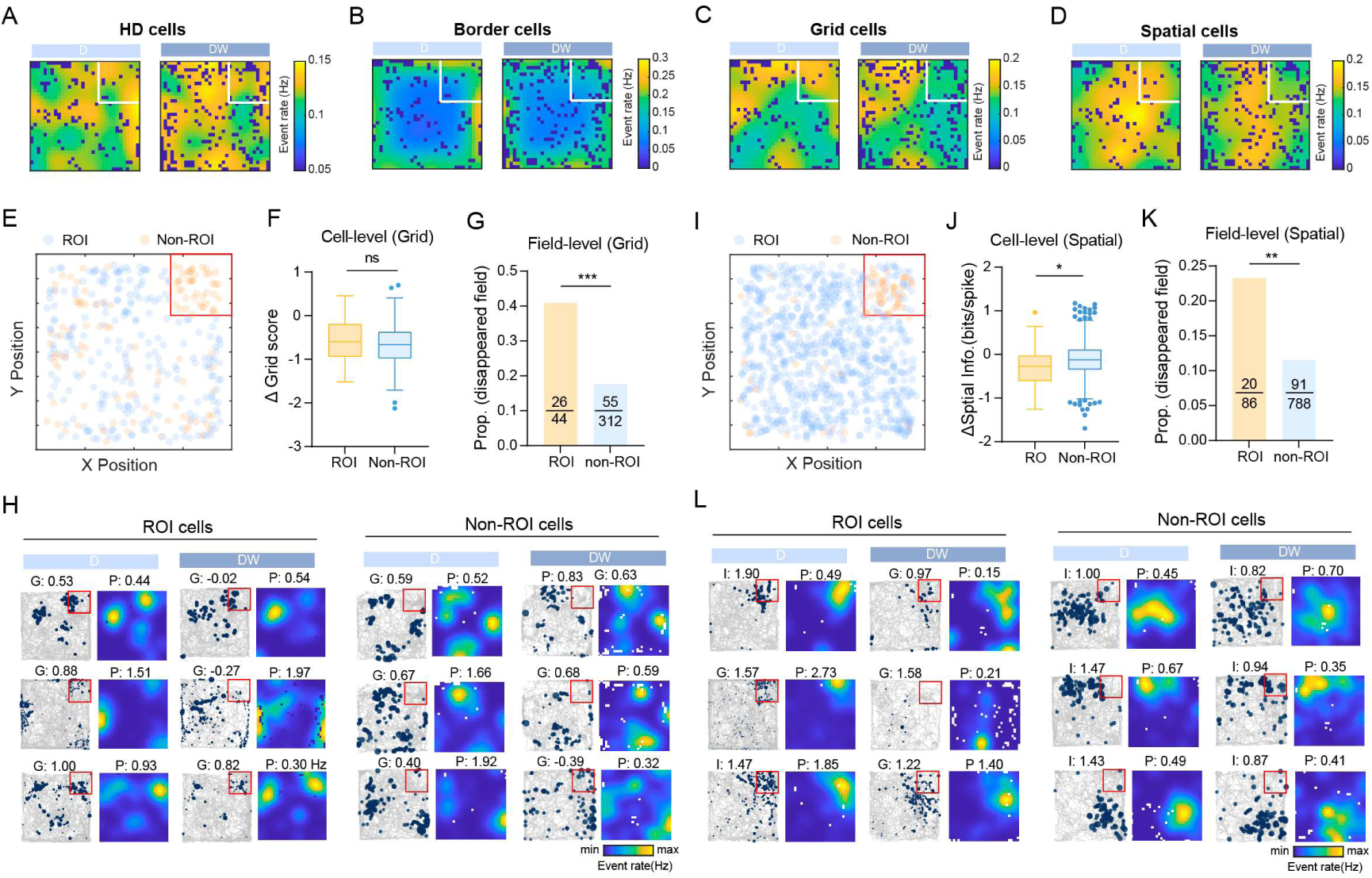
Fields anchored to dominant tactile floor cues are preferentially disrupted by whisker trimming. (A-D) Population-averaged rate maps for HD cells (A), border cells (B), grid cells (C), and spatial cells (D) in the Dark (D) and Dark, whisker-trimmed (DW) sessions. For each cell type, the mean event rate at each spatial bin was computed by averaging the rates of all cells of that type. The white square indicates the tactile cue region (sandpaper). Notably, grid cells and spatial cells show the most pronounced reduction in population mean rate within the tactile cue region after whisker trimming. (E) Spatial distribution of fields of grid cells, with field centroids located within the tactile cue region classified as region-of-interest (ROI) fields (blue) and all others classified as non-ROI fields (orange). The tactile cue region is outlined by the red box. (F) Cell-level comparison of changes in grid score (D/DW) between ROI grid cells and non-ROI grid cells. Grid cells were classified as ROI cells if at least one of their firing fields was located within the tactile cue region. No significant difference was observed between the two groups (Welch’s *t*-test, n = 44 ROI cells and n2 = 120 non-ROI cells, P = 0.94). (G) Field-level analysis showing the proportion of grid fields that disappeared after whisker trimming (defined as having no overlapping spatial bins between D and DW sessions). ROI grid fields exhibited a significantly higher disappearance probability than non-ROI fields (Chi-square test, P = 0.00034). (H) Representative examples of ROI and non-ROI grid cells across D and DW sessions. For each example, the animal’s trajectory (gray) and spatial rate maps are shown, with grid score (G) and peak rate (P) indicated, illustrating preferential degradation of grid fields anchored to the tactile cue region. (I) Spatial distribution of fields of spatial cells, with fields whose primary (largest) field centroid was located within the tactile cue region classified as ROI fields (blue), and others as non-ROI fields (orange). (J) Cell-level comparison of changes in spatial information (D/DW) between ROI and non-ROI spatial cells. ROI spatial cells show a significantly larger reduction in spatial information following whisker trimming (Welch’s t-test, n1 = 52 ROI cells and n2 = 561 non-ROI cells, P = 0.023). (K) Field-level comparison of the probability of field disappearance after whisker trimming for fields of spatial cells. Fields anchored to the tactile cue region exhibit a significantly higher disappearance probability than non-ROI fields (Chi-square test, P = 0.0020). (L) Representative examples of ROI and non-ROI spatial cells across D and DW sessions. Trajectories and spatial rate maps are shown, with spatial information (I) and peak rate (P) indicated, demonstrating that spatial cells anchored to the tactile cue region are more vulnerable to whisker trimming. **** P ≤ 0.0001; *** P ≤ 0.001; ** P ≤ 0.01; * P ≤ 0.05; ns, P > 0.05.

**Supplementary Figure 12.**
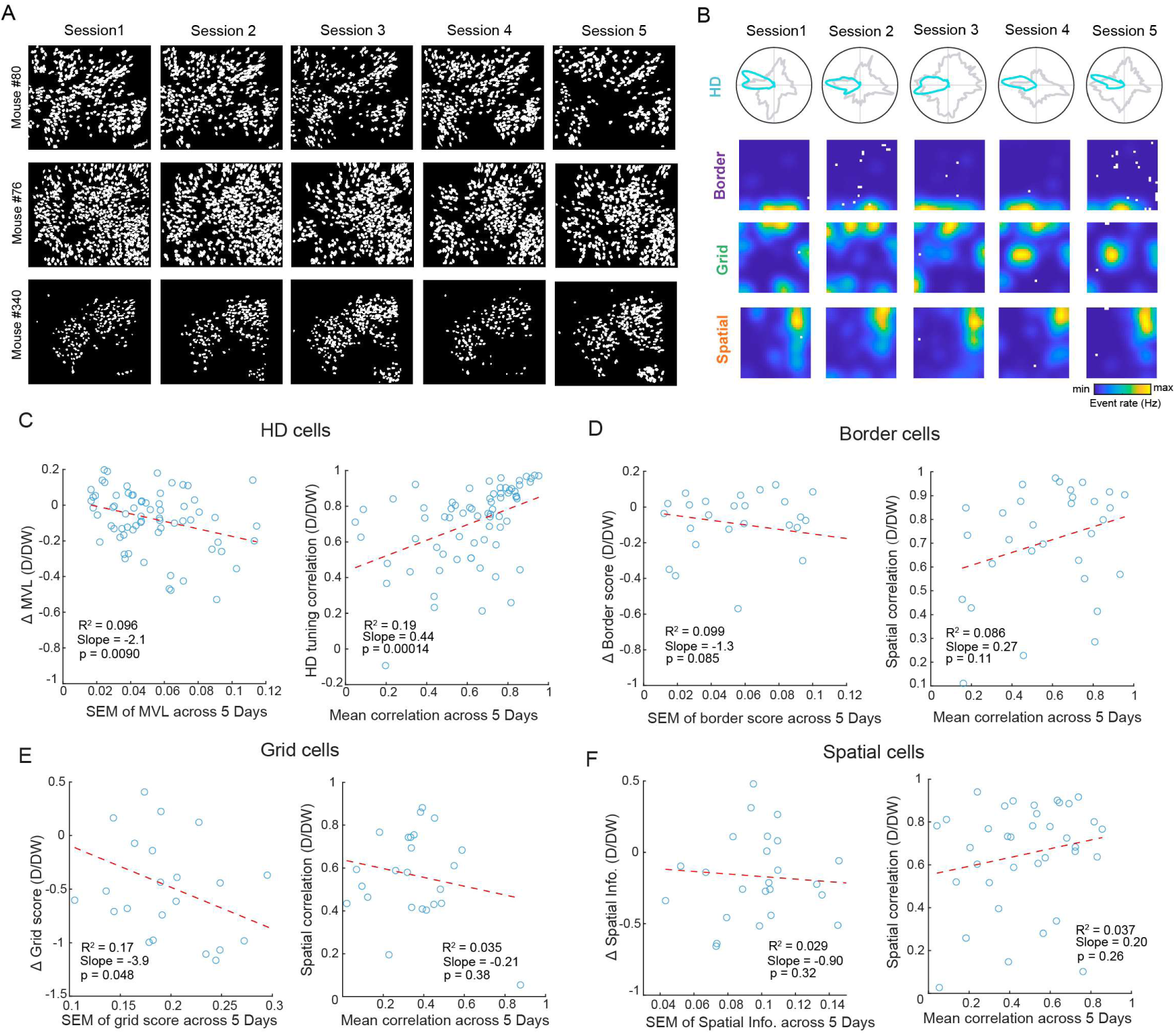
Cross-day stability modestly predicts sensory deprivation-induced changes in a subset of tuning metrics. (A) Cross-day spatial footprints after rigid transformation of three imaging mice exploring a dark, tactile cue-enriched environment (E2). (B) Representative examples of stable tuning curves across five recording sessions in darkness, demonstrating that spatially modulated cell types can form consistent representations based on tactile cues in the environment. From top to bottom: HD cells, border cells, grid cells and spatial cells. (C) HD cells. Left, relationship between whisker-trimming-induced change in tuning strength (ΔMVL, D/DW) and cross-day variability of tuning strength, quantified as the SEM of MVL across five imaging days. A significant negative association was observed (linear regression: R² = 0.096, slope = −2.1, P = 0.0090). Right, relationship between cross-day stability (mean tuning correlation across five days) and HD tuning correlation between D and DW sessions. A significant positive association was observed (linear regression: R² = 0.19, slope = 0.44, P = 0.00014). (D) Border cells. Left, relationship between whisker-trimming-induced change in border score (Δ border score, D/DW) and cross-day variability of border score (SEM of border score across five days). No significant association was detected (linear regression: R² = 0.099, slope = −1.3, P = 0.085). Right, relationship between cross-day stability (mean correlation across five days) and spatial correlation between D and DW sessions, showing no significant association (linear regression: R² = 0.086, slope = 0.27, P = 0.11). (E) Grid cells. Left, relationship between whisker-trimming-induced change in grid score (Δ grid score, D/DW) and cross-day variability of grid score (SEM of grid score across five days). A significant negative association was observed (linear regression: R² = 0.17, slope = −3.9, P = 0.048). Right, relationship between baseline cross-day stability (mean correlation across five days) and spatial correlation between D and DW sessions, showing no significant association (linear regression: R² = 0.035, slope = −0.21, P = 0.38). (F) Spatial cells. Left, relationship between whisker-trimming-induced change in spatial information (Δ Spatial Info., D/DW) and cross-day variability of spatial information (SEM of Spatial Info, across five days), showing no significant association (linear regression: R² = 0.029, slope = −0.90, P = 0.32). Right, relationship between cross-day stability (mean correlation across five days) and spatial correlation between D and DW sessions, showing no significant association (linear regression: R² = 0.037, slope = 0.20, P = 0.26). ****P ≤ 0.0001; ***P ≤ 0.001; **P ≤ 0.01; *P ≤ 0.05; ns, P > 0.05.

**Supplementary Figure 13.**
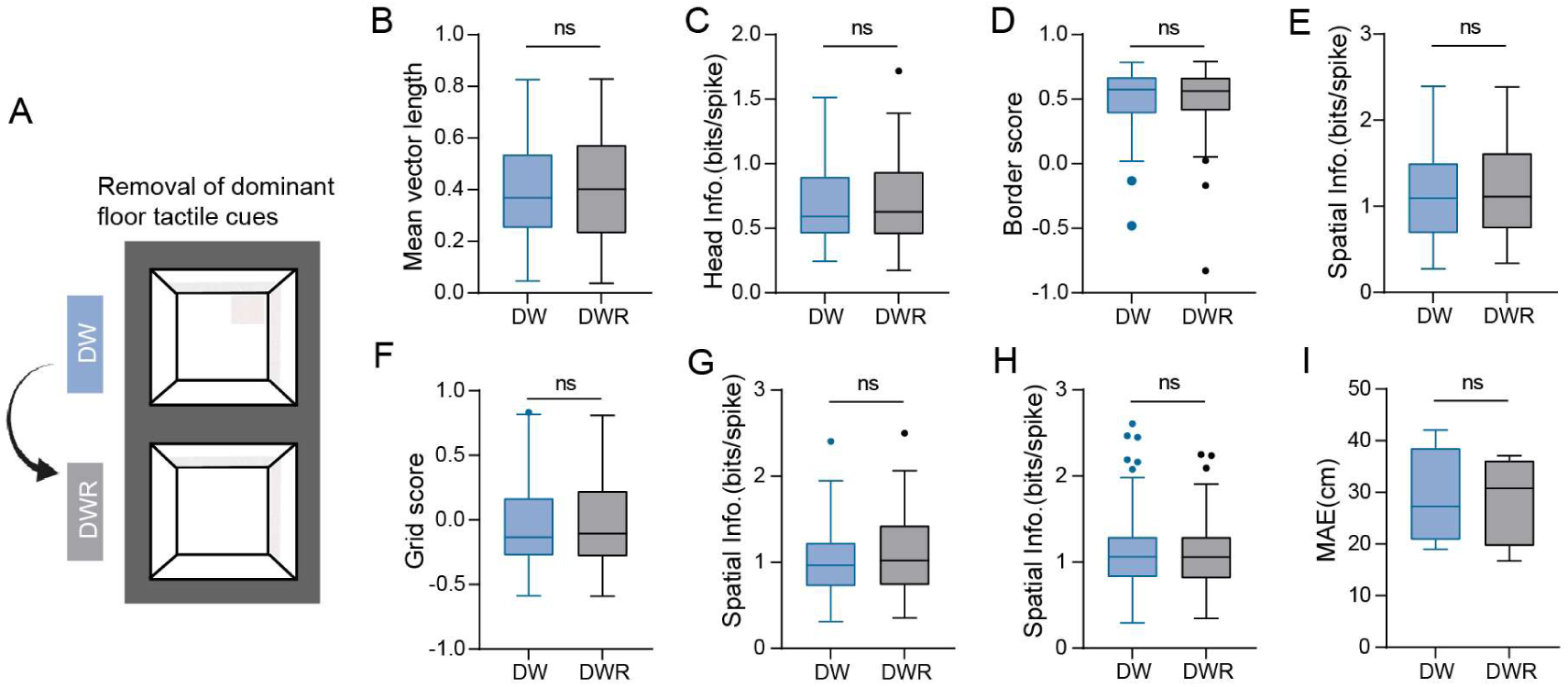
Removal of dominant floor tactile cues does not further degrade spatial representations after whisker trimming in darkness. (A) Paradigm schematic showing removal of the dominant floor tactile cue (white sandpaper patch) following whisker trimming in darkness (DW to DWR). (B) Box plot showing no significant change in MVL of HD cells in DWR compared with DW (Wilcoxon signed-rank test, n = 87, P = 0.053). (C) Box plot showing no significant change in head-direction information (bits/spike) of HD cells in DWR compared with DW (Wilcoxon signed-rank test, n = 87, P = 0.051). (D) Box plot showing no significant change in border score of border cells in DWR relative to DW (Wilcoxon signed-rank test, n = 83, P = 0.63). (E) Box plot showing no significant change in spatial information (bits/spike) of border cells in DWR compared with DW (Wilcoxon signed-rank test, n = 83, P = 0.48). (F) Box plot showing no significant change in grid score of grid cells in DWR compared with DW (Wilcoxon signed-rank test, n = 41, P = 0.95). (G) Box plot showing no significant change in spatial information (bits/spike) of grid cells in DWR compared with DW (Wilcoxon signed-rank test, n = 41, P = 0.14). (H) Box plot showing no significant change in spatial information (bits/spike) of spatial cells in DWR compared with DW (Wilcoxon signed-rank test, n = 189, P = 0.83). (I) Box plot showing no significant change in MAE of position decoding accuracy in DWR compared with DW using a decoder trained on neuronal activity from all spatially modulated cells in the baseline dark session (D) and tested on DW and DWR (Wilcoxon signed-rank test, n = 4 mice, P > 0.99). **** P ≤ 0.0001; *** P ≤ 0.001; ** P ≤ 0.01; * P ≤ 0.05; ns, P > 0.05.

**Supplementary Figure 14.**
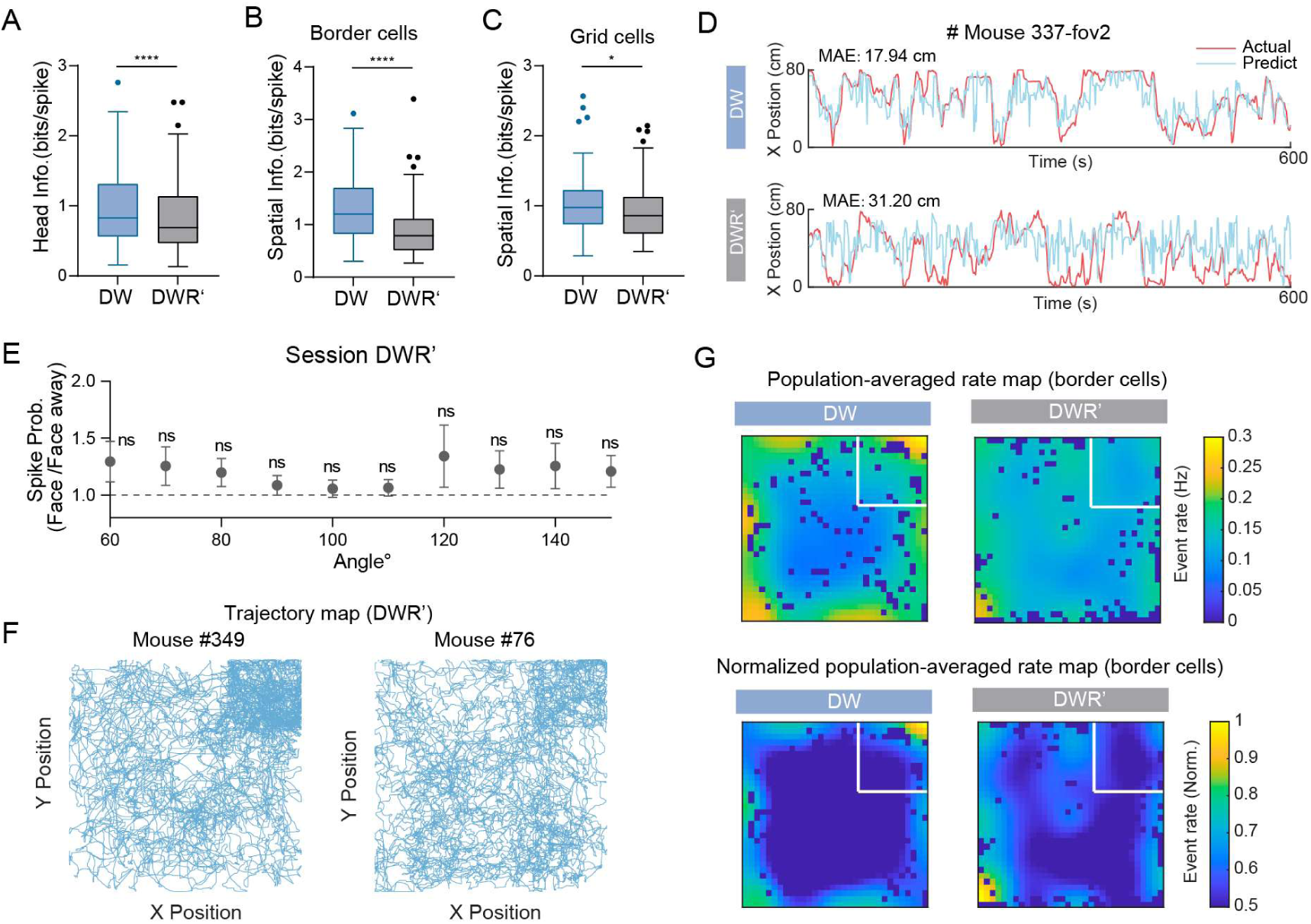
Removal of the arena wall further disrupts spatial representations after whisker trimming in darkness. (A) Box plot showing a significant decrease in head-direction (HD) information (bits/spike) in DWR′ compared with DW (Wilcoxon test, n = 276 cells, P = 2.6 × 10^-8^). (B) Box plot showing a significant decrease in spatial information (bits/spike) of border cells in DWR′ relative to DW (Wilcoxon test, n = 190 cells, P = 3.7 × 10^-15^). (C) Box plot showing a significant decrease in spatial information (bits/spike) of grid cells in DWR′ compared with DW (Wilcoxon test, n = 94 cells, P = 0.028). (D) Representative examples of position decoding in DW and DWR′ sessions using a decoder trained on neuronal activity from the baseline session (D). Actual position (red) and decoded position (blue) are shown, with the mean absolute error (MAE) indicated for each session. (E) After removal of the arena wall, the spike probability of border cells when the animal faced the wall relative to when it faced away from the wall (Face / Face away) was not significantly different from 1 across all tested angles (one-sample Wilcoxon signed-rank test vs. 1; 60°, P = 0.42; 70°, P = 0.27;70°, P = 0.27; 70°, P = 0.27; 80°, P = 0.17;90°, P = 0.09;100°, P = 0.07;110°, P = 0.20;120°, P = 0.32;130°, P = 0.37;140°, P = 0.14;150°, P = 0.17). (F) Representative trajectory maps from two example mice in the DWR ′ session. Behavioral occupancy was enriched in the upper-right corner of the open field, where a prominent tactile cue (white sandpaper patch) was placed on the floor. (G) Population-averaged firing-rate maps of border cells in the DW and DWR′ sessions. Top, population-averaged rate maps (event rate, Hz). Bottom, normalized population-averaged rate maps. The white square indicates the location and extent of the tactile cue (sandpaper) on the arena floor. **** P ≤ 0.0001; *** P ≤ 0.001; ** P ≤ 0.01; * P ≤ 0.05; ns, P > 0.05.

**Supplementary Figure 15.**
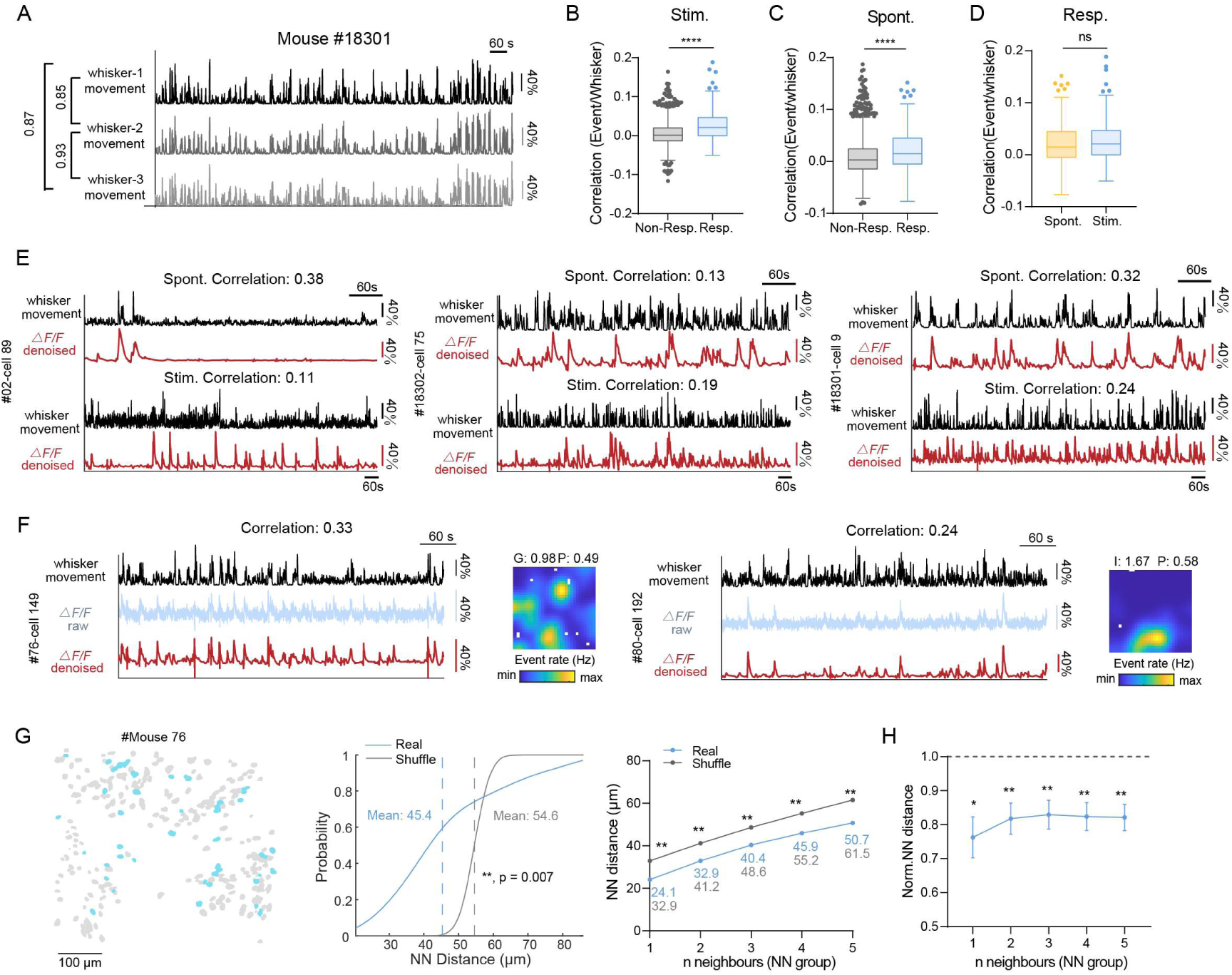
Certain MEC neurons respond to whisker movement, encompassing various spatially modulated cell types. (A) Representative traces of movements from three ipsilateral whiskers in an example mouse, illustrating highly consistent whisker motion patterns across whiskers. Whisker movement amplitude is expressed as normalized displacement (%). (B) Box plots showing that whisker-responsive cells (Resp.) exhibit significantly higher correlations between whisker movement and calcium event activity compared with whisker-non-responsive neurons (Non-Resp.) during the “stimulated-whisking” session (Stim.) (Welch’s *t*-test; n = 1,987 and n = 191, P = 7.8 × 10^-13^)。 (C) Same as (B), but during the “spontaneous-whisking” session (Spont.). Whisker-responsive cells again show significantly stronger correlation than non-responsive neurons (Welch’s t-test, n = 1981 and n = 190, P = 5.5 × 10^-6^). (D) Comparison of whisker-calcium correlation strength in whisker-responsive cells between the stimulated-whisking and spontaneous-whisking sessions, revealing no significant difference (Wilcoxon test, n = 190, P = 0.41). (E) Representative examples of “ whisker-responsive” cells, demonstrating consistent neuronal activity correlated with whisker movement in both "stimulated-whisking" and "spontaneous-whisking" sessions. Black traces indicate whisker movement; red traces show denoised ΔF/F calcium signals. (F) Example “whisker-responsive” cells with spatially-modulated properties. Right: grid cells, Left: Spatial cells. The black trace represents whisker movement over time, the blue trace shows raw ΔF/F, and the red trace shows denoised ΔF/F. The right panels display the corresponding spatial firing maps in the “free-exploration” session. (G) Spatial organization of whisker-responsive cells. Left: Anatomical distribution of whisker-responsive (blue) and non-responsive (gray) neurons within a representative FOV. The horizontal black line denotes a 100 μm scale bar. Middle: Cumulative distribution of nearest-neighbor (NN) distances among whisker-responsive cells (blue) compared with a shuffled distribution (gray)(Permutation test, P = 0.0070). Dashed lines indicate population means. NN distance was calculated using the three nearest neighbors. Right: Mean NN distance as a function of the number of nearest neighbors (NN group). Statistical significance for each NN group was assessed by permutation tests comparing real and shuffled data (1-5 neighbors; P = 0.0020, 0.0040, 0.0070,0.0040 and 0.0020). (H) Normalized mean NN distance of whisker-responsive cells across increasing NN groups, pooled across mice and normalized to shuffled controls. Only mice with more than 10 whisker-responsive cells were included. Blue dots indicate the mean real value across 8 mice, with error bars representing SEM. Statistical significance was assessed using a one-sample Wilcoxon signed-rank test against 1 (1-5 neighbors; P = 0.016, 0.0078, 0.0078,0.0078 and 0.0078). ****P ≤ 0.0001; ***P ≤ 0.001; **P ≤ 0.01; *P ≤ 0.05; ns, P > 0.05.

**Supplementary Figure 16.**
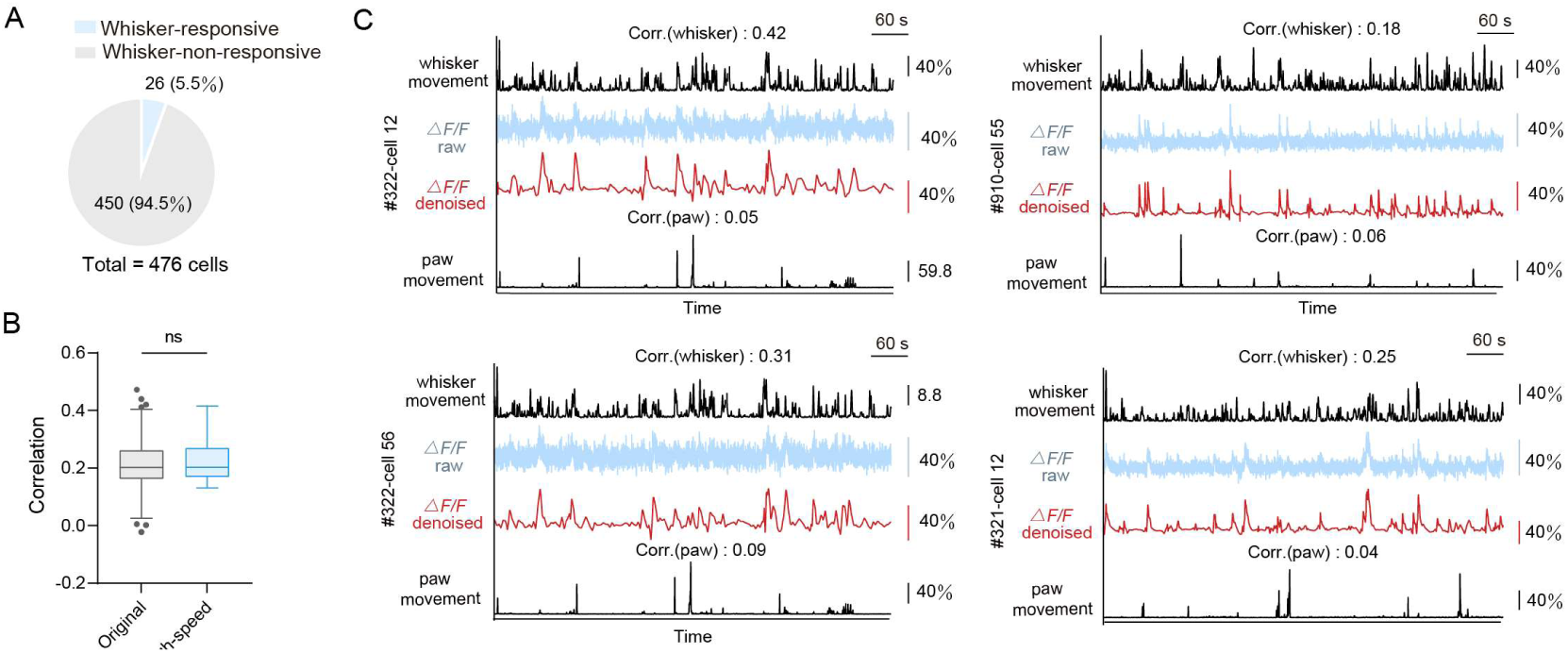
Whisker-responsive cells recorded using a dual-camera behavioral paradigm. (A) Proportion of MEC neurons classified as whisker-responsive versus whisker-non-responsive based on calcium activity recordings from 4 mice under high-speed infrared camera. (B) Box plot comparing the correlation between whisker movement and neuronal calcium activity across two experimental paradigms (original vs. high-speed recording). No significant difference was observed (Welch’s t-test, n = 26 and 191 whisker-responsive cells, P = 0.38). (C) Representative traces from individual whisker-responsive cells showing whisker movement (black), raw and denoised ΔF/F signals (blue and red), and paw movement (gray). Activity of some neurons show strong correlation with whisker movement but weak or no correlation with paw movement. ****P ≤ 0.0001; ***P ≤ 0.001; **P ≤ 0.01; *P ≤ 0.05; ns, P > 0.05.

## Notes

### Competing Interest Statement

The authors have declared no competing interest.

### Summary of Updates

We used another new tactile cue-rich open field to observe the changes in cells in the absence of visual cues in complete darkness, and found that in this condition, all spatially modulated cells could anchor to the tactile cues in the environment (Figure 4). Additionally, trimming the whiskers had a drastic impact on all cell types (Figure 5). Some cells were still able to maintain relatively stable firing, possibly due to the physical contact with the environmental walls (Figure 6). Furthermore, we raised the selection criteria for "whisker-responsive" cells and revised Figure 7 (formerly Figure 6).

